# Disturbed Flow Induces Reprogramming of Endothelial Cells to Immune-like and Foam Cells under Hypercholesterolemia during Atherogenesis

**DOI:** 10.1101/2025.03.06.641843

**Authors:** Christian Park, Kyung In Baek, Ruei-Chun Hung, Leandro Choi, Kiyoung Jeong, Paul Kim, Andrew Keunho Jahng, Jung Hyun Kim, Mostafa Meselhe, Ashwin Kannan, Chien-Ling Chou, Dong Won Kang, Eun Ju Song, Yerin Kim, Jay Aaron Bowman-Kirigin, Michael David Clark, Sander W. van der Laan, Gerard Pasterkamp, Nicolas Villa-Roel, Alyssa Panitch, Hanjoong Jo

## Abstract

**Background:** Atherosclerosis occurs preferentially in the arteries exposed to disturbed flow (d-flow), while the stable flow (s-flow) regions are protected even under hypercholesterolemic conditions. We recently showed that d-flow alone initiates flow-induced reprogramming of endothelial cells (FIRE), including the novel concept of partial endothelial-to-immune-cell-like transition (partial EndIT), but was not validated using a genetic lineage-tracing model. Here, we tested and validated the two-hit hypothesis that d-flow is an initial instigator of partial FIRE but requires hypercholesterolemia to induce a full-blown FIRE and atherosclerotic plaque development.

**Methods:** Mice were treated with adeno-associated virus expressing proprotein convertase subtilisin/kexin type 9 and a Western diet to induce hypercholesterolemia and/or partial carotid ligation (PCL) surgery to expose the left common carotid artery (LCA) to d-flow. Single-cell RNA sequencing (scRNA-seq) analysis was performed using cells obtained from the intima and leftover LCAs and the control right common carotid arteries at 2 and 4 weeks post-PCL. Comprehensive immunohistochemical staining was performed on EC-specific confetti mice treated with PCL and hypercholesterolemic conditions at 4 weeks post-PCL to validate endothelial reprogramming.

**Results:** Atherosclerotic plaques developed by d-flow under hypercholesterolemia at 2 and 4 weeks post-PCL, but not by d-flow or hypercholesterolemia alone, as expected. The scRNA-seq results of 98,553 single cells from 95 mice revealed 25 cell clusters; 5 EC, 3 vascular smooth muscle cell (SMC), 5 macrophage (MΦ), and additional fibroblast, T cell, natural killer cell, dendritic cell, neutrophil, and B cell clusters. Our scRNA-seq analyses showed that d-flow under hypercholesterolemia transitioned healthy ECs to full immune-like (EndIT) and, more surprisingly, foam cells (EndFT), in addition to inflammatory and mesenchymal cells (EndMT). Further, EC-derived foam cells shared remarkably similar transcriptomic profiles with foam cells derived from SMCs and MΦs. Comprehensive lineage-tracing studies using immunohistochemical staining of canonical protein and lipid markers in the EC-specific confetti mice clearly demonstrated direct evidence supporting the novel FIRE hypothesis, including EndIT and EndFT, when d-flow was combined with hypercholesterolemia. Further, reanalysis of the publicly available human carotid plaque scRNA-seq and Perturb-seq datasets supported the FIRE hypothesis and a potential mechanistic link between the genes and FIRE.

**Conclusion:** We provide evidence supporting the two-hit hypothesis: ECs in d-flow regions, such as the branching points, are partially reprogrammed, while hypercholesterolemia alone has minimal endothelial reprogramming effects. Under hypercholesterolemia, d-flow fully reprograms arterial ECs, including the novel EndIT and EndFT, in addition to inflammation and EndMT, during atherogenesis. This single-cell atlas provides a crucial roadmap for developing novel mechanistic understanding and therapeutics targeting flow-sensitive genes, proteins, and pathways of atherosclerosis.

## Introduction

Atherosclerosis, a chronic inflammatory disease resulting in accumulation of plaque in the arterial wall and lumen narrowing, is the major underlying cause of myocardial infarction, ischemic stroke, and peripheral arterial disease^1–3^. Despite the widespread and successful use of therapeutics, including statins that effectively lower one of the most critical proatherogenic risk factors, blood cholesterol level, atherosclerotic disease continues to be a leading cause of death worldwide^4^. It highlights the dire need to develop additional therapeutics to address the residual non-lipid targets. For example, CANTOS trial, using the IL-1β inhibitor canakinumab, demonstrated potential promise of targeting inflammatory factors as anti-atherogenic therapy, although it was not approved by the FDA due to a concern with fatal infections^5^. Since atherosclerosis preferentially occurs in flow-disturbed regions such as branching points, we and others have been studying flow-sensitive factors as potential therapeutic targets^6–12^. Here, we report a single cell RNA-sequencing (scRNA-seq) study to identify flow-sensitive genes and pathways that may be responsible for atherosclerosis development in response to disturbed flow (d-flow) and hypercholesterolemic condition in the mouse carotid arteries.

Even though hypercholesterolemia, diabetes, and hypertension are major systemic risk factors, atherosclerotic plaques preferentially occur focally involving curved, branched, and bifurcated regions of arteries exposed to d-flow^4,7,8,10^. In contrast, straight arterial regions exposed to stable flow (s-flow) tend to be protected from atherosclerosis. The mechanisms underlying the different effects of d-flow and s-flow on atherosclerosis involve endothelial gene expression and function are still unclear^7,9–11^. To address the mechanisms, we developed the mouse partial carotid ligation (PCL) model, by ligating 3 of 4 downstream branches of the left common carotid artery (LCA) to induce d-flow while using the contralateral right common carotid artery (RCA) exposed to s-flow as a control within the same mouse^13^. With the PCL model, we directly demonstrated that d-flow rapidly induces robust atherosclerotic plaque development within 2-3 weeks only under hypercholesterolemic condition. The study also showed that d-flow alone in the absence of hypercholesterolemia or hypercholesterolemia under s-flow failed to develop atherosclerosis, demonstrating that atherosclerosis development is promoted by the interaction between the systemic hypercholesterolemia and focal d-flow. This is consistent with the preferential atherosclerosis development in the branching points exposed to d-flow in hypercholesterolemic patients^6,12^.

D-flow and s-flow differently regulate endothelial function through mechanosensors^14–18^ and gene expression, leading to initiation and prevention of atherosclerosis, respectively^9,19–26^. To explore the mechanisms, we previously carried out a scRNA-seq and single cell assay for transposase-accessible chromatin sequencing (scATAC-seq) study in response to d-flow alone under normal cholesterol conditions to avoid the complex mechanisms under the combination of d-flow and hypercholesterolemia^27^. In the previous study, we also collected single cells from the intima of the LCAs and RCAs to obtain a sufficient number of ECs for an in-depth examination of the effects of d-flow alone on endothelial cells (ECs). In the study, C57BL/6 mice underwent PCL surgery and fed a chow diet, followed by single cell isolation from the LCAs and RCAs 2 days and 2 weeks post-PCL in a time- and flow-dependent manner^27^. The study showed that ECs are highly plastic and heterogeneous. More importantly, healthy ECs transition to proatherogenic phenotypes, including inflammation, endothelial-to-mesenchymal transition (EndMT), and the novel finding of endothelial-to-immune cell-like transition (EndIT), which we collectively named as flow-induced reprogramming of ECs (FIRE)^9,27^.

Interestingly, our pseudotime trajectory analysis showed that many ECs underwent full EndMT. However, only a small number of ECs underwent EndIT and did not fully transition into immune cells in response to d-flow alone, which we refer to as partial EndIT. These prior results showed that d-flow alone initiates partial FIRE, involving inflammation, EndMT, and partial EndIT. However, the partial EndIT suggested by scRNA-seq required validation by methods such as genetic lineage-tracing, which remained to be done. Since d-flow alone could not induce atherosclerotic plaques in the absence of systemic risk factors such as hypercholesterolemia^13,28^, we hypothesized that a combination of d-flow and hypercholesterolemia would induce atherosclerotic plaques by inducing robust FIRE, especially EndIT in addition to inflammation and EndMT.

Here, we tested the hypothesis by performing an extensive scRNA-seq study to compare the effects of d-flow alone, hypercholesterolemia alone, and d-flow under hypercholesterolemia on ECs as well as all other arterial cell types in the atherosclerotic plaques. Our scRNA-seq results demonstrate that d-flow under hypercholesterolemia induces atherosclerotic plaques and show a robust EndIT in addition to inflammation and EndMT. Most surprisingly, we found evidence that ECs underwent endothelial-to-foam cell transition (EndFT).

We further validated FIRE revealed by our scRNA-seq study in a lineage-tracing mouse model using extensive immunostaining studies for markers of endothelial inflammation, EndMT, EndIT, and EndFT in EC-specific confetti (EC-Confetti) mice. Our results of scRNA-seq and the lineage tracing model studies demonstrate that d-flow and hypercholesterolemia induce a robust FIRE, especially the novel EndIT and EndFT in addition to the expected endothelial inflammation and EndMT, during atherogenesis. These are by far the most comprehensive scRNA-seq atlas of mouse atherosclerotic plaques, especially for ECs, providing invaluable resources for the field.

## Methods

### Transparency and Openness

The authors will make the data, methods used in the analysis, and materials used to conduct the research available to any researcher for purposes of reproducing the results or replicating the procedure.

### Mouse scRNA-seq study

Mouse atherosclerosis scRNA-seq study was approved (PROTO202100052) by the Emory University institutional animal care and use committee and conducted in accordance with the federal guidelines and regulations. A total of 75 male C57BL/6 mice (Jackson Lab, 9 to 10-week-old) were used in this scRNA-seq study performed in 3 independent experiments (Table S2). As described previously, hypercholesterolemia was induced through a combination of AAV8-PCSK9 administration (1×10^11^ viral genome) (Vector Biolabs #AAV8-D377Y-mPCSK9) via tail-vein injection for low-density lipoprotein receptor (LDLR) knockout and feeding with a high-fat Western diet a week before the PCL surgery^29^. PCL surgery was performed as previously described through ligation of 3 of the 4 caudal branches (left external carotid, left internal carotid, and left occipital arteries) of the LCAs in mice that were anesthetized, with the RCAs left intact as the contralateral control^13^. Ultrasound imaging was performed to confirm the exposure of the LCA to d-flow, as described previously^13^. Two or 4 weeks post-PCL surgery, mice were sacrificed for scRNA-seq experiments by CO_2_ asphyxiation.

### Isolation of arterial cells from the carotid arteries and library preparation for scRNA-seq

Single cell preparation and sequencing were conducted as we previously reported^27^. Briefly, cell dissociation buffer was composed of 500 U/mL of collagenase type I (EMD Millipore #SCR103), 500 U/mL of collagenase type II (MP Biomedical #0.2100502.5), 150 U/mL of collagenase type XI (Sigma-Aldrich #C7657), 60 U/mL of hyaluronase type I-S (Sigma-Aldrich #H3506), and 60 U/mL of DNASE I (Zymo #E1011) in HBSS (Cytiva #SH30031) and filtered through a 0.45 μm syringe filter (Celltreat #229753).

Upon euthanization, blood was collected from the inferior vena cava with heparin-coated 25G needle (Air-Tite #N2558) to measure the plasma cholesterol level. A lobe of liver was also collected to validate LDLR knockout by Western blot. Then, LCAs and RCAs were cleaned and perfused with saline solution before injecting the dissociation buffer. The ends of the carotid arteries were ligated using sutures and the arteries were dissected out. After incubation in the 35 mm dishes containing PBS at 37°C for 45 min, sutures were removed from the ends. The carotid arteries were luminally flushed using the dissociation buffer into 1.5 mL Eppendorf tubes containing fetal bovine serum (FBS) for neutralization of enzymatic activity and then kept on ice.

Leftover LCAs and RCAs were placed into new tubes containing dissociation buffer and minced using microscissors. The minced tissues were additionally incubated at 37°C for 45 min, after which the entire cell solution was moved to 1.5 mL Eppendorf tubes containing FBS on ice. Both luminal and leftover digestion samples were filtered through a 70 μm cell strainer to remove large pieces of extracellular matrix, followed by a series of washes with 2% bovine serum albumin (BSA) in PBS solution and centrifugation at 1000 g for 5 min. RBC lysis was performed via incubation of the cell solutions in the Hybri-Max buffer (Sigma-Aldrich #R7757) at RT for 5 min, followed by incubation in accutase (Sigma-Aldrich #A6964) at 37°C for 5 min to prepare single cell solutions. The single cells were finally resuspended in 2% BSA in PBS solution and immediately encapsulated at the Emory Integrated Genomics Core (EIGC) using the 10X Genomics Chromium Next GEM Single Cell 3’ Kit v3.1 and the Chromium X device. The cDNA libraries were constructed, followed by sequencing on Illumina NovaSeq instrument to a minimum depth of 25,000 reads per cell.

### scRNA-seq data preprocessing and batch effect correction

scRNA-seq data files were preprocessed and aligned to the mouse reference genome (mm10) with 10X Genomics CellRanger software. CellRanger barcode ranked plots are provided in Figure S1. Downstream analyses of the scRNA-seq data were performed using the *Seurat* v5 R package^30^. Quality control measures were implemented to filter out the cells expressing <200 or >7,600 genes and those with >10% counts aligned to mitochondrial genes to remove non-singlets or damaged cells, respectively. Normalization, scaling, clustering, and visualization using UMAP were then performed for each of the scRNA-seq datasets from our previous study^27^ and the newly conducted studies here composed of 3 independent libraries (Table S2).

Batch effect correction arising from combining multiple scRNA-seq datasets was performed using the *Seurat* integration function, where the datasets were merged and normalized, followed by integration using reciprocal principal component analysis (RPCA) in *Seurat*. The 4 independent libraries were considered as batches (Table S2). To quantify the changes upon batch effect correction, 2 types of benchmarking metrics were used. Batch mixing metrics quantify the degree of how well-mixed the datasets are in the projected space, while cell type conservation metrics quantify how well-conserved the biological variations of cells are upon mixing^31,32^. Batch mixing metrics used are iLISI: integration local inverse Simpson’s index (LISI), ASW_batch: batch average silhouette width (ASW), and kBET: k-nearest neighbor batch effect test. Cell type conservation metrics used are ARI: adjusted rand index, cLISI: cell type LISI, and ASW_celltype: cell type ASW. ARI value ranges from 0 to 1, with higher score indicating better cell type conservation. Value of LISI ranges from 1 to the number of batches, which is 4 in this study. iLISI score closer to 4 indicates enhanced batch mixing while cLISI score closer to 1 indicates higher cell type conservation. ASW value ranges from −1 to 1, where lower ASW_batch score suggests better batch mixing, while higher ASW_celltype score suggests enhanced cell type conservation. kBET rejection rates range from 0 to 1, with lower value indicative of improved batch mixing. Batch mixing metrics quantified using all cells showed better batch mixing as indicated by higher iLISI and lower kBET using sample size of 10 and 20% after integration, while those using only the ECs also showed better batch mixing as shown by higher iLISI and lower ASW_batch and kBET with a sample of size of 10% after integration (Table S2). Similarly, cell type conservation assessed using all cells showed higher cell type purity as indicated by higher ARI after integration. Due to the presence of a single cell type, cell type conservation metrics were unable to be measured using only the ECs (indicated as N/A). Furthermore, ASW metrics using all cells were unable to be quantified with the publicly available packages. The *Seurat* RPCA method was selected for batch effect correction of our datasets since it provided better improvements in the overall benchmarking metrics above compared to Seurat canonical correlation analysis (CCA), Harmony, and LIGER integration methods (data not shown).

ARI was measured using the *fossil* R package, while iLISI and cLISI were quantified using the *lisi* package. ASW_batch and ASW_celltype were quantified using the *cluster* package, while kBET was measured on 10-20% sample size using the *kBET* package.

### scRNA-seq cell clustering analysis

Following integration, scaling, clustering, and visualization were performed on the integrated scRNA-seq object in *Seurat*. Manual annotation of cell clusters was performed on the list of enriched genes generated from differential gene expression (DGE) analysis of each cluster against the others and sorted by avglog2FC. Canonical cell markers were used for the overall cell type annotation. More detailed clustering analysis of cell types with multiple clusters was performed by subsetting these cells, followed by DGE analysis of every cluster for each cell type independently.

Gene Ontology (GO) biological process (BP) terms were generated by uploading top 100 differentially expressed genes to the Gene Ontology consortium website (https://geneontology.org/) and sorted by p-value. Heatmap was generated using *Seurat* to display the top 20 most highly enriched genes for each cluster. Cell-cell communication analyses were performed using the *CellChat* R package following the developer’s instructions^33^.

### scRNA-seq trajectory analysis

Diffusion map analysis, an unsupervised and unbiased form of trajectory analysis, was performed on the subset of ECs, SMCs, and MΦs and projected into both 2D and 3D using the *destiny* R package following the developer’s instructions^34^. Monocle pseudotime trajectory analysis was also carried out on these 3 cell types using the *Monocle 3* R package following the developer’s instructions^35^. EC1, SMC1, MΦ1, and MΦ5 were set as roots manually. RNA velocity analysis was performed by generating loom files for spliced and unspliced matrices using the *Velocyto* Python package^36^ and merging them to the adata file created from annotated *Seurat* scRNA-seq object using the *scVelo* Python package^37^. Stochastic mode was implemented for RNA velocity. Maps showing velocity length and velocity confidence were generated using *scVelo*.

### Reference map analysis to human carotid scRNA-seq data

The publicly available scRNA-seq data of 4,811 single cells isolated, sequenced, and cluster annotated from 44 human carotid endarterectomy samples was used as a reference for reference map analysis^38,39^. Our mouse atherosclerosis scRNA-seq data genes were first converted to human gene orthologs using the *biomaRt* R package. Reference map analysis was performed using *SingleR* package^40^, which effectively predicted the cell annotations for the human carotid scRNA-seq object using the transcriptomic profiles of cells and their annotations from our mouse scRNA-seq data. The cell number for each predicted cell cluster was quantified based on the predicted cell annotations from reference map analysis.

### Reanalysis of CRISPRi-Perturb-seq data

The publicly available CRISPRi-Perturb-seq data knocking down 2,285 genes associated with 306 coronary artery disease genome-wide association study (CAD-GWAS) signals at a single cell resolution in human ECs was reanalyzed^41^. Perturbation of 2,285 genes significantly altered expressions of 17,849 genes (P value < 0.001). The lists of perturbed genes and genes altered in response to Perturb-seq were compared against 1,291 flow-sensitive genes that were significantly and differentially expressed (average log_2_fold change > 1) in the 5 EC clusters. Again, the mouse genes were converted to human gene orthologs using the *biomarRt* package. These reanalyses enabled the identification of flow-sensitive genes that were perturbed and flow-sensitive genes that were altered in response to perturbations. Furthermore, an alluvial plot linking flow-sensitive genes that altered expressions of FIRE markers used to annotate EC clusters was generated using the *ggalluvial* package.

### Lineage Tracing Study using EC-Confetti Mice

The lineage tracing study was approved (PROTO202100052) by the Emory University institutional animal care and use committee and conducted in accordance with federal guidelines and regulations. We established an EC-specific confetti mouse model (EC-Confetti) as previously described^42^ by crossing loxP-flanked (floxed) cytosolic RFP, cytosolic YFP, nuclear GFP, and membrane CFP (Confetti^FL/WT^) mice (Jackson Lab) with tamoxifen-inducible, endothelial-specific Cdh5-iCreER^T2^ mice provided by Dr. Ralf Adams. EC-Confetti mice were genotyped and screened by PCR and DNA sequencing and used for lineage tracing study^42^ to validate FIRE.

A total of 21 (6 Male/15 Female) 4-6-week-old EC-Confetti mice were injected intraperitoneally with tamoxifen dissolved in corn oil (75 mg/kg body weight) over the course of 1 week (series of 5 injections)^42^. Following a 2 week resting period, AAV-PCSK9 injection, Western diet feeding, and PCL surgery were carried out and sacrificed at 4 weeks post-PCL, as described for the scRNA-seq study. The carotid arteries and aortic arches were isolated immediately after mouse euthanization. LCAs and RCAs were post-fixed using 4% PFA (Santa Cruz Biotechnology #sc-281692). As we reported previously, a section of the RCA and two segments of the LCA, as well as the aortic arch, were longitudinally cryosectioned for imaging and subsequent analyses^19^. Immunohistochemical staining against multiple canonical markers of FIRE was performed as previously described^19^. FIRE protein markers were endothelial inflammation (Vcam1 and Icam1), EndMT (Snai1, Acta2, and Cnn1), immune cells (Cd68, C1qa, C1qb, and Lyz2), and foam cells (Spp1, Lgals3, and Trem2).

The specificity of each primary antibody used in this study (Table S1) was validated by conducting dilution curve studies (data not shown) and isotype and secondary antibody alone controls (Figure S15). After blocking at room temperature for 1 hour, sections were incubated at 4°C overnight with primary antibodies diluted in PBS supplemented with 0.1% Triton X100 and 5% BSA. AlexaFluor 568- (Thermo Fisher Scientific #A20184) or 647-conjugated secondary antibody (Thermo Fisher Scientific #A20186) was used at room temperature for 2 hours. For BODIPY staining, the cryosectioned LCAs and RCAs were stained with 2 µM BODIPY493/503 (Thermo Fisher Scientific #D3922) for 1 hour at 37°C as previously described^19^. The stained sections were then Dapi-mounted for fluorescence imaging.

Confetti (nuclear GFP, cytosolic YFP, and cytosolic RFP) and FIRE marker expressions were imaged by using inverted fluorescence microscope (Keyence #BZ-X800, Japan) as previously described^22^. Membrane CFP was not detectable as previously reported^42^. Following blind deconvolution and dehazing of out-of-focus illumination, image superimposition and automated image stitching were performed by using custom-coded MATLAB algorithms (MathWorks, MA), ImageJ, and Fiji (NIH, MD). To quantify the level of confetti induction, luminal Confetti^+^ ECs within the LCAs and RCAs (N = 69 slides, 21 mice) were manually defined and segmented as described^43^. To determine the proportion of confetti^+^ ECs undergoing FIRE, multi-level image thresholding was performed via combined use of custom-developed MATLAB algorithms and ImageJ^43^.

Confetti^+^/FIRE^+^ region was defined via binarized masks, while non-overlap or unmasked region was derived computationally to evaluate Confetti^+^/FIRE^-^ region. Intensity histogram of Confetti^+^/FIRE^+^ region in the LCAs and RCAs was normalized for quantification. Clonal expansion of ECs was assessed by color distribution of each fluorescent reporter protein as previously described^42^. Co-immunofluorescence staining was performed using appropriate combinations of primary antibodies with compatible host species (Table S1).

### Validation of LDL receptor knockdown by AAV-PCSK9 and hypercholesterolemia

To validate knockout of LDLR, mouse liver tissues were homogenized using BeadBug™ 6 microtube homogenizer (#31-216) in tubes containing RIPA buffer (Boston BioProducts #BP-115) with complete Mini protease inhibitor cocktail (Roche Diagnostics #11836153001) to extract proteins. The total protein concentrations were determined using BCA assay (Thermo Fisher Scientific #23225). For SDS-PAGE analysis, 10% gel was used, and the electrophoresis was run at 110 V for 1.5 hrs. The proteins were then transferred overnight at 25 V and 4°C onto the PVDF membranes (Bio-Rad #1620174). For protein detection, the PVDF membranes were incubated overnight at 4°C with primary antibodies: anti-LDLR rabbit monoclonal antibody (1:1000; Abcam #AB286156) and anti-β-actin mouse monoclonal antibody (1:1000; Sigma-Aldrich #A5316), which served as a loading control for the liver protein samples. After a series of washes with TBS-T buffer, the membranes were incubated at room temperature (RT) for 1 hr with the corresponding secondary antibodies: goat anti-mouse IgG horseradish peroxidase (HRP) (Cayman Chemical #10004302) and goat anti-rabbit IgG HRP (Cayman Chemical #10004301). Protein bands were visualized using the Immobilon Western chemiluminescent HRP substrate (Millipore #WBKLS0500) and the iBright FL1000 imaging system, which were quantified using ImageJ software and normalized based on the average expression level at control groups (Ctrl_2wk and Ctrl_4wk).

As reported previously, mouse plasma lipid levels were analyzed by the Emory Biomarker Core Laboratory using Beckman CX7 biochemical analyzer^19^.

### Statistical analyses

Statistical analyses were conducted using GraphPad Prism version 10.0.0 software. Sample number N is included in each figure caption, and all data are presented as mean ± standard error of the mean (SEM). All datasets were analyzed for normality using the Shapiro-Wilk test, and for equal variance using the F test for datasets with 2 groups, or the Brown-Forsythe test for groups of 3 or more. Comparisons between 2 groups were conducted using either 2-tailed unpaired Student t test for normally distributed data, or 2-tailed unpaired Mann-Whitney test for non-normally distributed data. For normally distributed data with unequal variances, the Student t test with Welch’s correction was used. Comparisons between 3 or more groups were conducted using the 1-way ANOVA for normally distributed data or the Kruskal-Wallis test for non-normally distributed data. For normally distributed data with unequal variances, the Brown-Forsythe ANOVA test was used. The Tukey, Dunnett and Dunn multiple-comparisons tests were used for post hoc pairwise comparisons following ANOVA. The Tukey test was used when all groups in the analysis were compared. The Dunnett test was used when all groups were compared with a single control group. The Dunn test was used to compare all groups against each other. *P*<0.05 was considered significant for all statistical tests that were performed.

## Results

### D-flow induces drastic shifts in arterial cell populations of mouse carotid arteries during atherogenesis

To define the transcriptomic landscape of arterial cells at a single cell resolution during atherogenesis in response to d-flow under hypercholesterolemia, we carried out a scRNA-seq study using the common carotid arteries. For this study, we used 75 male C57BL/6 mice treated with 1) control (s-flow and chow diet), 2) d-flow alone (chow diet), 3) hypercholesterolemia alone under s-flow (s-flow), and 4) d-flow under hypercholesterolemic conditions at 2 and 4 weeks after PCL surgery (Figure 1A, S2, Table S2).

**Figure 1.**
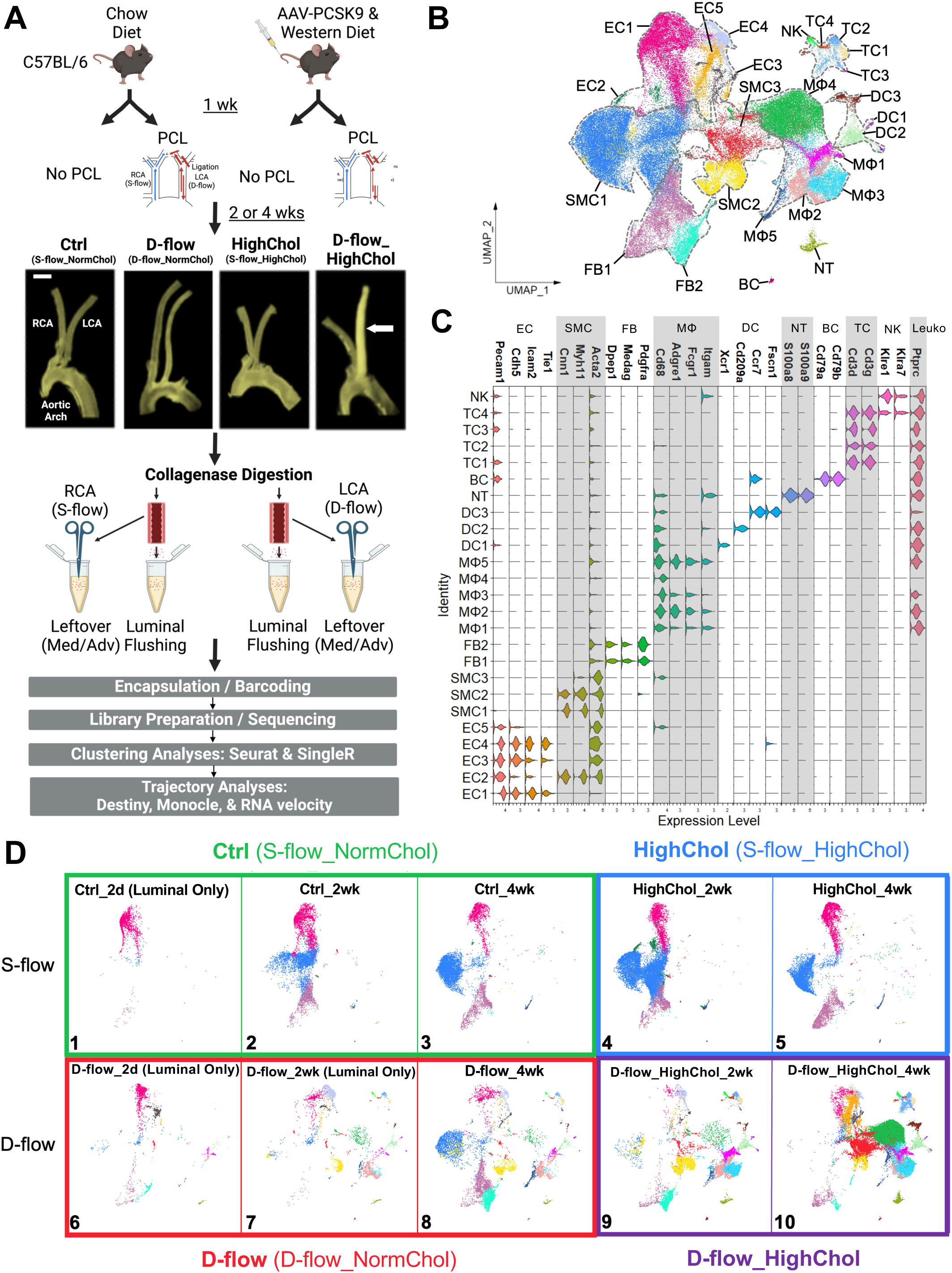
scRNA-seq data clustering analysis of mouse carotid artery cells exposed to d-flow and/or hypercholesterolemia during atherogenesis. **A**. C57BL/6 mice were treated with or without AAV-PCSK9 injection and Western diet for 2 or 4 weeks with or without PCL surgery. Representative macroscopic images of mouse carotid arteries and aortic arch shown for 4 weeks post-PCL time points. Atherosclerotic plaque development occurred only in the LCAs of the d-flow and hypercholesterolemia group (white arrow). Scale bar: 1 mm. **B**. UMAP plot of 98,553 cells from the scRNA-seq data of Ctrl (s-flow, normal cholesterol), D-flow (d-flow, normal cholesterol), HighChol (s-flow, hypercholesterolemia), and D-flow_HighChol (d-flow, hypercholesterolemia) groups at 2 and 4 weeks post-PCL mice reveals 25 unique cell clusters. Major cell populations include endothelial cells (ECs), vascular smooth muscle cells (SMCs), fibroblasts (FBs), macrophages (MΦs), dendritic cells (DCs), neutrophils (NTs), B cells (BCs), T cells (TCs), and natural killer cells (NKs). Leukocytes include MΦs, DCs, NTs, BCs, TCs, and NKs. **C**. Stacked violin plot shows expression levels of canonical marker genes used to annotate each cell cluster. **D**. UMAP plot of each experimental condition is shown across time (2 days, 2 weeks and 4 weeks). S-flow (top): Ctrl (left, boxed in green) and HighChol (right, blue) and D-flow (bottom): D-flow (left, red) and D-flow_HighChol (right, purple) are shown. N= 5-20 mice for each condition. Note that Ctrl_2d, D-flow_2d, and D-flow_2wk conditions contain only luminally collected cells from the previous work.

Hypercholesterolemia was induced by a single AAV8-PCSK9 injection and feeding with Western diet, as we previously published^29^. D-flow was induced in the LCA by PCL, while the contralateral RCA was continued to be exposed to s-flow. LCAs and RCAs of mice without PCL surgery were exposed to s-flow. As expected, LCAs developed atherosclerotic plaques only in the d-flow under hypercholesterolemia group at 2 and 4 weeks post-PCL (Figure 1A, S2). AAV-PCSK9 injection effectively knocked out the LDL receptors (LDLR) in mouse liver as validated by Western blots (Figure S3A-B), and hypercholesterolemia (plasma total cholesterol > 400 mg/dl) was confirmed for hypercholesterolemia alone and d-flow under hypercholesterolemia groups (Figure S3C-G).

Single cells were first prepared from luminal digestion and flushing of the LCAs and RCAs to obtain the maximum number of ECs (Figure 1A). The remaining arteries were further digested to obtain single cells from the media and adventitia. Single cells from the lumen and leftover were then encapsulated, barcoded, library prepared, and sequenced separately. The scRNA-seq datasets from both the lumen and the leftover were pooled together during downstream analysis to enrich the EC numbers. Consistent with our previous publication^27^, quality control measures were performed on the scRNA-seq data in *Seurat*^30^ by selecting the single cells that expressed 200-7,600 genes per cell and <10% mitochondrial genes to exclude the non-singlets or damaged cells, respectively, yielding a total of 88,884 single cells. For a more comprehensive analysis, we included 9,709 single cells from 20 mice, mostly ECs obtained from luminal digestion (Luminal Only) from our previous scRNA-seq datasets from the LCAs and RCAs exposed to d-flow alone at both 2 days and 2 weeks post-PCL time points^27^. Therefore, we analyzed a total of 98,553 single cells from 95 mice (75 mice from the present study and 20 mice from our previously published study since they were generated using the identical mouse model, single cell preparation, and scRNA-seq analytical pipeline) for downstream analysis.

We used the *Seurat* integration function to remove batch effects arising from combining scRNA-seq datasets generated from multiple experiments conducted at different times, including our prior study (Figure S4). There were 4 independent libraries, or batches, that were constructed, including 1 from the previous study (Table S2). Uniform manifold approximation and projection (UMAP) plot of the cells labelled by batch and by cell type revealed both a cohesive mixing of the different batches while maintaining the cell type identities upon integration compared to simple merging. Testing multiple benchmarking metrics using both all cells in the dataset and only the ECs showed that *Seurat* integration effectively mixed the batches while conserving biological variation of different cell types (Table S3)^31,32^.

UMAP clustering analysis of all single cells revealed 25 unique clusters representing 9 unique cell types (Figure 1B). Each cluster was manually annotated using canonical cell markers: 5 EC (*Pecam1*, *Cdh5*, *Icam2*, and *Tie1*), 3 vascular smooth muscle cell (SMC) (*Cnn1*, *Myh11*, and *Acta2*), 2 fibroblast (FB) (*Dpep1*, *Medag*, and *Pdgfra*), 5 macrophage (MΦ) (*Cd68*, *Adgre1*, *Fcgr1*, and *Itgam*), 3 dendritic cell (DC) (*Ccr7*, *Fscn1*, *Xcr1*, and *Cd209a*), 1 neutrophil (NT) (*S100a8* and *S100a9*), 1 B cell (BC) (*Cd79a* and *Cd79b*), 4 T cell (TC) (*Cd3d* and *Cd3g*), and 1 natural killer cell (NK) (*Klre1* and *Klra7*) clusters (Figure 1C) ^27,38,44–50^. Furthermore, the pan-leukocyte marker *Ptprc* (*Cd45*) was expressed only in the leukocytes (Leuko: MΦ, DC, NT, BC, TC, and NK) (Figure 1C). Importantly, none of the EC and SMC clusters expressed *Cd45*, excluding their leukocyte origin.

We next split the UMAP plot of the whole cell populations into 10 panels: 1) Ctrl_2 day (Luminal Only), 2) Ctrl_2 wk (Luminal Only + new Ctrl_2wk group), 3) Ctrl_4 wk, 4) HighChol_2 wk, 5) HighChol_4 wk, 6) D-flow_2 day (Luminal Only), 7) D-flow_2 wk (Luminal Only), 8) D-flow_4 wk, 9) D-flow_HighChol_2 wk, and 10) D-flow_HighChol_4 wk (Figure 1D). Note that panels 1, 6, and 7 contained single cells obtained from luminal digestion only from our previous study, while panel 2 contained luminal only cells and the new additional single cells from the LCAs and RCAs, consistent with the higher EC presence in these experimental groups (Figure S5)^27^. Comparison of the UMAP plot generated with or without the luminal only groups (Figure 1D, Panels 1, 2, 6, and 7) did not reveal any obvious changes to the unique clusters (data not shown), especially the EC clusters.

Hypercholesterolemia alone groups (Figure 1D, Panels 4-5, Figure S5) revealed no remarkable difference compared to the control groups (Panels 2-3). In contrast, d-flow alone for 4 weeks increased the appearance of different cell clusters, especially immune cells (Panel 8 vs. 3, Figure S5). Moreover, d-flow under hypercholesterolemia drastically increased the accumulation of distinct population of ECs (EC5), SMCs (SMC3), and MΦs (MΦ3 and MΦ4), especially at 4 weeks post-PCL (Panels 9-10, Figure S5). These results suggest that 1) hypercholesterolemia alone has minimal effects while d-flow alone induces significant changes in the arterial cell population and 2) the combination of d-flow and hypercholesterolemia induces a dramatic shift in arterial cell composition as plaques develop.

### D-flow under hypercholesterolemia induces reprogramming of ECs to immune-like and foam cells

To better understand the effects of d-flow alone, hypercholesterolemia alone, and d-flow under hypercholesterolemia on EC population change during atherogenesis, 5 EC clusters expressing canonical EC markers (*Pecam1*, *Cdh5*, *Icam2*, and *Tie1*) were further analyzed in depth (Figure 2A). Differential gene expression (DGE) analysis revealed 1,291 flow-sensitive genes that were significantly and differentially expressed with average log_2_Fold Change > 1 in the 5 EC clusters as shown in the heatmap (Figure S6A).

**Figure 2.**
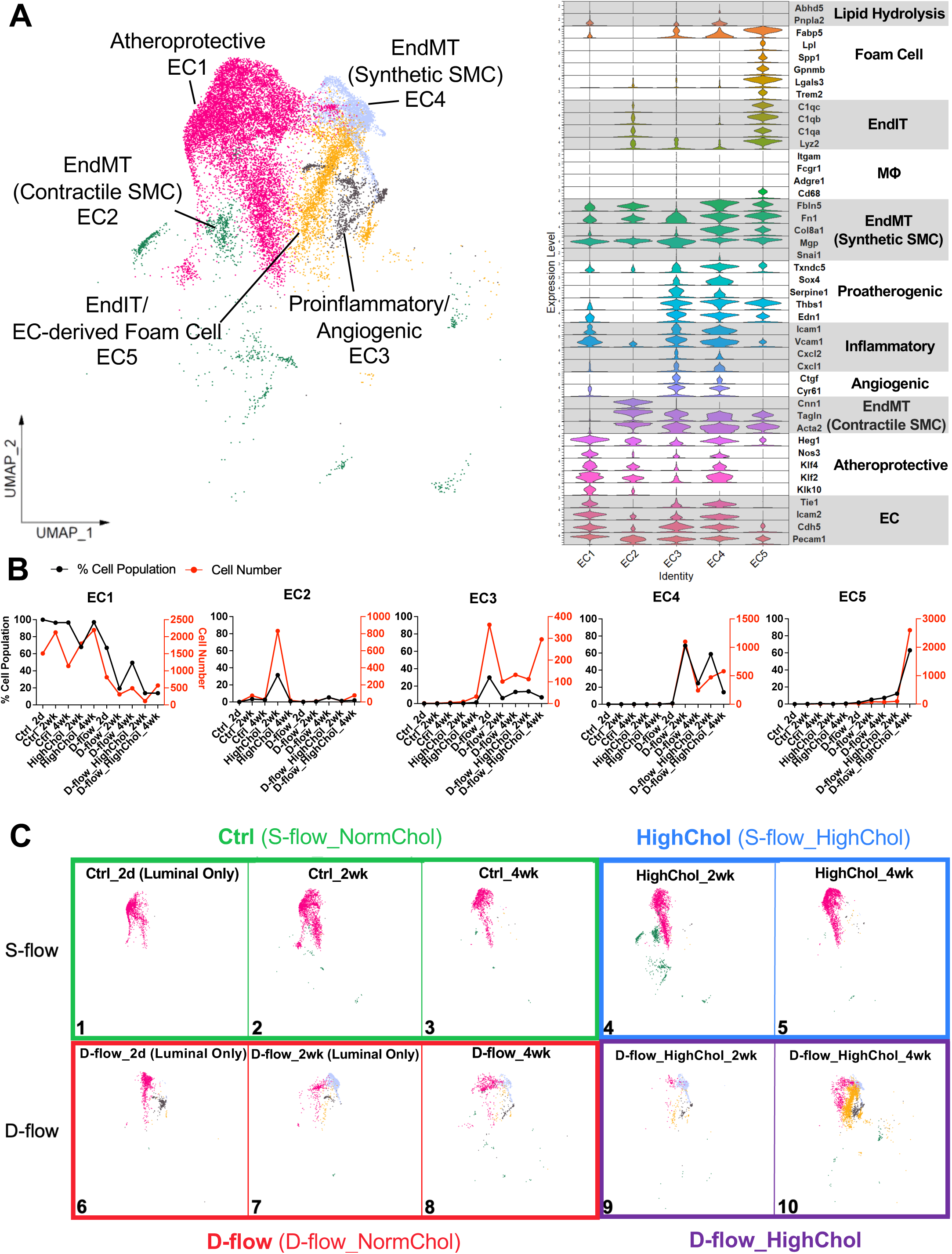
ECs transition to immune-like (EndIT) and foam cells (EndFT) by d-flow and hypercholesterolemia during atherogenesis. **A**. UMAP plot of 5 EC clusters (18,527 cells in total). Stacked violin plot shows expression levels of genes used to annotate each EC cluster. EC clusters include atheroprotective EC1, EndMT (contractile SMC) EC2, proinflammatory/angiogenic EC3, EndMT (synthetic SMC) EC4, and EndIT/EC-derived foam cell EC5. **B**. Cell number and % cell population (cell number for each EC cluster normalized by the total number of ECs per group) quantifications for each EC cluster across all 10 experimental groups. **C**. UMAP plot of each experimental group is shown for ECs. N= 5-20 mice for each condition. EndIT/EC-derived foam cells (EC5) dominate the EC population in the D-flow_HighChol at 4 weeks post-PCL (Panel 10).

Gene ontology (GO) analysis using the top 100 differentially expressed genes (DEGs) (Figure S6B) and literature-based predicted functions of each cell cluster (Figure 2A) suggested the following annotations: EC1 as healthy and atheroprotective ECs based on *Klk10*, *Klf2*, *Klf4*, *Nos3*, and *Heg1* expression; EC2 as ECs undergoing EndMT with contractile SMC markers (*Acta2*, *Tagln*, and *Cnn1*); EC3 as proinflammatory (*Cxcl1*, *Cxcl2, Vcam1,* and *Icam1*) and angiogenic ECs (*Cyr61* and *Ctgf*); and EC4 as ECs undergoing EndMT with synthetic SMC markers (*Mgp*, *Col8a1*, *Fn1*, and *Fbln5*). Both EC3 and EC4 clusters highly expressed many proatherogenic EC genes, including *Edn1*, *Thbs1*, *Serpine1*, *Sox4*, and *Txndc5*. Interestingly, EC5 included nearly 2,800 ECs (more than 60% of all ECs in D-flow_HighChol_4 wk group) abundantly expressing the markers of immune cells (*Cd68*, *Lyz2*, *C1qa*, *C1qb*, and *C1qc*), which we refer to as EndIT. In contrast, d-flow alone induced accumulation of only a few EC5 cells consistent to our previous report^27^ (Figure 2A, B). To our further surprise, EC5 highly expressed the markers of foam cells (*Trem2*, *Lgals3*, *Gpnmb*, *Spp1*, *Lpl*, and *Fabp5*) along with the canonical EC markers *Cdh5* and *Pecam1* albeit at a reduced level, which we refer to as a potential endothelial-to-foam cell transition (EndFT) occurring during atherogenesis^45^ (Figure 2A). In addition, EC5 cells showed a very low expression of markers of lipid droplet hydrolysis, *Pnpla2* (*Ctgl*) and its co-activator *Abhd5* (*Cgi58*), indicating that these ECs may accumulate endothelial lipid droplets as reported recently^51,52^ (Figure 2A).

We next analyzed the effects of hypercholesterolemia alone, d-flow alone, and d-flow under hypercholesterolemia on the distribution of EC clusters compared to the control. Hypercholesterolemia alone induced a transient increase in the EC2 (contractile EndMT) cells at 2 weeks (Figure 2C, Panel 4). In contrast, d-flow alone increased EC3 (proinflammatory/angiogenic) at 2 days, EC4 (synthetic EndMT) and just a few cells in EC5 (EndIT and EndFT) at 2 and 4 weeks post-PCL (Panels 6-8). Most importantly, d-flow under hypercholesterolemia dramatically increased the number of EC5 cells at 4 weeks post-PCL, which corresponds to robust plaque formation (Panel 10). These results indicate that d-flow alone initiates a partial FIRE, including inflammation, EndMT, and partial EndIT, while d-flow under hypercholesterolemia induces a robust FIRE, including full EndIT and the novel concept of EndFT, in addition to inflammation and EndMT, during atherogenesis.

### D-flow under hypercholesterolemia induces reprogramming of SMCs to foam cells during atherogenesis

We next examined the effects of d-flow alone, hypercholesterolemia alone, and d-flow under hypercholesterolemia on the SMC phenotypes. 3 SMC clusters were annotated using the markers of contractile (SMC1), synthetic (SMC2), and SMC-derived foam cells (SMC3) based on the DEGs, GO terms, and literature-based prediction of each cluster (Figure 3A, S7). SMC1 cells expressed contractile genes (*Acta2*, *Myh11*, *Tpm2*, *Myl9*, *Tagln*, *Cnn1*, and *Smtn*)^38,46,47^. Hypercholesterolemia alone did not induce any significant SMC phenotypic switches (Figure 3B, 3C Panels 4-5). D-flow alone at 2 and 4 weeks post-PCL induced significant accumulation of SMC2 cells expressing markers of synthetic SMCs (*Tpm4*, *Col8a1*, *Myh10*, *Mgp*, *Col1a1*, *Col3a1*, *Fn1*, *Dcn*, *Lum*, *Lox*, *Thbs1*, and *Fbln5*) (Figure 3B, 3C Panels 7-8)^38,46,47^. In contrast, d-flow under hypercholesterolemia induced a further increase in the number of synthetic SMC2 cells at 2 weeks post-PCL, while the number of SMC3 cells (SMC-derived foam cells) expressing markers of foam cells (*Trem2*, *Lgals3*, *Gpnmb*, *Spp1*, *Lpl*, and *Fabp5*) along with the canonical SMC marker *Acta2* albeit at a reduced level^45,53^ exploded at 4 weeks post-PCL (Figure 3B, 3C Panels 9-10). These results show that hypercholesterolemia alone had minimal effects on SMC phenotypic switch, while d-flow alone stimulates phenotypic switching to synthetic SMC2 cells. The combination of d-flow and hypercholesterolemia induces dramatic switching to SMC-derived foam cells (SMC3), consistent to prior reports^53,54^.

**Figure 3.**
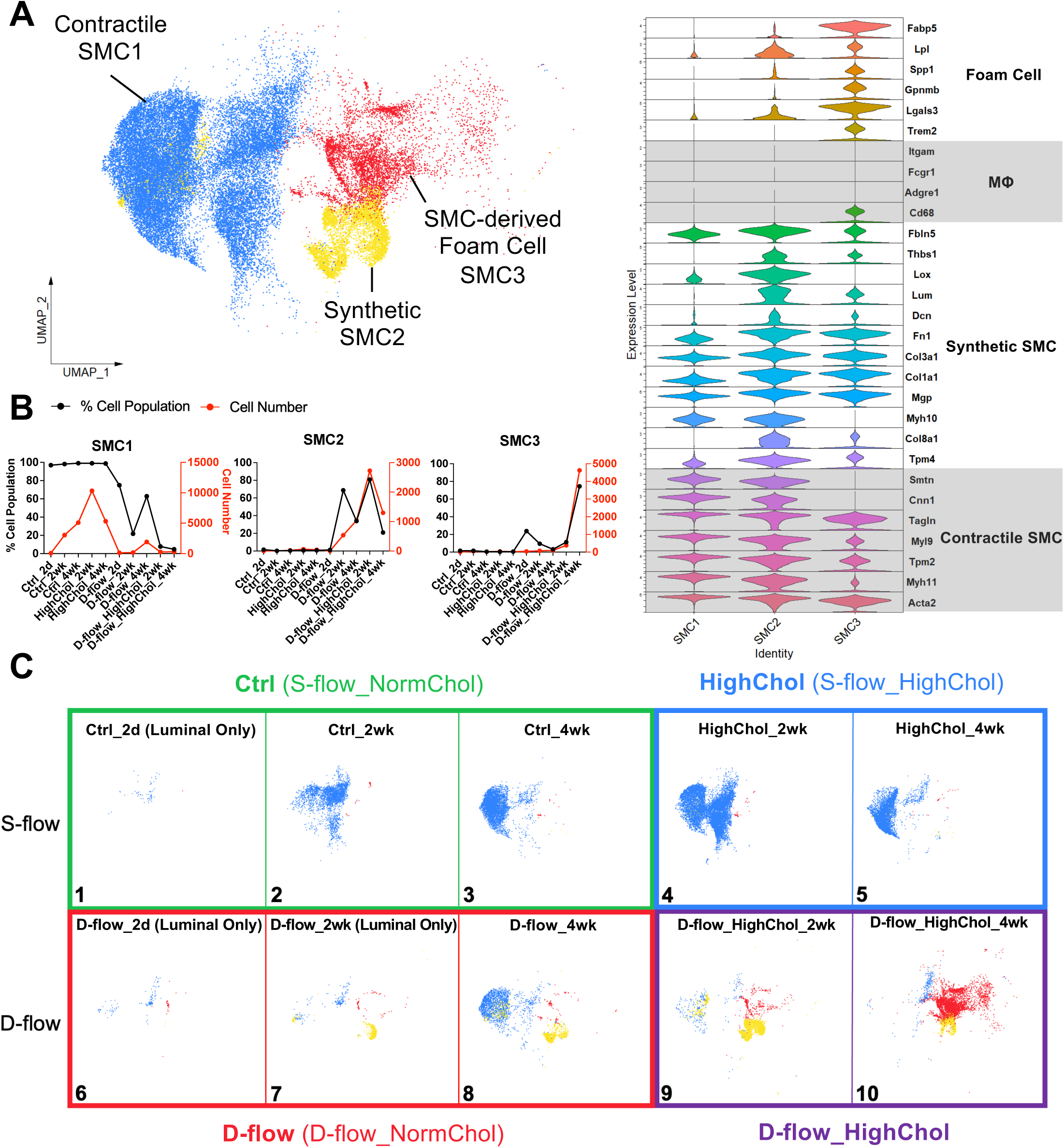
D-flow and hypercholesterolemia condition increases SMC-derived foam cells. **A**. UMAP plot of 3 SMC clusters (37,703 cells in total). Stacked violin plot shows expression levels of genes used to annotate each SMC cluster. SMC clusters include contractile SMC1, synthetic SMC2, and SMC-derived foam cell SMC3. **B**. Cell number and % cell population (cell number for each SMC cluster normalized by the total number of SMCs per group) quantifications for each SMC cluster across all 10 experimental groups. **C**. UMAP plot for each experimental group is shown for SMCs. N= 5-20 mice for each condition. Number of SMC-derived foam cells (SMC3) increase in the D-flow_HighChol at 4 weeks post-PCL (Panel 10).

### D-flow under hypercholesterolemia transforms MΦs into foam cells during atherogenesis

We identified 5 MΦ clusters expressing canonical MΦ markers (*Cd68*, *Adgre1*, *Fcgr1*, and *Itgam*) and predicted their functions based on the DEGs, GO terms, and the literature (Figure 4A, S8). MΦ5 cells expressing homeostatic, resident MΦ markers (*Lyve1*, *Pf4*, *F13a1*, *Mrc1*, *Cbr2*, *Folr2*, *Maf*, and *C4b*) were present in all conditions^45^ (Figure 4). Interestingly, hypercholesterolemia alone under s-flow condition had minimal effects on the MΦ population (Figure 4B, 4C Panels 4-5). It demonstrates the protective role of s-flow on prevention of monocyte infiltration into the arterial wall. In contrast, d-flow alone significantly increased the accumulation of MΦ1 (monocyte/macrophage) (Panel 6) and MΦ2 (inflammatory) (Panels 7-8) cells that highly expressed the markers of monocytes (*Nr4a1*, *Mef2a*, *Plac8*, *Ccr2*, *Ly6c2*, *Clec4e*, *F10*, *Cx3cr1*) at 2 days post-PCL and inflammatory markers (*Il1b*, *Tnf*, *Cxcl2*, *Ccl2*, *Ccl3*, *Ccl4*, and *Ccrl2*) at 2 and 4 weeks post-PCL, respectively (Figure 4B). In response to d-flow under hypercholesterolemia, all MΦ clusters significantly accumulated, most notably MΦ3 and MΦ4 that highly expressed the markers of foam cells (*Trem2*, *Lgals3*, *Gpnmb*, *Spp1*, *Lpl*, and *Fabp5*) (Figure 4B, 4C Panels 9-10). Interestingly, as plaques increase at 4 weeks post-PCL under hypercholesterolemic condition, MΦ4 cells expressing lower levels of efferocytosis markers (*Abca1*, *Axl*, *Cd36*, and *Ucp2*)^55,56^ than those of MΦ3 cells accumulated significantly (Figure 4B, 4C Panel 9 vs.10). Similar to the SMCs, hypercholesterolemia alone under s-flow condition had minimal effects on macrophage accumulation, whereas d-flow alone stimulated the initial accumulation of MΦ1 (monocyte/macrophage) and MΦ2 (inflammatory). In contrast, d-flow under hypercholesterolemia induced an explosion of all macrophages, especially macrophage-derived foam cells MΦ3 (*Abca1*^high^) and MΦ4 (*Abca1*^low^).

**Figure 4.**
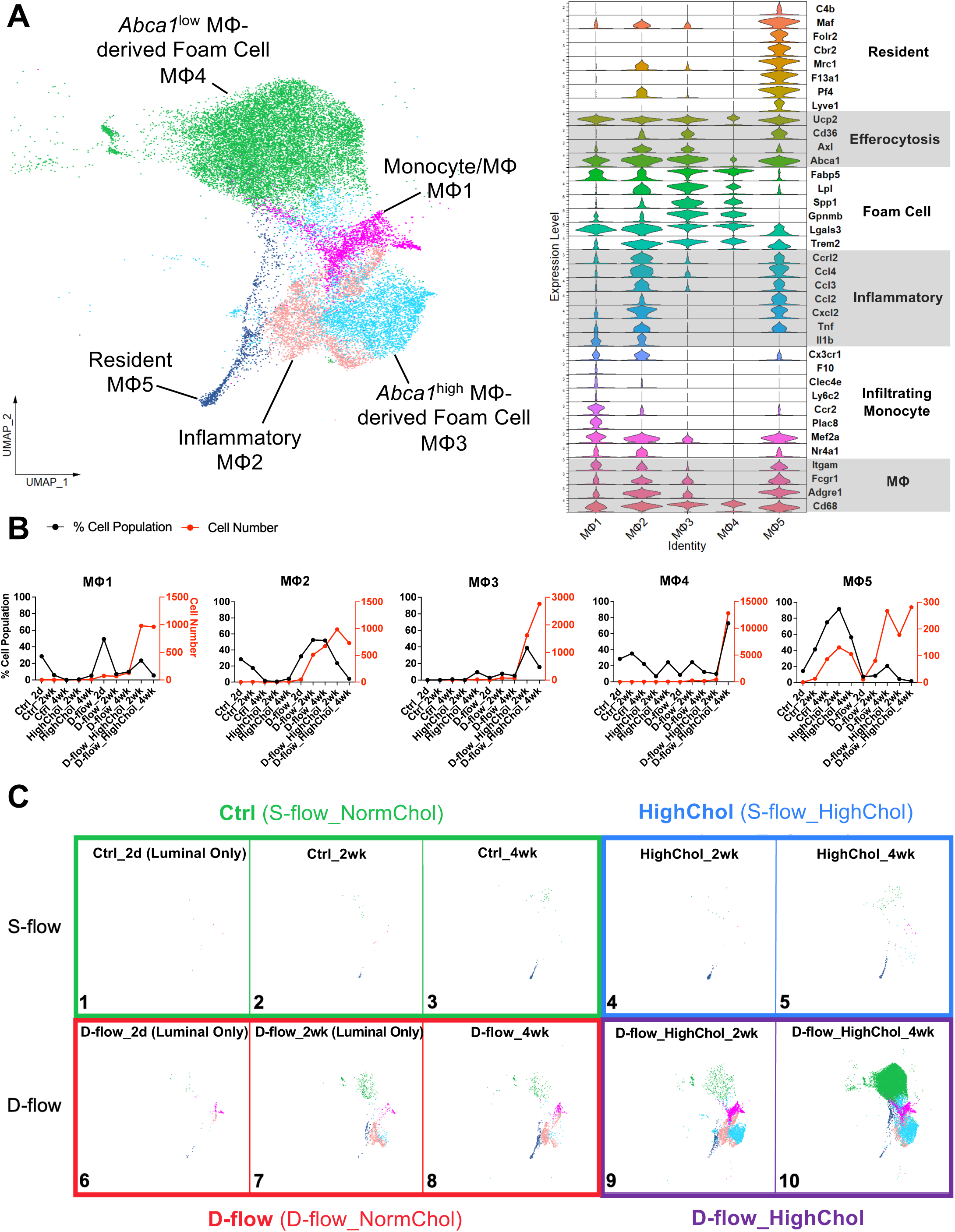
D-flow and hypercholesterolemia condition increases foamy MΦs. **A**. UMAP plot of 5 MΦ clusters (24,638 cells in total). Stacked violin plot showing expression levels of genes used to annotate each MΦ cluster. MΦ clusters include monocyte/MΦ MΦ1, inflammatory MΦ2, *Abca1*^high^MΦ-derived foam cell MΦ3, *Abca1*^low^ MΦ-derived foam cell MΦ4 and resident MΦ5. **B.** Cell number and % cell population (cell number for each MΦ cluster normalized by the total number of MΦs per group) quantifications for each MΦ cluster across all 10 experimental groups. **C**. UMAP plot for each experimental group is shown. N= 5-20 mice for each condition. MΦ-derived foam cells showing high expression levels of efferocytosis markers (*Abca1*, *Axl*, *Cd36*, and *Ucp2*) (MΦ3) are dominant in the D-flow_HighChol at 2weeks post-PCL (Panel 9), while MΦ-derived foam cells showing low expression levels of efferocytosis markers increase in the D-flow_HighChol at 4weeks post-PCL (Panel 10).

### D-flow induced accumulation of inflammatory and apoptotic fibroblasts and various immune cell types in the arterial wall during atherogenesis

We identified 2 fibroblast clusters and annotated as fibroblast progenitor/homeostatic fibroblasts (FB1) and inflammatory/apoptotic fibroblasts (FB2) using the same criteria discussed above (Figure S9)^48–50^. Our detailed analysis revealed that hypercholesterolemia alone had minimal effects while d-flow alone and d-flow under hypercholesterolemia increased FB2 population similarly (Figure S9B, C).

Analysis of other immune cells, including 4 TCs, 1 NKs, 3 DCs, 1 NTs, and 1 BCs, have been annotated using the canonical marker genes^45^ (Figures S10-12). Hypercholesterolemia alone had minimal effects on the accumulation of these cells. In contrast, d-flow alone increased the accumulation of most TC clusters, which were further increased significantly by d-flow under hypercholesterolemia (Figure S10).

Analysis of DCs, BCs, and NTs showed similar changes by d-flow alone and d-flow under hypercholesterolemia, but not by hypercholesterolemia alone (Figure S11, S12), demonstrating the dominant role of d-flow in immune cell infiltration during atherogenesis.

### Immune-like and foam cells derived from ECs share similar transcriptomic profiles with foam cells derived from MΦs and SMCs

The UMAP plot of our scRNA-seq data suggests an interesting possibility that EC-derived immune-like and foam cells (EC5) share similar transcriptomic profiles with foam cells derived from classical MΦs (MΦ3 and MΦ4) and SMCs (SMC3) during atherogenesis in response to d-flow under hypercholesterolemia, especially at 4 weeks post-PCL, as suggested from the close proximity of these foam cell clusters (Figure 1E, Panels 9-10). To address this hypothesis, multiple independent and unbiased trajectory analyses were performed using *destiny*, *Monocle 3*, and *scVelo* packages^34,35,37^.

Diffusion map trajectory analysis using *destiny* was performed using all EC, SMC, and MΦ clusters and projected into Diffusion Component space (Figure 5A, B). These projections indicated that ECs transitioned from the atheroprotective EC1 toward EndIT/EC-derived foam cell (EC5) phenotype, SMCs transitioned from the contractile SMC1 toward SMC-derived foam cell (SMC3) phenotype, and MΦs transitioned from the monocyte/macrophage (MΦ1) toward *Abca1*^high^ MΦ-derived foam cell (MΦ3) and then to *Abca1*^low^ MΦ-derived foam cell (MΦ4) phenotype. Although EC1, SMC1, and MΦ1 (initial cell populations) all originated from distinct nodes, the foam cells derived from ECs (EC5), SMCs (SMC3), and MΦs (MΦ3 and MΦ4) converged at the center (Figure 5B). Monocle 3 pseudotime trajectory analysis further confirmed the diffusion map analysis (Figure S13A, B). In addition, RNA velocity analysis was also carried out using *scVelo* to model cell fate transitions using the entire scRNA-seq data, further validating the diffusion map trajectory analysis (Figure S13C-E). We next quantified the % distribution of the foam cell origins by comparing EC5, SMC3, MΦ3, and MΦ4. As expected, MΦ3 (17%) and MΦ4 (52%) were the major source of foam cells (totaling 69%), while 20% of foam cells were derived from SMCs, with a surprising contribution of EC-derived foam cells at 11% (Figure 5C). These results for the first time revealed that ECs may undergo EndIT and EndFT. Moreover, foam cells could be derived not only from MΦs and SMCs but also from ECs during atherogenesis in response to d-flow under hypercholesterolemia.

**Figure 5.**
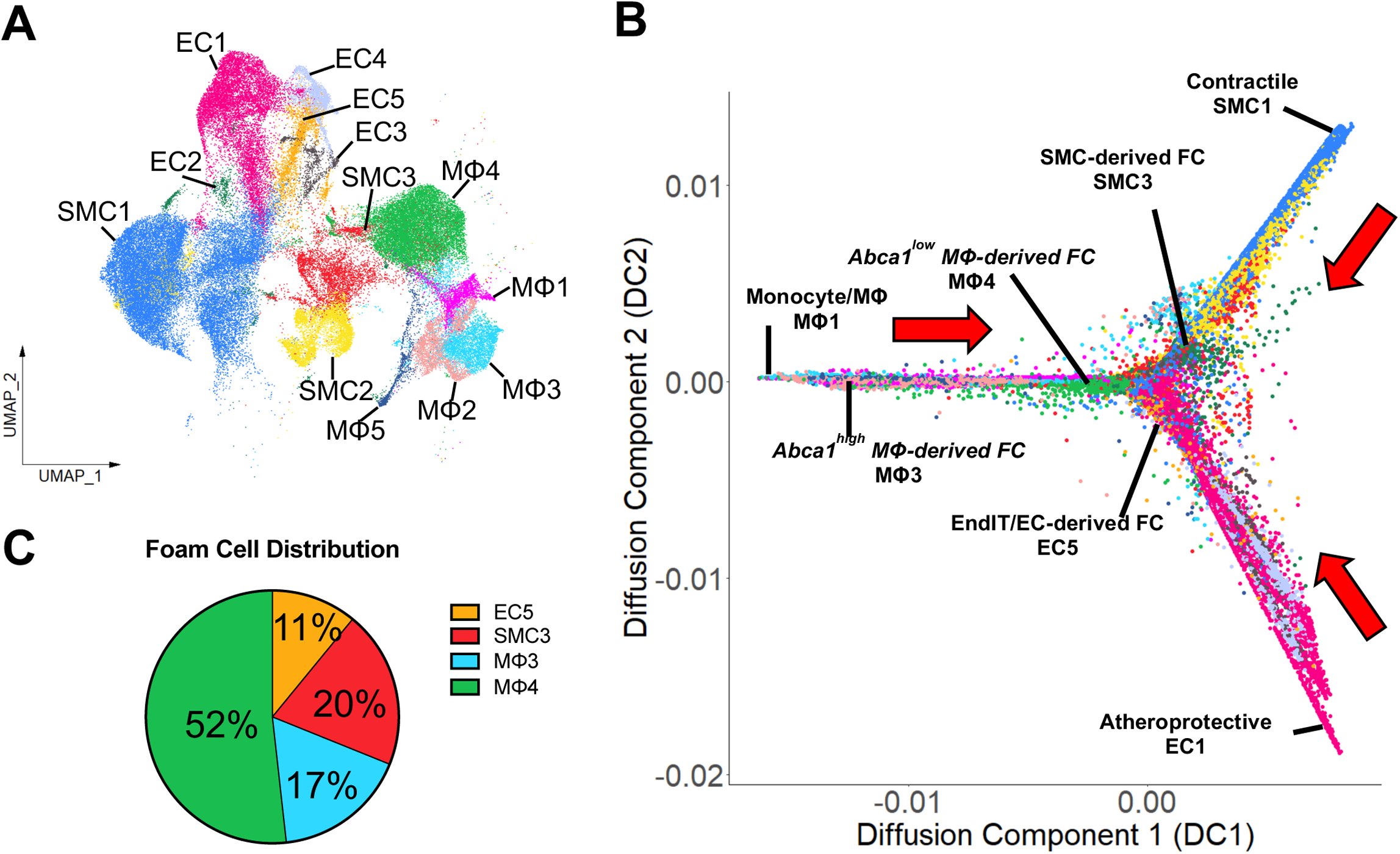
Transcriptomic profiles of foam cells derived from ECs are similar to those of foam cells derived from SMCs and MΦs. **A**. UMAP plot of all EC, SMC, and MΦ clusters used for further trajectory analysis in **B**. **B**. Diffusion map trajectory analysis of the 3 cell types reveals significant overlap of EC5 clusters with SMC3, MΦ3, and MΦ4 clusters from distinct origins. **C**. Pie chart showing % distribution of the origins of foam cells reveals 11% of foam cells are derived from ECs, 20% are derived from SMCs, and 69% are derived from MΦs.

To explore the potential cell-cell interaction pathways among all arterial cell clusters during atherogenesis, *CellChat* cell-cell communication analysis was performed on our scRNA-seq data^33^. We identified 113 predicted signaling pathways across all cell clusters, demonstrating similar levels of interactions among them without any dominant cell type or direction (Figure S14A). To identify cell-cell signaling pathways involving ECs undergoing EndIT and EndFT, we compared the ligand-receptor interactions that were sent by EC5 with those foam cells derived from SMCs (SMC3) and MΦs (MΦ3 and MΦ4) (Figure S14B). Some signals were uniquely sent by EC5, SMC3, and MΦ3/MΦ4, respectively, to all cell clusters, while there were some signals commonly sent by all foam cell clusters. EC5-originated interactions were Bmp4/Bmp6-Bmp receptors, Edn1-Ednra, Hspg2-Dag1, Thbs1-Itga3+Itgb1, Thbs1-Itgav+Itgb3, and Vwf-Itgav+Itgb3. SMC3-originated interactions were Col4a2-integrins/Cd44/Sdc and Postn-integrins. MΦ3 and MΦ4-originated interactions included Ccl3/4/6/9-Ccr1/2/5, Cd22-Ptprc, Cxcl2/4-Cxcr2/3, Il1a-Il1r1+Il1rap, Sema4d-Cd72/Plxnb2, Tgfb1-receptors, and Tnf-receptors. Interestingly, 3 signaling pathways, App-Cd74, Mif-receptors, and Spp1-receptors, were commonly sent by all foam cell clusters. In contrast, our attempt to identify ligand-receptor signals received by the foam cells revealed only App-Cd74 interaction (Figure S14C). Interestingly, MΦ3 foam cells were the most common receivers of many ligand-receptor interactions from various sender cells. These ligand-receptor interactions identified among arterial cells may play an important role in endothelial reprogramming and atherogenesis.

### EC lineage tracing model study validates EndIT and EndFT, in addition to inflammation and EndMT, during atherogenesis by d-flow under hypercholesterolemia

Next, we validated the endothelial reprogramming, especially the novel EndIT and EndFT revealed from our scRNA-seq study, while using endothelial inflammation and EndMT as controls by an independent immunohistochemical staining approach using an EC lineage tracing model. For this study, we used Cdh5-iCreER^T2^-Confetti^FL/WT^ mouse model expressing confetti markers, nuclear GFP, cytoplasmic YFP, or cytoplasmic RFP, specifically in ECs in a tamoxifen-inducible and stochastic manner (Figure S15A), as reported previously^42^. We hypothesized that the markers of endothelial reprogramming would be co-expressed in confetti^+^ ECs exposed to d-flow, such as the LCA in the PCL model and lesser curvature (LC) of the aortic arch naturally exposed to d-flow under hypercholesterolemic condition. Upon tamoxifen induction, EC-Confetti mice (6 males and 15 females) were injected with AAV-PCSK9, fed a high-fat diet, and underwent the PCL surgery, identical to the scRNA-seq study design (Figure S15B). Similar to the scRNA-seq study in C57BL/6 mice, atherosclerotic plaques developed specifically in the LCAs of the EC-Confetti mice but not in the RCAs at 4 weeks post-PCL (Figure S15C, D). PCSK9-mediated knockout of LDLR in the liver and hypercholesterolemia were also validated (Figure S15E-G).

For immunofluorescence staining of the FIRE markers, the RCA and two halves of the LCA were longitudinally sectioned as we previously reported^19^ (Figure 6A, B). We quantified confetti induction in the ECs across different flow conditions (RCA vs. LCA) in males and females by analyzing the number of confetti^+^ cells relative to the total number of luminal ECs of the carotid arteries. Approximately 16% and 17% of ECs expressed confetti in male and female RCAs which were significantly increased by 16% and 9% in male and female LCAs, respectively (Figure S15H). The results indicate that d-flow increased proliferation of confetti^+^ ECs, as expected^9,10^, in a sex-independent manner. Interestingly, we frequently observed patches of confetti+ ECs expressing the same fluorescently colored proteins, indicating a clonal expansion of ECs under d-flow conditions (Figure S15I) As controls, we first validated the protein markers of endothelial inflammation (Vcam1 and Icam1) and EndMT (Snai1, Acta2, and Cnn1) by immunofluorescence staining using specific antibodies that have been validated in the carotid arteries (LCA vs. RCA) and the aortic arch (LC vs. greater curvature (GC) exposed to s-flow) (Figure S16). To avoid any potential complication of these protein markers arising from the neighboring non-confetti ECs or other cell types, we focused on examining the confetti^+^ ECs in the luminal layer with minimal plaques. As expected, confetti^+^ ECs expressed Vcam1 and Icam1 (white signals, arrows) (Figure 6B-D, S), respectively, in the LCAs but not in the RCAs. In addition, EndMT markers (Snai1, Acta2, and Cnn1) were clearly and abundantly expressed in the confetti^+^ ECs in the lumen of LCAs (arrows) but not in the RCAs (Figure 6E-H, T, S17). Next, the other regions with more plaques were also examined for these markers, further demonstrating their co-expression in the confetti^+^ ECs (Figure S18A-E). These markers of inflammation and EndMT were also abundantly expressed in the confetti^+^ ECs in the LCs of the aortic arches but not in the GCs (Figure S19A-G). These results validate that endothelial inflammation and EndMT occur in the arterial regions of d-flow induced by surgery (LCA) or natural (LC) conditions, as expected^57–62^.

**Figure 6.**
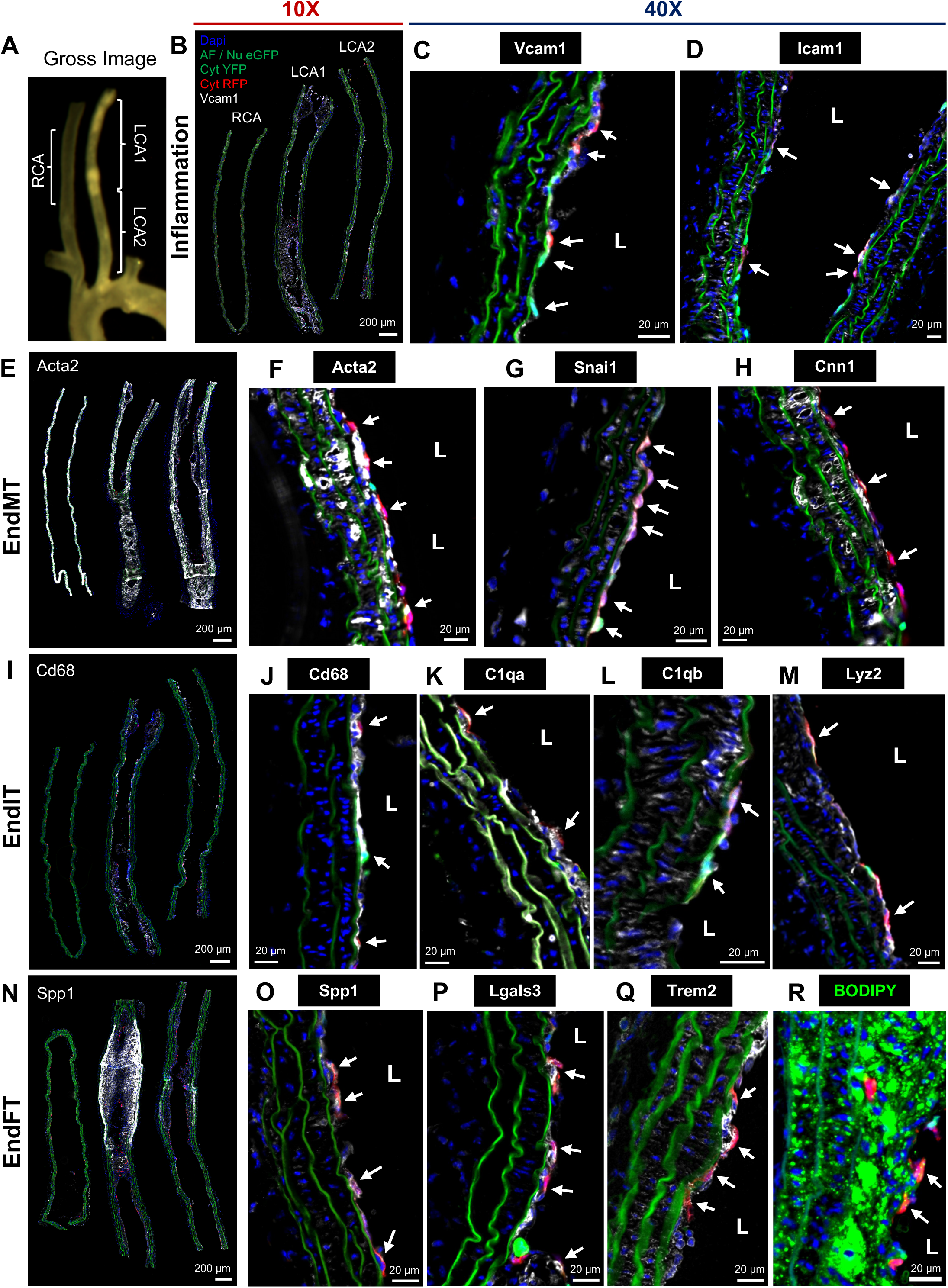
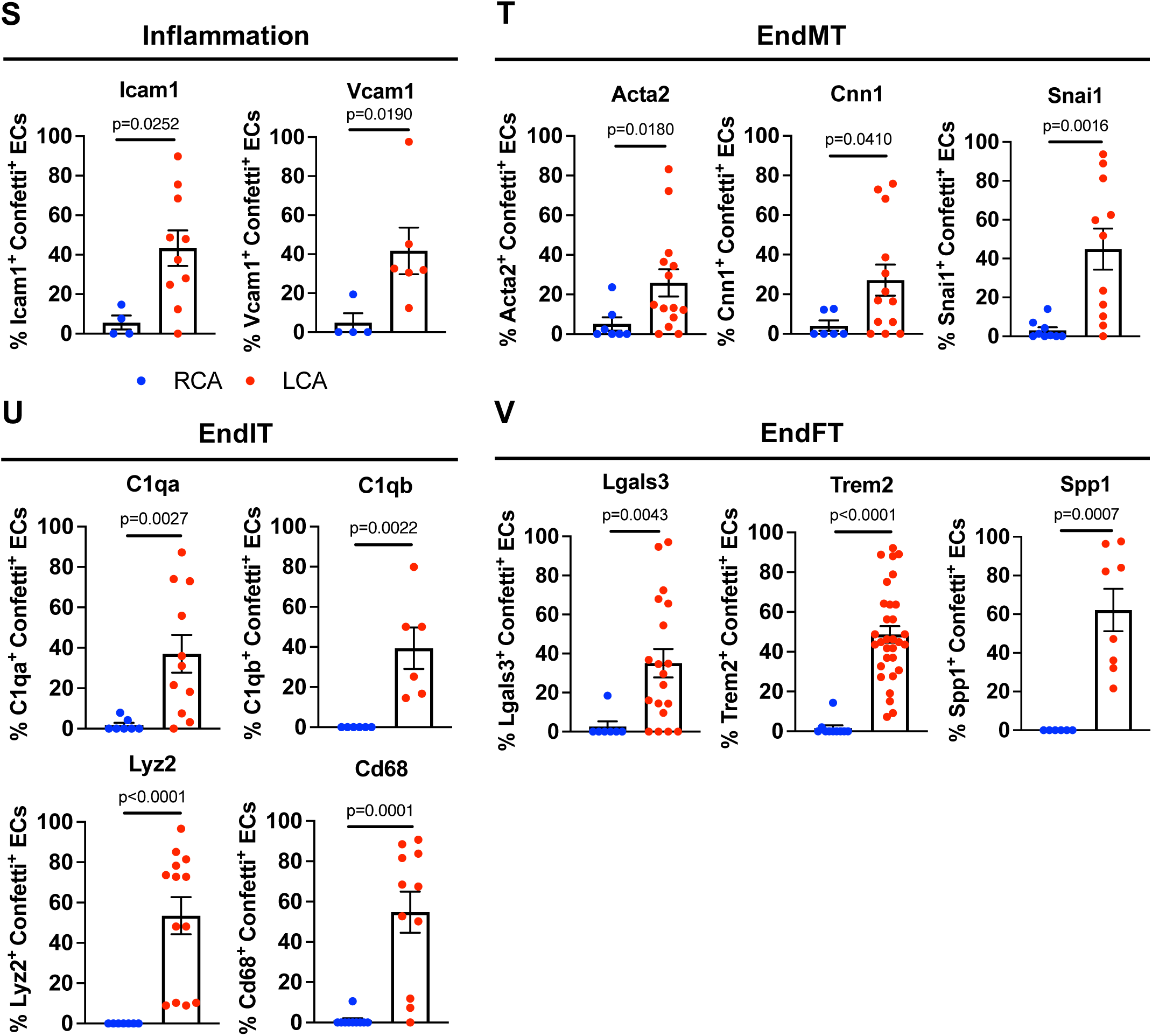
Lineage tracing study on EC-Confetti mice validates FIRE (endothelial inflammation, EndMT, EndIT, and EndFT) under d-flow and hypercholesterolemia at 4 weeks post-PCL. EC-Confetti mice treated with d-flow and hypercholesterolemia at 4 weeks post-PCL (N=6 male and 15 female) were imaged macroscopically (**A**) and LCAs/RCAs were longitudinally sectioned, stained, imaged by fluorescence microscopy (**B-R**), and quantified (**S-V**). **A** shows representative gross image of LCA, RCA, and aortic arch. **B-R**. LCAs and RCAs were immunostained with markers of endothelial inflammation (Vcam1 and Icam1, **B-D**); EndMT (Acta2, Snai1, and Cnn1, **E-H**); EndIT (Cd68, C1qa, C1qb, and Lyz2, **I-M**); and EndFT (Spp1, Lgals3, Trem2, and BODIPY, **N-R**). **B, E, I,** and **N** show merged images of confetti and FIRE markers at low magnification (10X), while the rest show 40X images. Confetti signals show eGFP (green), YFP (green), and RFP (red). All FIRE markers are shown in white except for green BODIPY (**R**). White arrows indicate confetti^+^ ECs co-expressing the FIRE markers. **S-V.** Percent confetti^+^ ECs co-expressing each FIRE marker was quantified by a combined Matlab and ImageJ analysis. Confetti^+^ ECs expressing markers of inflammation (Icam1 and Vcam1, **S**); EndMT (Acta2, Cnn1, and Snai1, **T**); EndIT (C1qa, C1qb, Lyz2, and Cd68, **U**); and EndFT (Lgals3, Trem2, and Spp1, **V**). Shown are mean ± SEM, each dot represents % of confetti^+^ ECs coexpressing FIRE markers in each longitudinal section used for quantification (N=4 to 10 longitudinal sections for RCA; N=6 to 31 longitudinal sections for LCA). *P* values were calculated by 2-tailed unpaired Student t test with or without Welch’s correction for normal data and 2-tailed unpaired Mann-Whitney U test for non-normal data.

Next, we validated the novel concept of EndIT and EndFT during atherogenesis in response to d-flow under hypercholesterolemia at the protein levels. Canonical protein markers of immune cells (Cd68, C1qa, C1qb, and Lyz2) and foam cells (Spp1, Lgals3, and Trem2) were examined in the carotid arteries and the aortic arches by immunostaining. To our surprise, we found robust expressions of Cd68, C1qa, C1qb, and Lyz2 in the confetti^+^ ECs in the lumen of the LCAs (arrows) but not in the RCAs (Figure 6I-M, U, S18F-I). The confetti^+^ ECs in the LCs also expressed the EndIT markers robustly but not in the GCs (Figure S19H-L). These results demonstrate that the confetti^+^ ECs undergo EndIT in response to d-flow under hypercholesterolemic condition during atherogenesis. Similarly, the foam cell markers (Spp1, Lgals3, and Trem2) were abundantly expressed in the confetti^+^ ECs in the lumen of the LCAs (arrows) but not in the RCAs (Figure 6N-Q, V, S18J-L). We also used BODIPY staining to determine lipid droplet accumulation in confetti^+^ ECs. We found 10 out of 36 LCA sections showed luminal confetti^+^ ECs with BODIPY^+^ lipid droplet accumulation (arrows), supporting foam cell formation derived from ECs (Figure 6R, S18M). However, confetti^+^ ECs stained with BODIPY in the LCs were rare (1 out of 25 sections) as shown by the rare example in Figure S19Q. Furthermore, Trem2 and Spp1 were expressed at a much lower frequency in the LCs compared to the LCAs (Figure S19M-P). These results show the correlation between the lack of plaque development due to less severe d-flow conditions and the reduced or lack of EndFT markers in aortic arches compared to the LCAs within 4 weeks, as they require more than 2 months to observe plaques in the LCs^13,29,63^.

Interestingly, there are many confetti^+^ ECs expressing FIRE markers found in the subendothelial and medial layers with or without plaques (Figure S18A-E, G-I, L). These results suggest that ECs undergoing FIRE, especially EndMT, EndIT, and EndFT, migrate into the subendothelial and medial layers. We also found luminal (Figure S18G, L) and medial (Figure S18C, I) confetti+ ECs with same confetti colors, indicating a potential clonal expansion and intraplaque angiogenesis (Figure S18A, C, I). To test whether the same ECs undergo EndMT, EndIT, and EndFT, we carried out co-immunofluorescence staining studies. Co-staining with Acta2 (green) and Cnn1 (white) showed that 34% of confetti^+^ ECs coexpressed these EndMT markers in the LCAs (Figure S20A, C). Moreover, 18% of confetti^+^ ECs coexpressed Acta2 and Cd68, indicating that EndMT and EndIT occur together within the same ECs (Figure S20B, C).

Taken together, these results demonstrate that d-flow under hypercholesterolemic condition induces robust endothelial reprogramming, including full EndIT and EndFT, in addition to endothelial inflammation and EndMT, during atherogenesis.

### Human scRNA-seq data from carotid atherosclerosis reveals EndIT and EndFT

To test the translational potential of the novel EndIT and EndFT in atherosclerosis, we reanalyzed the scRNA-seq results from human carotid atherosclerosis patient samples. For this study, the publicly available scRNA-seq data of 4,811 single cells collected from 44 human carotid endarterectomy samples was reanalyzed by *SingleR*-based reference mapping using our mouse cluster annotations^38,39^. The original UMAP plot of the human data showed 20 unique clusters, including 4 MΦ, 1 DC, 7 TC, 2 EC, 1 SMC, 2 NK, 1 mast cell (MC), and 2 BC clusters (Figure S21A). Our reanalysis of the human scRNA-seq data using our mouse cluster annotations predicts that human carotid endarterectomy samples contain not only MΦ-derived foam cells (MΦ3/4) and SMC-derived foam cells (SMC3) but also the novel EC-derived immune-like and foam cells (EndIT/EndFT EC5) (Figure S21B, C). This result suggests that ECs in the human carotid atherosclerotic plaques underwent EndIT and EndFT consistent with our mouse data.

### Reanalysis of Perturb-seq data reveals 40 flow-sensitive genes that regulate expression of FIRE markers

To investigate the roles of the 1,291 flow-sensitive genes that are significantly and differentially expressed by the 5 EC clusters on FIRE, we reanalyzed the recently published and publicly available CRISPRi-Perturb-seq data^41^. Briefly, Schnitzler et al. knocked down 2,285 genes related to 306 coronary artery disease (CAD)-GWAS signals at a single cell resolution in human ECs and identified sets of genes that may regulate CAD and EC-specific functional programs. Interestingly, 257 out of our 1,291 flow-sensitive genes (∼20%) from our scRNA-seq data were included among the 2,285 target genes in the Perturb-seq study (Figure 7A). They reported a total of 17,849 genes that were significantly altered by perturbing the 2,285 CAD-GWAS related genes. Interestingly, 1,045 of our 1,291 flow-sensitive genes (∼81%) were altered by the gene perturbations^41^. Moreover, 14 of our 1,045 flow-sensitive genes are FIRE markers altered by CAD-GWAS perturbations (Figure 7A), suggesting their potential importance in CAD.

**Figure 7.**
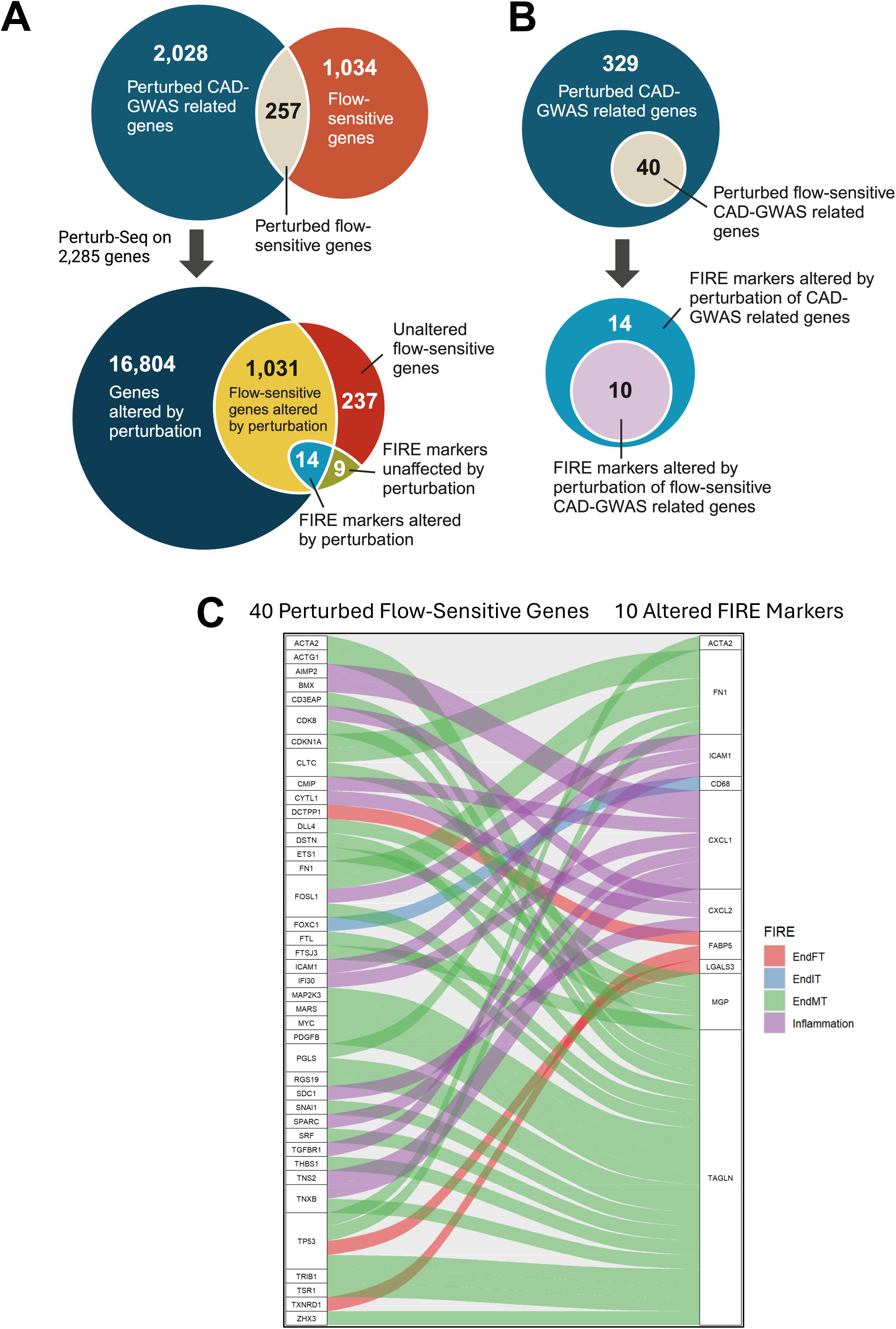
Reanalysis of Perturb-seq data on teloHAECs identifies 40 flow-sensitive genes as novel therapeutic targets for FIRE and atherosclerosis. CRISPRi-Perturb-seq data knocking down 2,285 genes associated with 306 CAD-GWAS signals that altered the expressions of 17,849 genes at a single cell resolution was reanalyzed and compared with 1,291 flow-sensitive DEGs in our FIRE scRNA-seq data. **A**. 257 of 1,291 flow-sensitive genes (∼20%) were perturbed due to their implication in CAD-GWAS (top), while 1,045 of 1,291 flow-sensitive genes (∼81%) were altered due to gene perturbations (bottom). 14 of these 1,045 altered flow-sensitive genes are FIRE markers used for EC cluster annotation. **B**. Perturbations of 329 CAD-GWAS related genes were responsible for altering the expressions of 14 FIRE markers, while 10 of these 14 FIRE markers were altered by 40 of 329 genes that were not only perturbed but are also flow-sensitive. C. Alluvial plot depicting the perturbation-gene pairs between the 40 perturbed flow-sensitive genes and 10 altered FIRE markers. Red: EndFT markers; Blue: EndIT markers; Green: EndMT markers; Purple: Inflammation markers.

Moreover, our reanalysis revealed that these 14 FIRE marker genes were regulated by 329 CAD-GWAS related genes, of which 40 are also flow-sensitive (Figure 7B). Of the 14 FIRE marker genes, 10 were perturbed by the 40 flow-sensitive and CAD-GWAS related genes. For example, knockdown of *TP53* altered markers of EndMT (*ACTA2*, *FN1*, *TAGLN*) and EndFT (*LGALS3*) (Figure 7C). EndMT markers (*FN1*, *MGP*, *TAGLN*) were regulated by many flow-sensitive genes. *FOXC1* knockdown altered an EndIT marker (*CD68*), while EndFT marker (*FABP5*) was regulated by *DCTPP1* and *TXNRD1*. These results provide a causal relationship between the flow-sensitive CAD-GWAS related genes and FIRE events.

## Discussion

Here, we showed that d-flow under hypercholesterolemic condition rapidly and robustly induced the novel EndIT and EndFT in addition to endothelial inflammation and EndMT, which we collectively coined as FIRE (flow-induced reprogramming of ECs), during atherosclerotic plaque development. These provocative, novel concepts of EndIT and EndFT were validated by comprehensive and extensive immunohistochemical staining studies using multiple protein and lipid markers in the EC-specific confetti mice. Further, these mouse-based concepts of EndIT and EndFT were supported by reanalysis of the published human carotid plaque scRNA-seq data^38,39^. These findings demonstrate that aortic ECs are highly plastic and heterogeneous and can transition to immune-like (EndIT) and foam cells (EndFT) in response to d-flow under hypercholesterolemic conditions during atherogenesis.

We previously showed that d-flow alone induces partial FIRE, including endothelial inflammation, EndMT, and partial EndIT in a few cells, in the mouse PCL model and in human aortic ECs *in vitro*^27^. As discussed in the Introduction, atherosclerosis preferentially develops in arterial regions exposed to d-flow conditions, such as branching points, in the presence of hypercholesterolemia^7,8,10^, demonstrating the interaction between the focal flow condition and systemic hypercholesterolemia through unclear mechanisms. Here, we tested the two-hit hypothesis that d-flow is the initial instigator of partial FIRE and systemic hypercholesterolemia is the major fuel, and the combination of the two leads to a full-blown FIRE and atherosclerotic plaque development (Figure 8).

**Figure 8.**
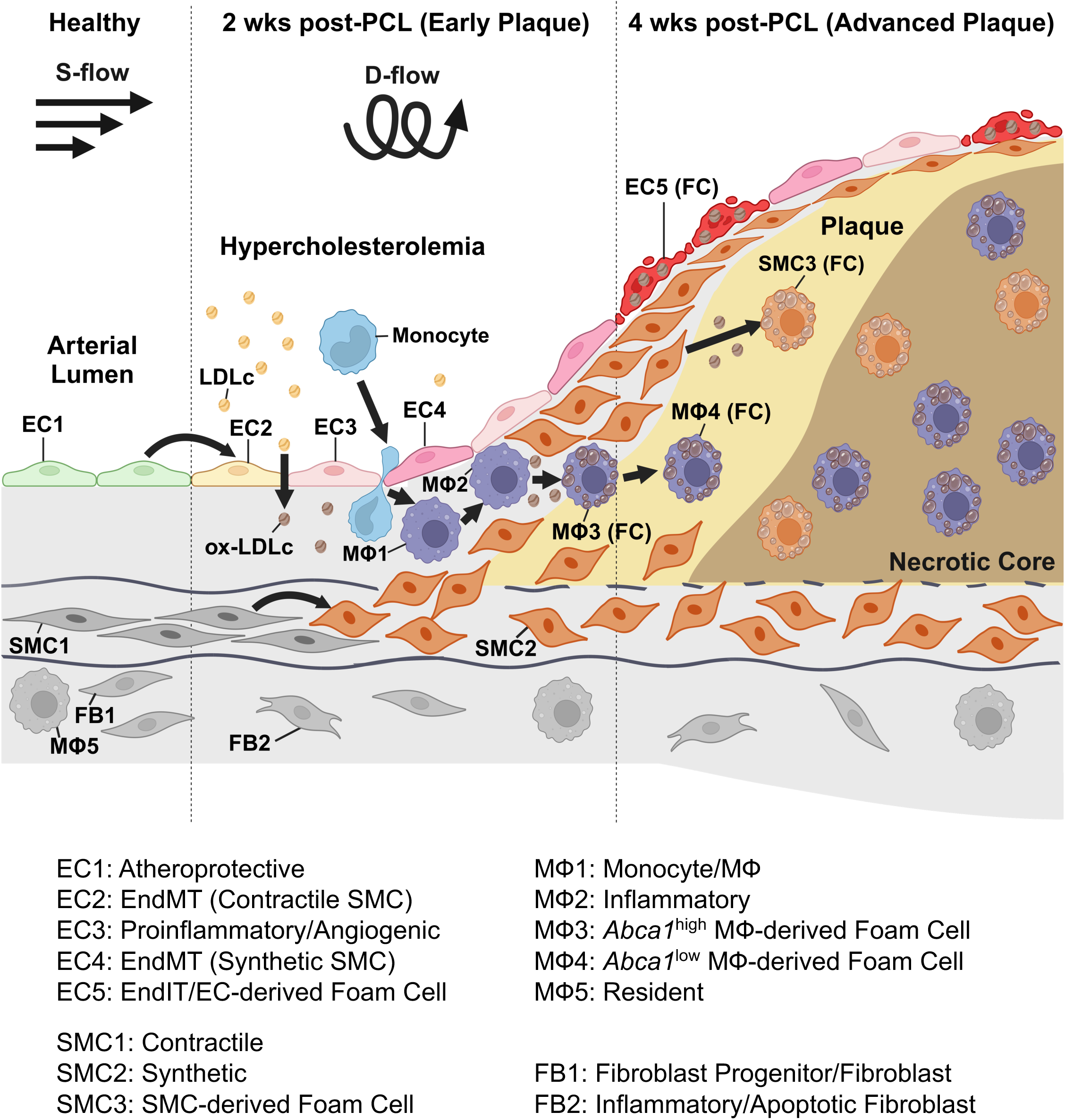
Summary and two-hit hypothesis of d-flow and hypercholesterolemia in atherogenesis. D-flow is the initial instigator of partial FIRE, including endothelial inflammation, EndMT, and partial EndIT. D-flow under hypercholesterolemic conditions triggers a robust FIRE, involving endothelial inflammation, EndMT, full EndIT, and EndFT, leading to atherosclerotic plaque development.

To test the two-hit hypothesis, we for the first time compared the effects of d-flow alone, hypercholesterolemia alone, or d-flow under hypercholesterolemia on single cell transcriptomics using more than 18,000 single ECs *in vivo*. Our data first confirmed that d-flow alone induces partial EndIT only in a few cells. Surprisingly, hypercholesterolemia alone had a minimal impact on arterial cell heterogeneity except for transient EndMT (EC2), indicating the dominant atheroprotective effect of s-flow even under the hypercholesterolemic conditions. In contrast, we found that d-flow under hypercholesterolemia induced a full and robust EndIT in the majority of ECs (∼60%) (Figure 2B), with the transcriptomic profiles that are nearly identical to those of foam cells derived from MΦs and SMCs (Figure 5B). Interestingly, those ECs undergoing EndIT/EndFT did not express most canonical MΦ markers, while expressing *Cd68* and other markers of immune and foam cells (Figure 2A), indicating that these are not of leukocyte-origin. These scRNA-seq results strongly supported the two-hit hypothesis.

We validated these provocative scRNA-seq data revealing the novel concepts of EndIT and EndFT by comprehensive immunohistochemical staining studies using an EC-specific lineage-tracing model. EC-specific confetti mice were used to examine whether genetically marked ECs with fluorescent confetti (GFP, YFP, or RFP) co-expressed the canonical markers of EndIT and EndFT, in addition to inflammation and EndMT. To minimize potential contributions from non-confetti ECs and other cell types, we initially focused on examining luminal confetti^+^ ECs adjacent to the internal elastic lamina with minimal plaques in the LCAs. These data were further validated in additional confetti^+^ ECs with plaques in the LCAs as well. We also demonstrated that these FIRE markers are co-expressed in confetti^+^ ECs in the LCs of aortic arches exposed to natural d-flow conditions in comparison to the LCAs exposed to the PCL model. In addition to the protein markers of EndFT, we also demonstrated lipid accumulation in the EndFT cells by using BODIPY staining in the confetti^+^ ECs of LCAs. In support of our EndFT concept, recent studies demonstrated that ECs indeed accumulate lipid droplets under hypercholesterolemic conditions^51,52^. We also found that EndFT cells lose the markers of lipid droplet hydrolysis, *Ctgl* and its co-activator *Cgi58*, which were reported to cause endothelial lipid droplet accumulation (Figure 2A).

We speculate the previously reported endothelial lipid droplets are related to the foam cell development reported here. Interestingly, BODIPY^+^/ confetti^+^ ECs were relatively rare in the LCs of aortic arches, which is due to a minimum plaque development in these areas under this acute condition. In addition, we found that the number of confetti^+^ ECs was significantly higher in the LCAs (Figure S14H) compared to the RCAs, indicating that d-flow induced proliferation of ECs, as expected^9,10^. Moreover, as shown in Figure S18G and L, we frequently observed patches of luminal ECs with the same confetti colors, suggesting a potential clonal expansion. In addition, some confetti^+^ ECs found in the subintimal and medial layers show same fluorescent colors, further indicating a potential clonal expansion (Figure S15I, S18C, I). However, these results and their pathophysiological implications need additional validation and studies.

We identified 4 different foam cell clusters derived from EC (EC5), SMC (SMC3), and MΦ (MΦ3 and MΦ4). Most surprisingly, our data for the first time demonstrate that foam cells are derived from not only the well-known MΦs and SMCs but also ECs. Our quantification indicates that 69% of foam cells are derived from MΦs, 20% from SMCs, and, to our surprise, 11% from ECs (Figure 5C). Previous studies showed that >50% and 60-70% of total foam cells in human plaques and ApoE^-/-^ mice, respectively, could be derived from SMCs as determined by flow cytometry and lineage tracing methods ^53,54,64^. The reasons for quantitative differences among these studies for the % of SMC-derived foam cells are unclear. Potential reasons include different mouse models (ApoE^-/-^ vs. C57BL/6 treated with AAV-PCSK9) and the quantitative methods (flow cytometry-based method vs. scRNA-seq). Nevertheless, in our model, we confirm the abundant presence of SMC-derived foam cells while reporting the novel EC-derived foam cells as well.

While the overall transcriptomic profiles of these foam cell clusters derived from ECs, SMCs, and MΦs largely overlap (Figure 5B), in-depth DEG analysis also showed their unique characteristics. Similar to the MΦ- and SMC-derived foam cells, EC-derived foam cells abundantly express the canonical markers of foam cells (*Trem2*, *Lgals3*, *Gpnmb*, *Spp1*, *Lpl*, and *Fabp5*) and their number increases in a time-dependent manner during atherogenesis (Figure 2)^45^. MΦ3 (*Abca1*^high^) that highly expresses the efferocytosis markers (*Abca1*, *Axl*, *Cd36*, and *Ucp2*)^55,56^ is predominant in early atherogenic phase at 2 weeks post-PCL under hypercholesterolemia. In contrast, EC5, SMC3, and MΦ4 (*Abca1*^low^) foam cells, predominant in advanced plaques at 4 weeks post-PCL under hypercholesterolemia, show a significantly reduced expression of efferocytosis markers (Figure S22), suggesting a potentially impaired efferocytotic capacity. Efferocytosis is a clearance of apoptotic cells by phagocytosis, which is effective during the early phase of atherosclerosis development but becomes impaired as atherosclerosis progresses, leading to necrotic core development^65^. Our results suggest an intriguing possibility that foam cells derived from ECs, SMCs, and *Abca1*^low^ MΦs may have an impaired efferocytotic capacity, leading to advanced plaque development.

Our current study identified and validated flow-sensitive genes involved in FIRE during atherosclerosis, revealing a treasure trove of information. Defining the genes and mechanisms regulating FIRE, especially the novel EndIT and EndFT, is a crucial future direction in understanding the role of flow in atherogenesis and developing anti-atherogenic therapies. Interestingly, a recent Perturb-seq study^41^ provides important direction and clues for the mechanisms. Our reanalysis of the Perturb-seq data identified 40 CAD-GWAS-related genes that are also flow-sensitive, regulating 10 FIRE markers involved in endothelial inflammation, EndMT, EndIT, and EndFT (Figure 7C). These results reveal the potential clinical importance of these flow-sensitive CAD-GWAS genes in FIRE and atherosclerosis for both mechanistic insights and therapeutic targets.

There are several limitations in this study. Due to the low number of cells (mostly ECs and immune cells) collected in the luminal digestion carotid samples, we had to pool single cell preparations from 5 or 10 mice to obtain a sufficient number of single cells for sequencing. With this approach of pooling the single cells from multiple mice, the mouse-level information for each single cell is lost, making it difficult to conduct statistical analysis across different conditions. However, to overcome this limitation, we conducted extensive immunostaining studies using the EC-Confetti mice for a genetic lineage-tracing study (Figure 6). Our extensive immunostaining studies using 12 different antibodies and a lipid staining method (BODIPY) in N=21 mice (6 males and 15 females) validate our key conclusions on FIRE revealed by the scRNA-seq study.

Furthermore, the underlying mechanisms responsible for EndIT and EndFT under d-flow and hypercholesterolemic conditions are unclear. The potential list of genes and pathways responsible for these pathways includes more than 1,000 flow-sensitive genes revealed in this study. Systematic screening approaches are needed to address these mechanisms. The current scRNA-seq study was conducted using only male mice due to the high cost of scRNA-seq studies. We addressed this limitation by carrying out validation studies using both male and female EC-specific confetti mice, which strongly validated the novel EndIT and EndFT during atherogenesis in a sex-independent manner.

Furthermore, we induced hypercholesterolemia in our mice using a combination of Western diet and AAV-PCSK9. AAV-PCSK9 was shown to induce systemic inflammation^66^, which may have affected our results on EndIT and EndFT. However, hypercholesterolemia alone caused by AAV-PCSK9 did not induce EndIT and EndFT, although EndMT was transiently increased at 2 wks but not at 4 wks post-PCL. The % confetti expression depends on when the tamoxifen injection is performed. Treatment with tamoxifen at an earlier time point at 4 weeks induced much higher confetti induction than those induced at 6 weeks, reflecting that their carotid EC development is reaching a young adult level with a relatively low EC proliferation rate. In our initial studies, mice were tamoxifen-treated at 6 wks, but we switched to an earlier timepoint at 4 wks. Nevertheless, the low confetti labelling efficiency of ECs remains a limitation.

In summary, we demonstrate that d-flow and hypercholesterolemia are two major atherogenic hits, inducing a full-blown FIRE, especially the novel EndIT and EndFT, as well as endothelial inflammation and EndMT, and atherosclerotic plaque development (Figure 8). The comprehensive single-cell atlas of mouse atherosclerotic plaques and the flow-sensitive genes, proteins, and pathways under d-flow and hypercholesterolemic conditions identified in this study are invaluable resources for mechanistic understanding and therapeutic targets of atherosclerosis.

## Acknowledgments

CP, KB, AP, and HJ conceptualized and planned the experiments. CP, KB, RH, LC, KJ, PK, AKJ, JHK, DWK, EJS, YK, JABK, and MDC performed the experiments. CP, KB, RH, LC, KJ, PK, MM, AK, CC, and NVR analyzed experimental results. SWL and GP contributed critical materials. CP, KB, and HJ wrote and edited the final manuscript with input and approval from all authors.

The authors are grateful to Dr. Ralf Adams for providing tamoxifen-inducible, endothelial specific Cdh5-iCreER^T2^ mice. We are also grateful to Dr. Jian Hu at Emory University for providing consultation on the statistical analysis of the scRNA-seq data. This study was supported in part by the Emory Integrated Genomics Core (EIGC) (RRID:SCR_023529), which is subsidized by the Emory University School of Medicine and is one of the Emory Integrated Core Facilities. Additional support was provided by the Georgia Clinical & Translational Science Alliance of the National Institutes of Health under Award Number UL1TR002378. The content is solely the responsibility of the authors and does not necessarily reflect the official views of the National Institutes of Health. Microscopy data for this study were acquired and/ or analyzed in the Microscopy in Medicine Core.

## Data Availability Statement

The datasets presented in this study can be found in NCBI BioProject repository (accession number: PRJNA1112537).

## Non-standard Abbreviations and Acronyms

AA: Aortic arch
Acta2: Actin alpha 2, smooth muscle
ARI: Adjusted rand index
ASW: Average silhouette
width BC: B cell
Cft: Confetti
Cnn1: Calponin 1
Ctrl: Control
DC: Dendritic cell
DEG: Differentially expressed gene
D-flow: Disturbed flow
DGE: Differential gene expression
EC: Endothelial cell
EndFT: Endothelial-to-foam cell transition
EndIT: Endothelial-to-immune cell-like transition
EndMT: Endothelial-to-mesenchymal cell transition
FB: Fibroblast
FBS: Fetal bovine serum
GC: Greater curvature
GFP: Green fluorescent protein
GO: Gene ontology
HighChol: Hypercholesterolemia
kBET: k-nearest neighbor batch effect test
LC: Lesser curvature
LCA: Left common carotid artery
Leuko: Leukocyte
Lgals3: Galectin 3
LISI: Local inverse Simpson’s index
Lyz2: Lysozyme 2
MC: Mast cell
MΦ: Macrophage
NT: Neutrophil
PCL: Partial carotid ligation
PCSK9: Proprotein convertase subtilisin/kexin type 9
RCA: Right common carotid artery
RFP: Red fluorescent protein
S-flow: Stable flow
SMC: Vascular smooth muscle cell
scATAC-seq: Single-cell assay for transposase-accessible chromatin sequencing
scRNA-seq: Single-cell RNA-sequencing
Snai1: Snail family transcriptional repressor 1
Spp1: Secreted phosphoprotein 1
Trem2: Triggering receptor expressed on myeloid cells 2
TC: T cell
UMAP: Uniform manifold approximation and projection
WD: Western diet
YFP: Yellow fluorescent protein

