## Supplemental Figures for "Disturbed Flow Induces Reprogramming of Endothelial Cells to Immune-like and Foam Cells under Hypercholesterolemia during Atherogenesis"

### Supplementary Table 1. Antibody information

| Antibodies | Vendor | Catalog # | Dilution |
|---|---|---|---|
| Recombinant Anti-TREM2 antibody | Abcam | ab245227 | 1:500 |
| Mouse Osteopontin/OPN Antibody | R&D | MAB808 | 1:20 |
| Recombinant Anti-Galectin 3 antibody | Abcam | ab76245 | 1:1000 |
| Recombinant Anti-alpha smooth muscle Actin antibody | Abcam | ab150301 | 1:100 |
| Recombinant Anti-Calponin 1 antibody | Abcam | ab46794 | 1:100 |
| SNAIL Polyclonal Antibody | Thermo Fisher Scientific | PA5-85493; RRID: AB_2792633 | 1:500 |
| Anti-C1QA antibody | Abcam | ab155052 | 1:200 |
| Lysozyme Recombinant Rabbit Monoclonal Antibody | Thermo Fisher Scientific | MA5-32154; RRID: AB_2809443 | 1:100 |
| C1QB Polyclonal Antibody | Thermo Fisher Scientific | PA5-102737; RRID: AB_2852128 | 1:100 |
| Anti-ICAM1 antibody | Abcam | ab171123 | 1:500 |
| Recombinant Anti-VCAM1 antibody | Abcam | ab134047 | 1:500 |
| Goat anti-Rabbit IgG (H+L) Highly Cross-Adsorbed Secondary Antibody, Alexa Fluor™ 647 | Thermo Fisher Scientific | A-21245; RRID: AB_2535813 | 1:500 |
| Goat Anti-Rabbit IgG (H+L) Cross Adsorbed Secondary Ab Alexa Fluor 568 | Thermo Fisher Scientific | A-11011; RRID: AB_143157 | 1:500 |
| Mouse IgG Isotype Control | Thermo Fisher Scientific | 31903 | Matched to primary antibodies |

### Supplementary Table 2. scRNA-seq sample information

| Original Sample | Batch | Grouped Sample **(Panel # in UMAP plot split by group)** | # of Mice (N) | # of Carotid Arteries | Mean Reads per Cell | Mean Genes per Cell | # of Cells Sequenced | # of Cells Analyzed | # of Cells from Luminal Digestion | # of Cells from Leftover Digestion |
|---|---|---|---|---|---|---|---|---|---|---|
| Luminal_S-flow_2d | Previous Study | Ctrl_2d (**1**) | 10 | 10 | 17,700 | 3,669 | 1,824 | **1,634** | 1,634 | 0 |
| Luminal_D-flow_2d | Previous Study | D-flow_2d (**6**) | | 10 | 19,122 | 3,583 | 2,032 | **1,878** | 1,878 | 0 |
| Luminal_S-flow_2wk | Previous Study | Ctrl_2wk (**2**) | 10 | 10 | 17,630 | 3,750 | 1,234 | **1,102** | 1,102 | 0 |
| Luminal_D-flow_2wk | Previous Study | D-flow_2wk (**7**) | | 10 | 14,223 | 3,060 | 4,127 | **3,793** | 3,793 | 0 |
| Ctrl_2wk | 2wk Experiment | Ctrl_2wk (**2**) | 5 | 10 | 4,892 | 1,501 | 5,903 | **5,368** | 609 | 4,759 |
| HighChol_2wk | 2wk Experiment | HighChol_2wk (**4**) | 5 | 10 | 13,588 | 3,536 | 4,157 | **3,924** | 107 | 3,817 |
| S-flow_HighChol_2wk | 2wk Experiment | HighChol_2wk (**4**) | 10 | 10 | 6,367 | 1,994 | 11,352 | **10,860** | 788 | 10,072 |
| D-flow_HighChol_2wk | 2wk Experiment | D-flow_HighChol_2wk (**9**) | | 10 | 16,470 | 3,789 | 13,140 | **11,793** | 5,703 | 6,090 |
| Ctrl_4wk | 4wk Experiment 1 | Ctrl_4wk (**3**) | 10 | 20 | 11,975 | 3,250 | 15,732 | **4,605** | 383 | 4,222 |
| HighChol_4wk | 4wk Experiment 1 | HighChol_4wk (**5**) | 10 | 20 | 14,124 | 3,590 | 12,309 | **5,088** | 678 | 4,410 |
| S-flow_HighChol_4wk | 4wk Experiment 1 | HighChol_4wk (**5**) | 20 | 20 | 11,346 | 3,013 | 8,538 | **4,723** | 1,917 | 2,806 |
| D-flow_HighChol_4wk | 4wk Experiment 1 | D-flow_HighChol_4wk (**10**) | | 20 | 3,424 | 1,016 | 59,151 | **32,052** | 29,696 | 2,356 |
| S-flow_4wk | 4wk Experiment 2 | Ctrl_4wk (**3**) | 15 | 15 | 14,450 | 3,481 | 3,306 | **3,099** | 324 | 2,775 |
| D-flow_4wk | 4wk Experiment 2 | D-flow_4wk (**8**) | | 15 | 12,592 | 3,192 | 9,067 | **8,634** | 338 | 8,296 |
| **Total** | | | 95 | 190 | | | 151,872 | **98,553** | 48,950 | 49,603 |

**Supplementary Table 3. Benchmarking metrics for batch mixing**

| Metric | Merging (All Cells) | *Seurat* Integration (All Cells) | Merging (ECs Only) | *Seurat* Integration (ECs Only) |
|---|---|---|---|---|
| **ARI** | 0.3635984 | 0.4669325 | N/A | N/A |
| **iLISI** | 1.191525 | 1.654378 | 1.215523 | 1.77661 |
| **cLISI** | 1.115989 | 1.1321798 | N/A | N/A |
| **ASW_batch** | N/A | N/A | 0.1334283 | -0.011824 |
| **ASW_celltype** | N/A | N/A | N/A | N/A |
| **kBET (10% sample size)** | 0.9983553 | 0.9979086 | 0.9992233 | 0.9974757 |
| **kBET (20% sample size)** | 0.9998528 | 0.999736 | 1 | 1 |

ARI: Adjusted rand index; LISI: local inverse Simpson's index; iLISI: integration LISI; cLISI: cell type LISI; ASW: average silhouette width; kBET: k-nearest neighbor batch effect test; N/A: not available

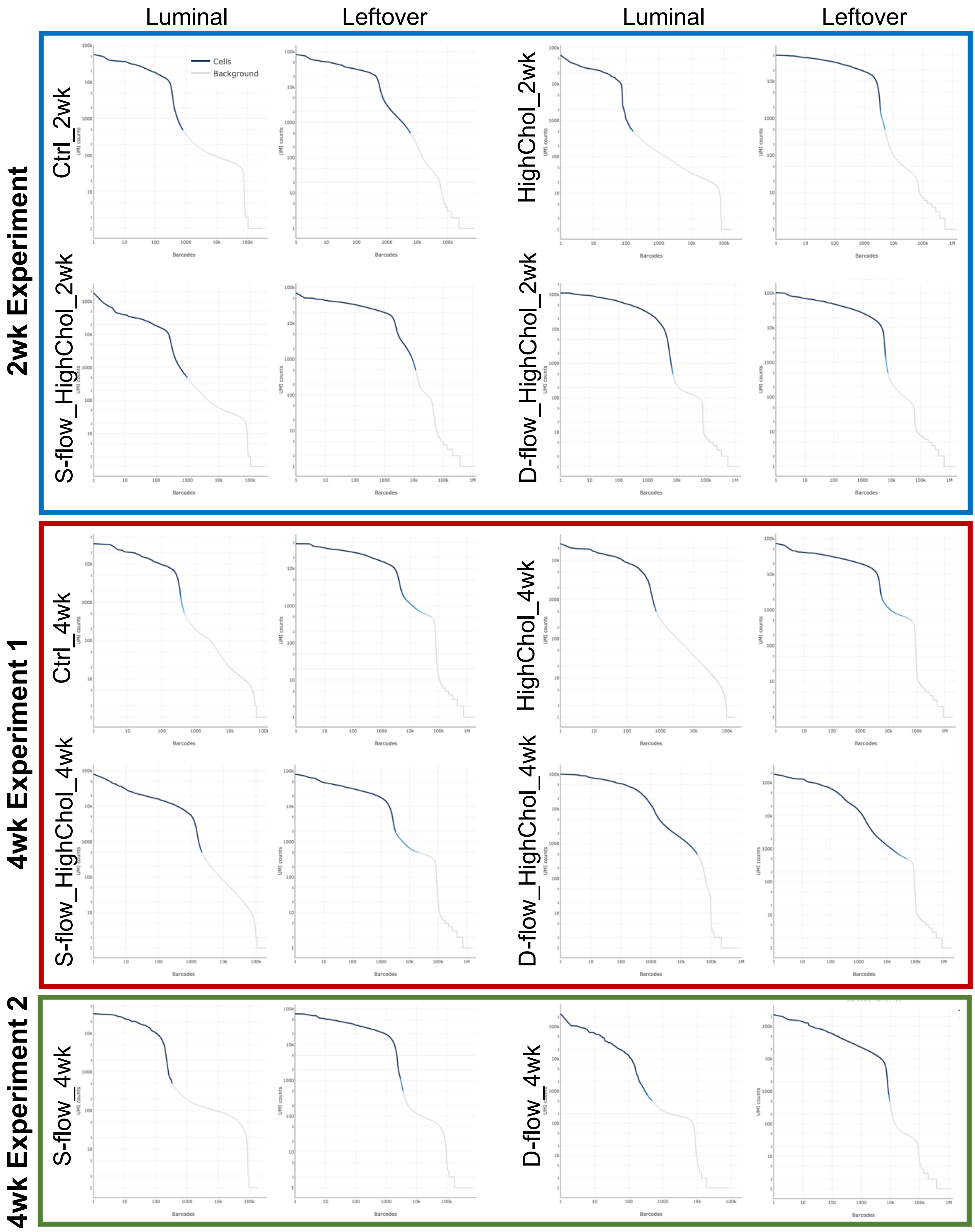
Luminal
Leftover
Luminal
Leftover
2wk Experiment
Ctrl_2wk
HighChol_2wk
S-flow_HighChol_2wk
D-flow_HighChol_2wk
Cells
Background
4wk Experiment 1
Ctrl_4wk
HighChol_4wk
S-flow_HighChol_4wk
D-flow_HighChol_4wk
4wk Experiment 2
S-flow_4wk
D-flow_4wk
UMI counts
Barcodes

**Supplementary Figure 1. Cell Ranger barcode ranked plots for each experimental group.** Cell Ranger barcode ranked plots for experimental group, organized by batches of libraries constructed: 2wk Experiment, 4wk Experiment 1, and 4wk Experiment 2.

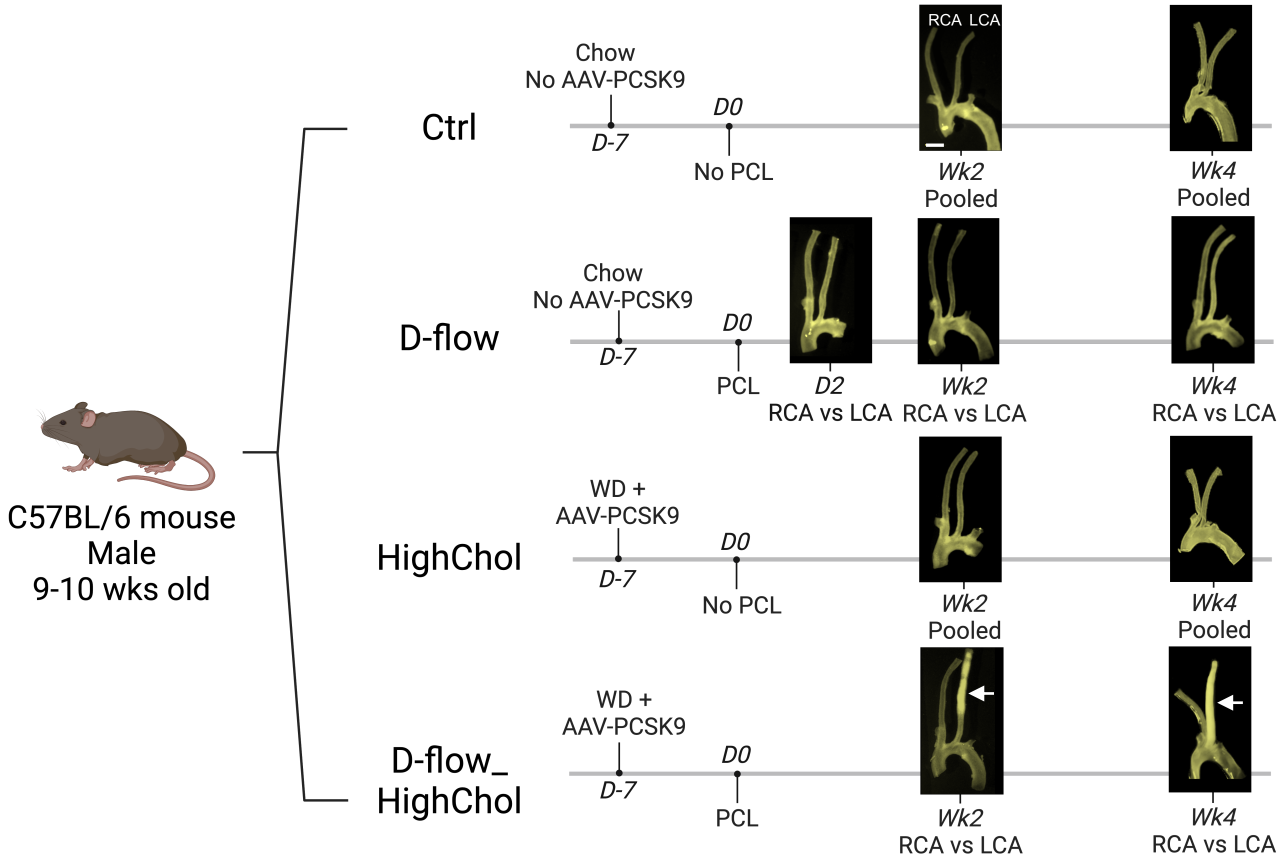


**Supplementary Figure 2. Experimental design.**
C57/BL6 mice (9-10-week-old, N = 75 Male mice) were treated with 4 experimental conditions (Ctrl, D-flow alone, HighChol alone, and D-flow_HighChol). To induce hypercholesterolemia, mice were injected with AAV-PCSK9 injection and fed Western diet. D-flow in the LCA was then induced by PCL surgery 1 week later. Ctrl and D-flow alone groups were fed a chow diet, while HighChol and D-flow_HighChol groups were fed Western diet for 2 or 4 weeks following the PCL surgery. Atherosclerotic plaques developed at 2 and 4 weeks post-PCL only in the D-flow_HighChol groups as shown by representative macroscopic images. Scale bar = 1 mm.

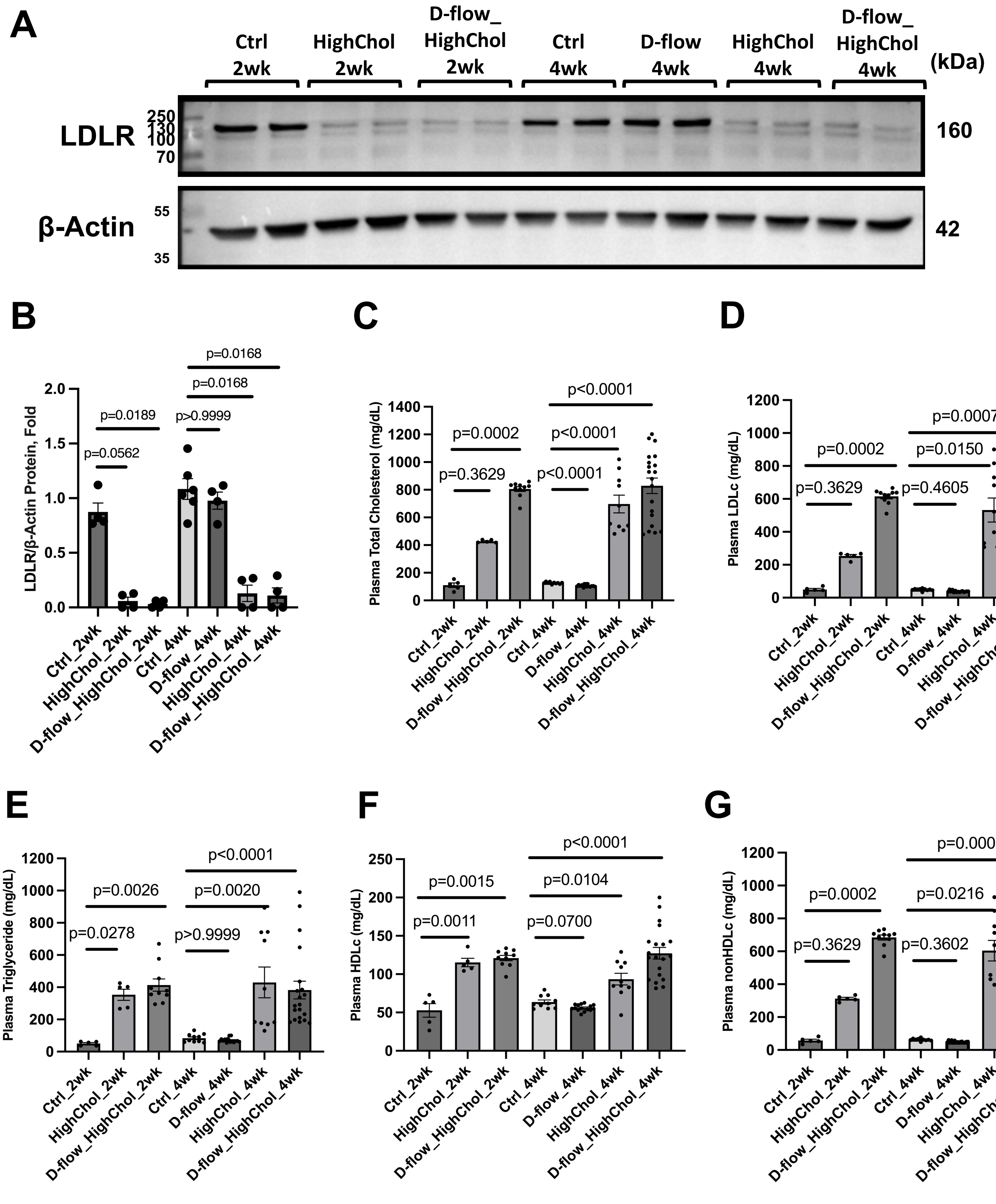


**Supplementary Figure 3. Validation of hypercholesterolemia induction in mice treated by AAV-PCSK9 and Western diet**

**A** shows representative western blots of mouse liver lysates showing the effect of LDLR knockdown by AAV8-PCSK9, as quantified by image analysis in **B** (N=4-6). Lipid analysis was performed on mouse blood plasma samples (N=5-20) (**C-G**), showing hypercholesterolemia. Quantifications are presented as mean ± SEM. *P*-values were calculated for 2 week and 4 week datasets separately, using Kruskal-Wallis one-way ANOVA for non-normal data and Brown-Forsythe ANOVA for normal data with non-equal variances.

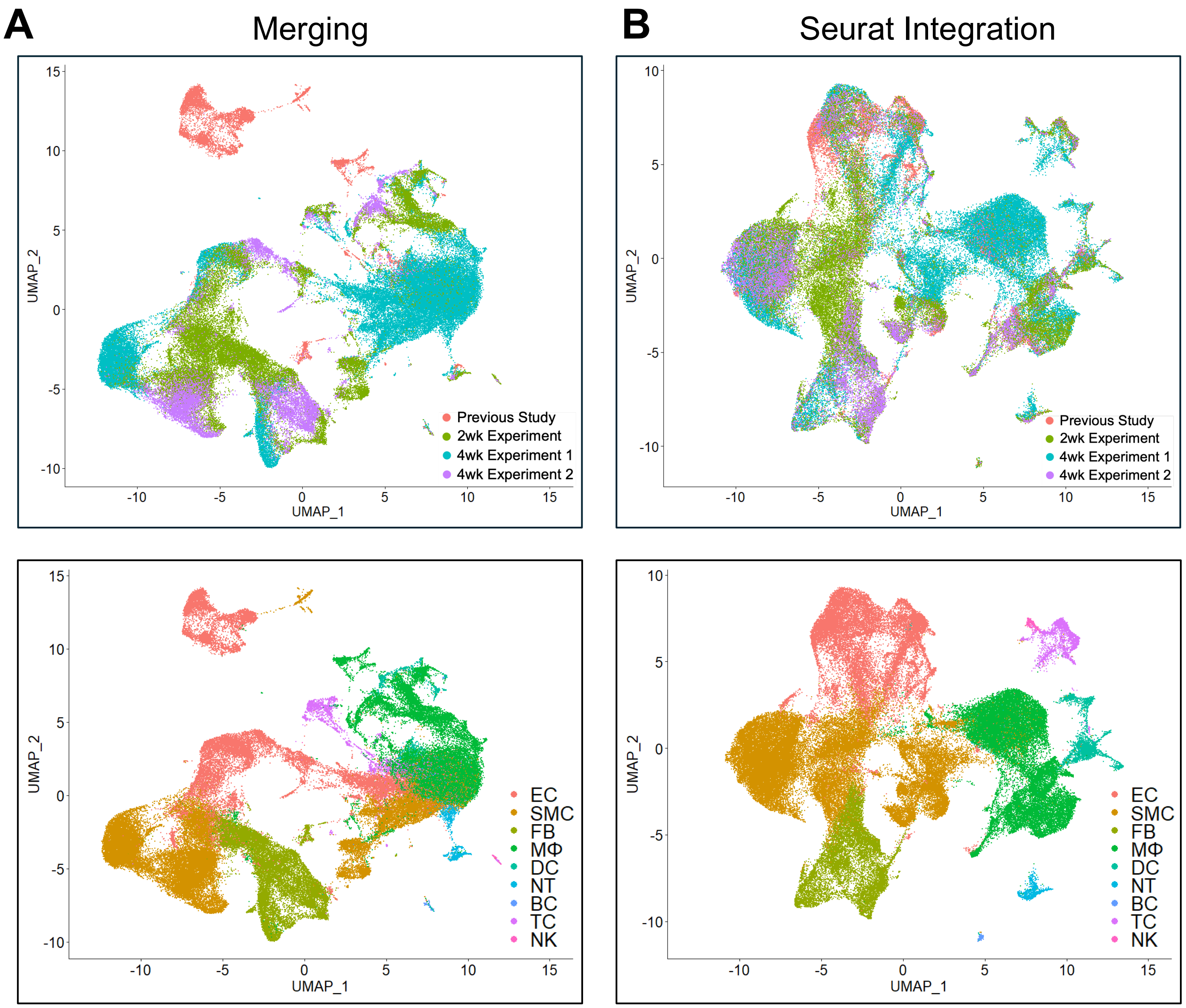


**Supplementary Figure 4. Seurat integration successfully performs batch effect correction**
UMAP plot of all single cells prepared in 4 different batches (our previous study[27], 2wk Experiment, 4wk Experiment 1, and 4wk Experiment 2) by batch at the top and by cell type at the bottom before (**A**) and after (**B**) batch effect correction using integration function in the *Seurat* R package.

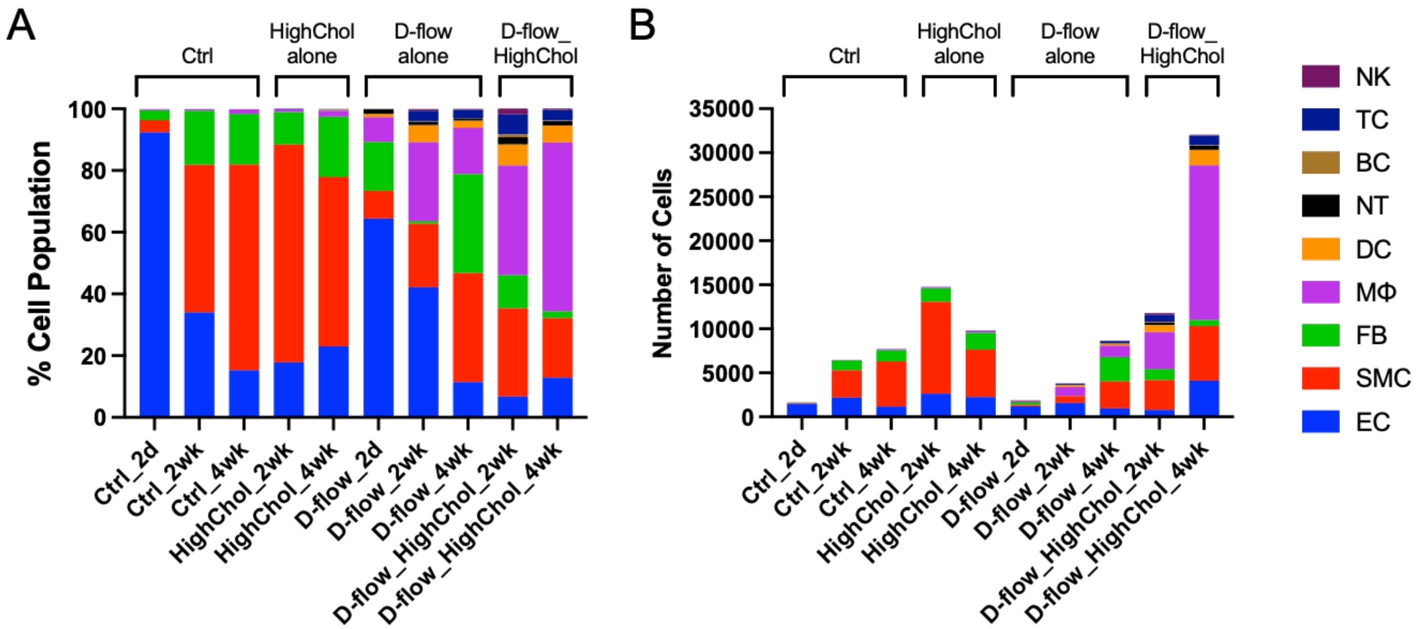


**Supplementary Figure 5. Stacked column graphs showing cell type composition for each experimental condition.** Column graphs showing **A.** % cell composition and **B.** absolute cell number of all cell types across the 10 experimental conditions (Ctrl, HighChol alone, D-flow alone, and D-flow_HighChol).

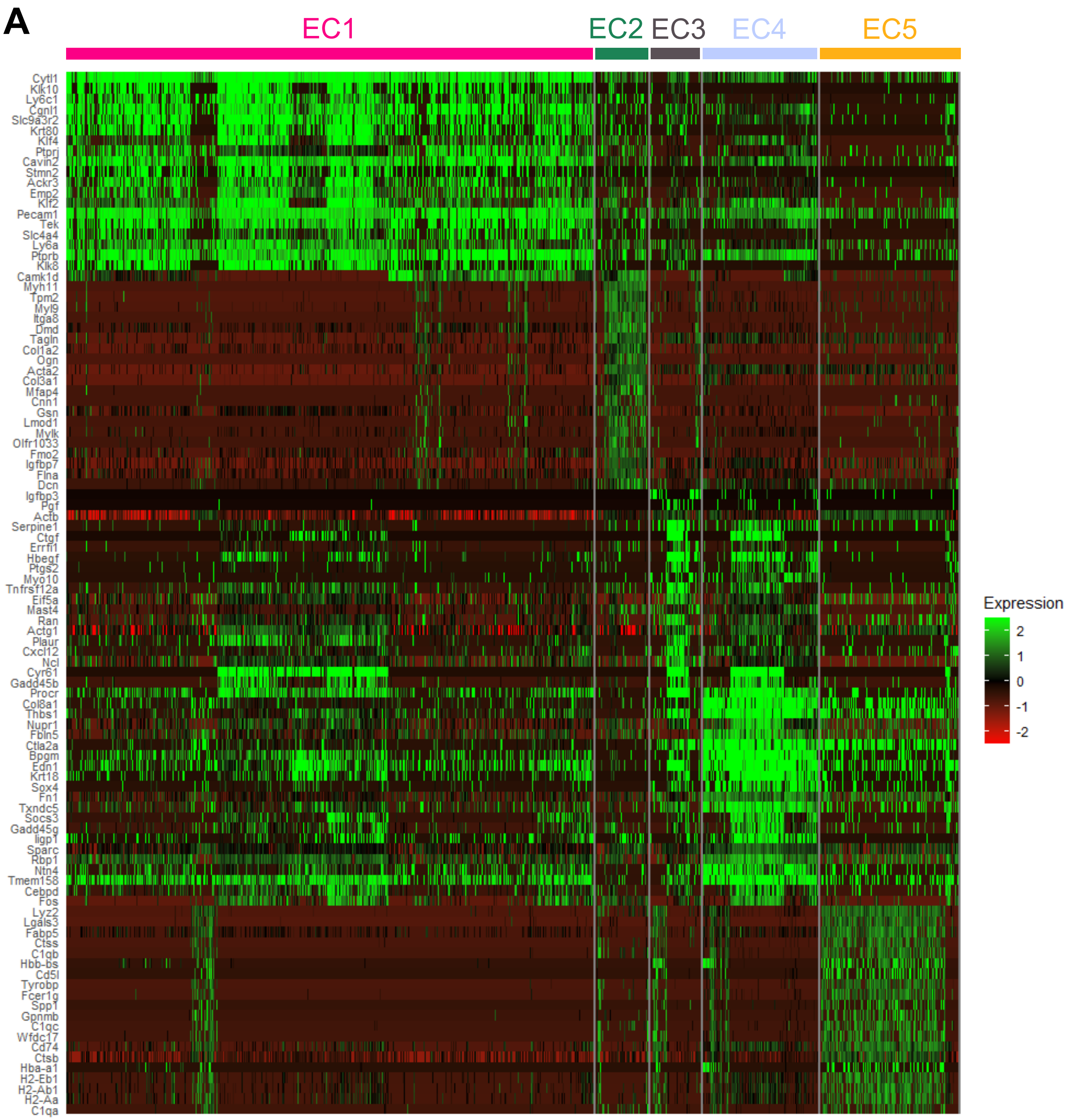
A
EC1
EC2
EC3
EC4
EC5
Cytl1
Klk10
Ly6c1
Cgnl1
Slc9a3r2
Krt80
Klf4
Ptprj
Cavin2
Stmn2
Ackr3
Emp2
Klf2
Pecam1
Tek
Slc4a4
Ly6a
Ptprb
Klk8
Camk1d
Myh11
Tpm2
Myl9
Itga8
Dmd
Tagln
Col1a2
Ogn
Acta2
Col3a1
Mfap4
Cnn1
Gsn
Lmod1
Mylk
Olfr1033
Fmo2
Igfbp7
Flna
Dcn
Igfbp3
Pgf
Actb
Serpine1
Ctgf
Errfi1
Hbegf
Ptgs2
Myo10
Tnfrsf12a
Eif5a
Mast4
Ran
Actg1
Plaur
Cxcl12
Ncl
Cyr61
Gadd45b
Procr
Col8a1
Thbs1
Nupr1
Fbln5
Ctla2a
Bpgm
Edn1
Krt18
Sox4
Fn1
Txndc5
Socs3
Gadd45g
Iigp1
Sparc
Rbp1
Ntn4
Tmem158
Cebpd
Fos
Lyz2
Lgals3
Fabp5
Ctss
C1qb
Hbb-bs
Cd5l
Tyrobp
Fcer1g
Spp1
Gpnmb
C1qc
Wfdc17
Cd74
Ctsb
Hba-a1
H2-Eb1
H2-Ab1
H2-Aa
C1qa
Expression
2
1
0
-1
-2

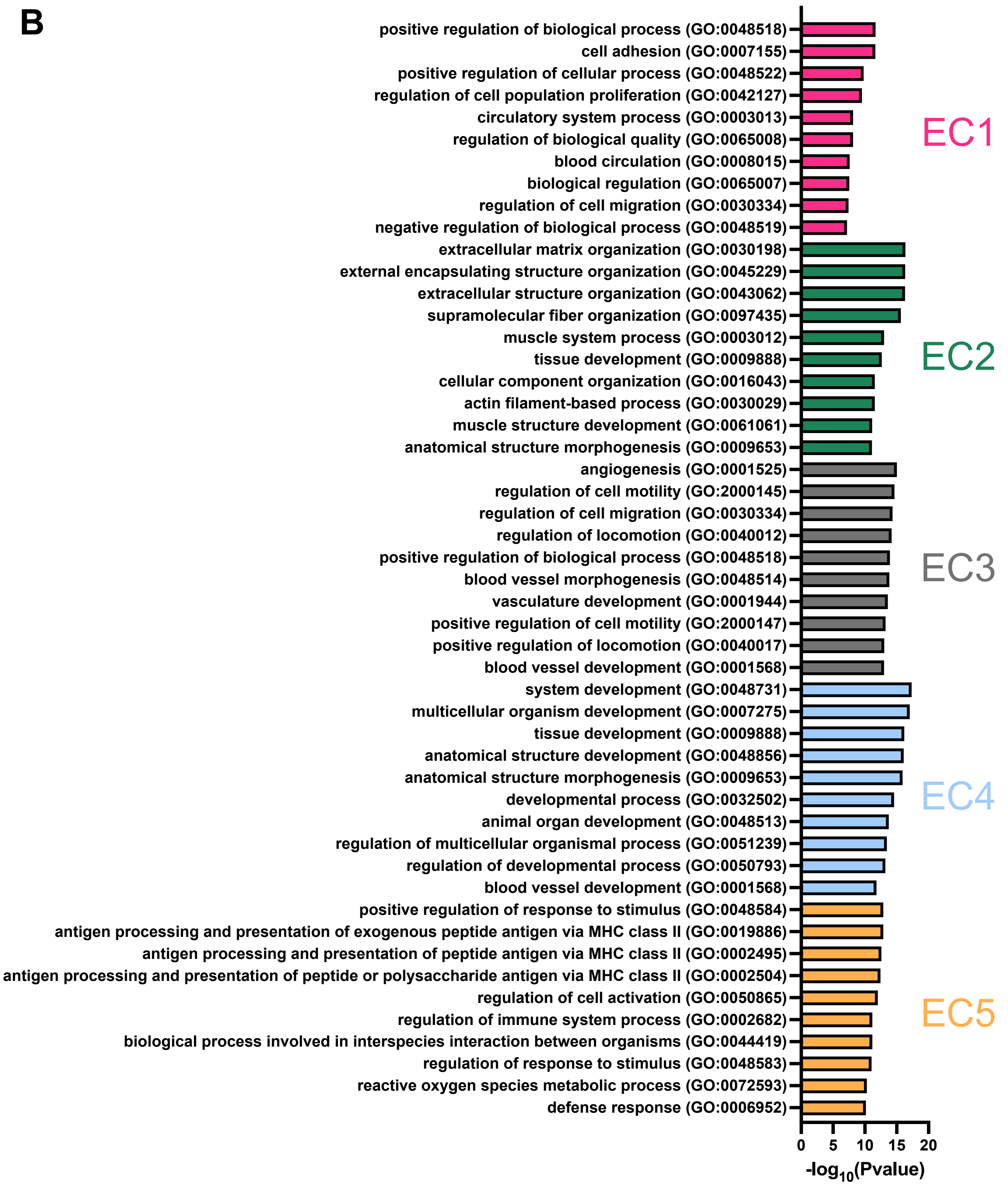


**Supplementary Figure 6. Heatmap and Gene Ontology analysis for the 5 EC clusters**
**A.** Heatmap of top 20 most highly enriched genes for each EC cluster. **B.** Top 10 Gene Ontology (GO) Biological Process (BP) terms using top 100 differentially upregulated genes for each EC cluster.

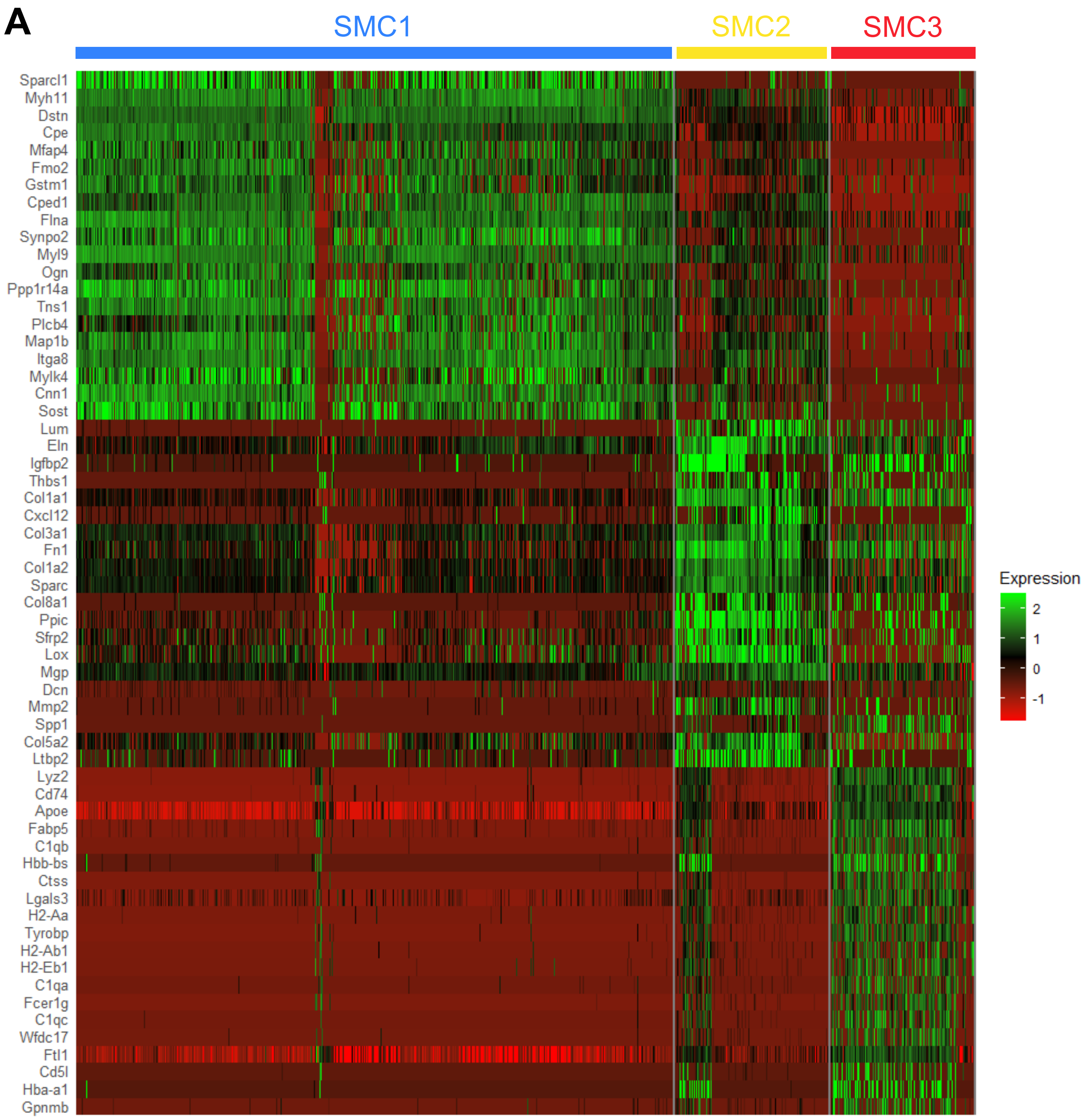
A
SMC1
SMC2
SMC3
Sparcl1
Myh11
Dstn
Cpe
Mfap4
Fmo2
Gstm1
Cped1
Flna
Synpo2
Myl9
Ogn
Ppp1r14a
Tns1
Plcb4
Map1b
Itga8
Mylk4
Cnn1
Sost
Lum
Eln
Igfbp2
Thbs1
Col1a1
Cxcl12
Col3a1
Fn1
Col1a2
Sparc
Col8a1
Ppic
Sfrp2
Lox
Mgp
Dcn
Mmp2
Spp1
Col5a2
Ltbp2
Lyz2
Cd74
Apoe
Fabp5
C1qb
Hbb-bs
Ctss
Lgals3
H2-Aa
Tyrobp
H2-Ab1
H2-Eb1
C1qa
Fcer1g
C1qc
Wfdc17
Ftl1
Cd5l
Hba-a1
Gpnmb
Expression
2
1
0
-1

B

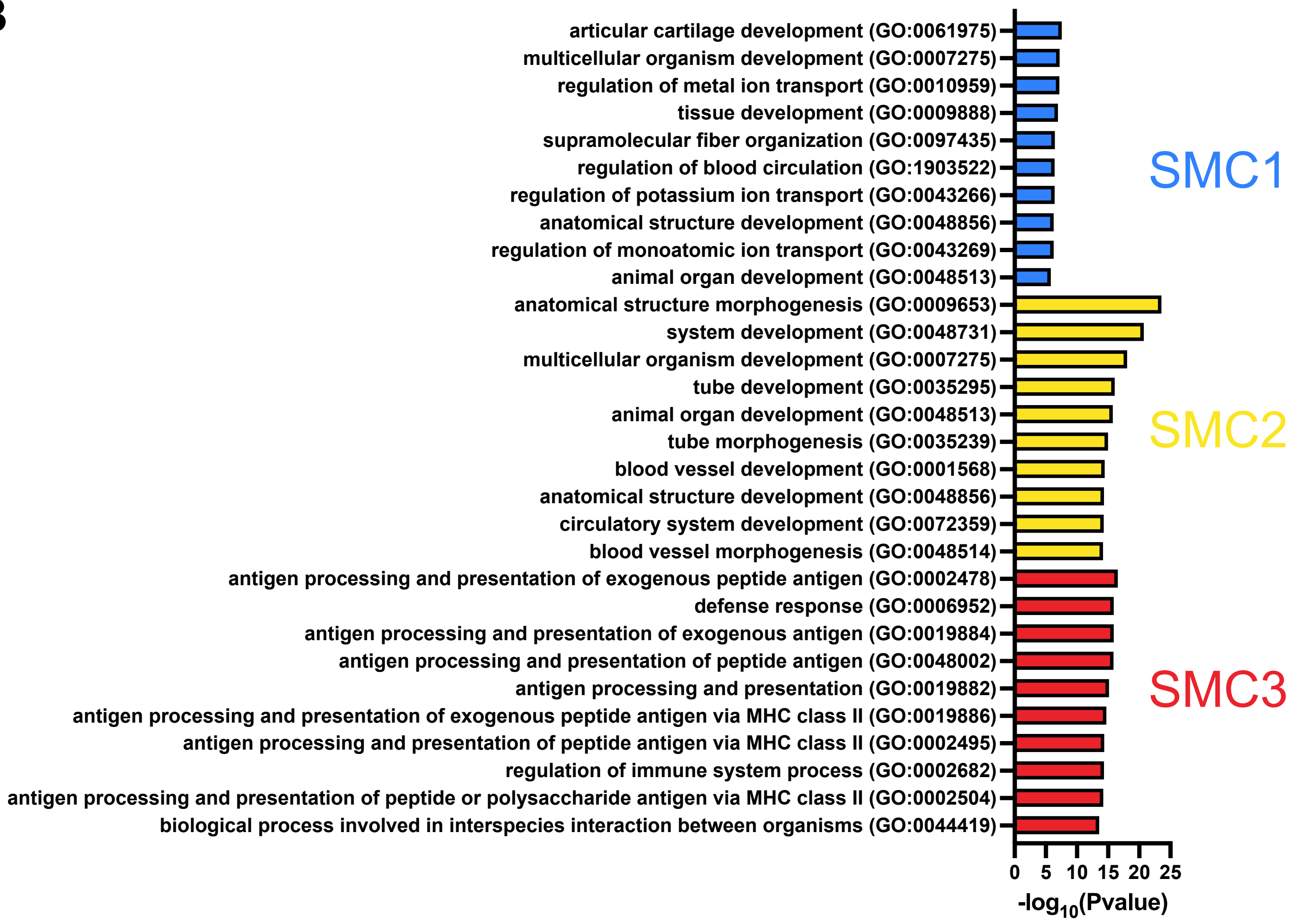


**Supplementary Figure 7. Heatmap and Gene Ontology analysis for the 3 SMC clusters**
**A.** Heatmap of top 20 most highly enriched genes for each SMC cluster. **B.** Top 10 GO BP terms using top 100 differentially upregulated genes for each SMC cluster.

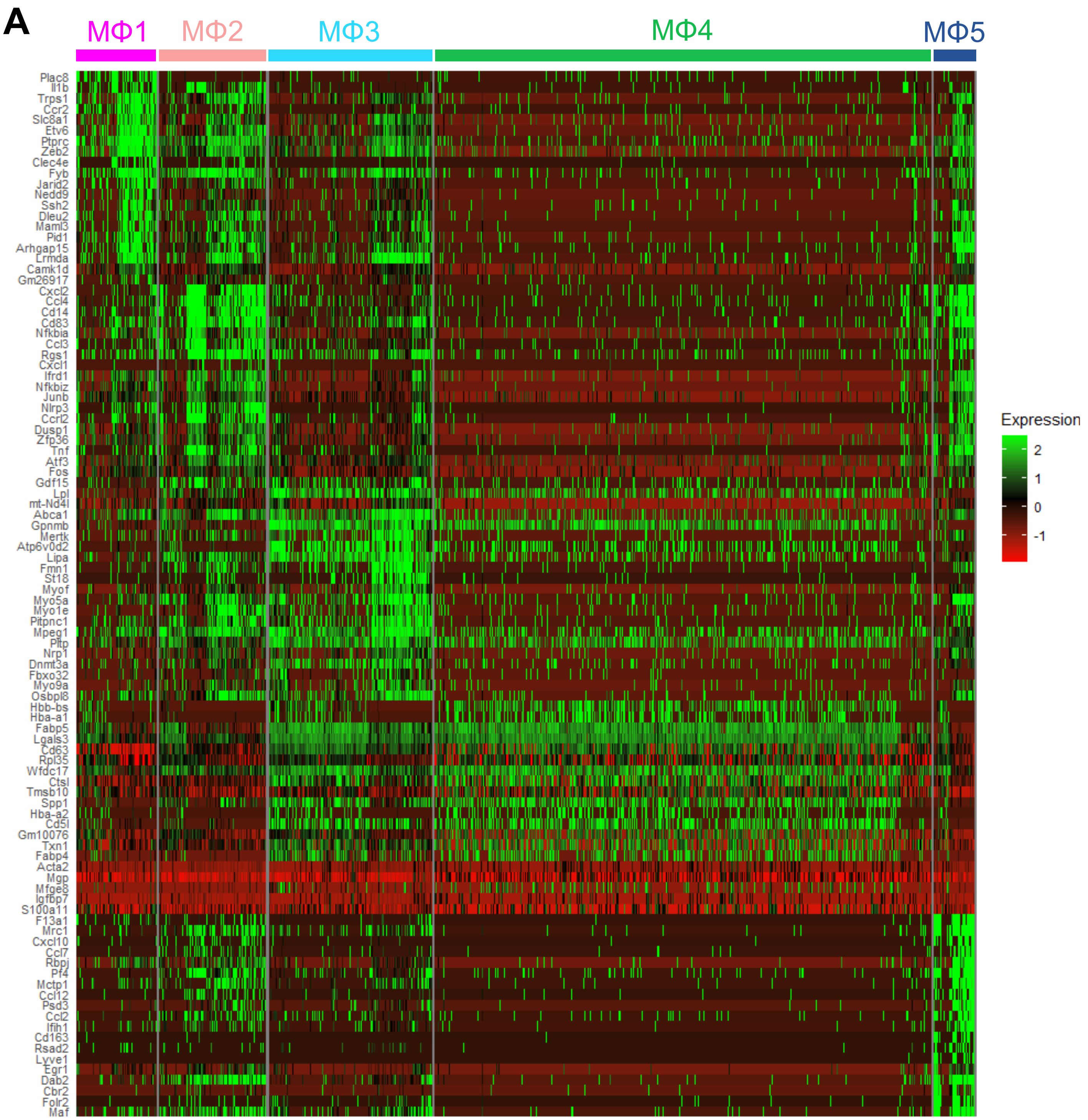
A
MΦ1
MΦ2
MΦ3
MΦ4
MΦ5
Plac8
Il1b
Trps1
Ccr2
Slc8a1
Etv6
Ptprc
Zeb2
Clec4e
Fyb
Jarid2
Nedd9
Ssh2
Dleu2
Maml3
Pid1
Arhgap15
Lrmda
Camk1d
Gm26917
Cxcl2
Ccl4
Cd14
Cd83
Nfkbia
Ccl3
Rgs1
Cxcl1
Ifrd1
Nfkbiz
Junb
Nlrp3
Ccrl2
Dusp1
Zfp36
Tnf
Atf3
Fos
Gdf15
Lpl
mt-Nd4l
Abca1
Gpnmb
Mertk
Atp6v0d2
Lipa
Fmn1
St18
Myof
Myo5a
Myo1e
Pitpnc1
Mpeg1
Pltp
Nrp1
Dnmt3a
Fbxo32
Myo9a
Osbpl8
Hbb-bs
Hba-a1
Fabp5
Lgals3
Cd63
Rpl35
Wfdc17
Ctsl
Tmsb10
Spp1
Hba-a2
Cd5l
Gm10076
Txn1
Fabp4
Acta2
Mgp
Mfge8
Igfbp7
S100a11
F13a1
Mrc1
Cxcl10
Ccl7
Rbpj
Pf4
Mctp1
Ccl12
Psd3
Ccl2
Ifih1
Cd163
Rsad2
Lyve1
Egr1
Dab2
Cbr2
Folr2
Maf
Expression
2
1
0
-1

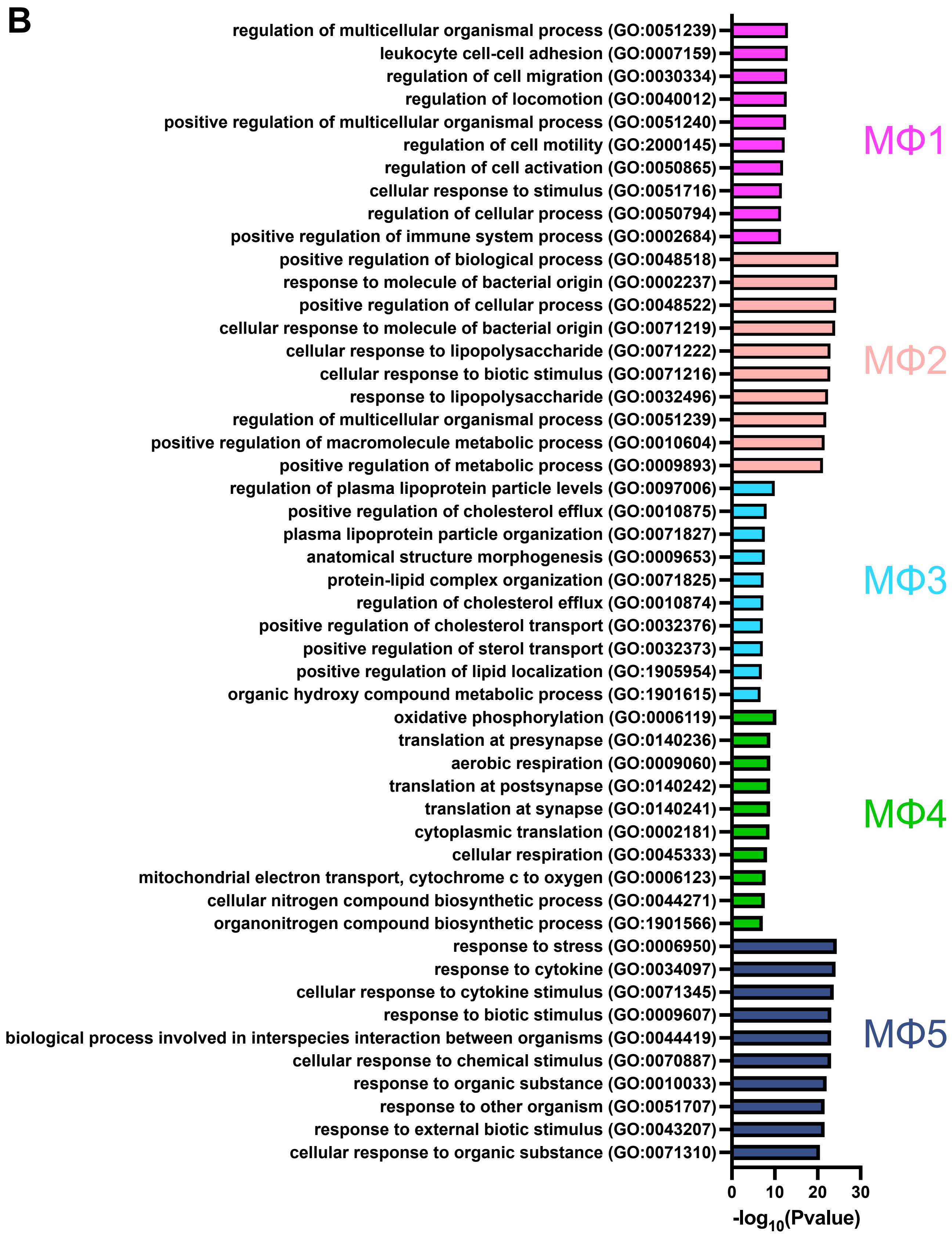


**Supplementary Figure 8. Heatmap and Gene Ontology analysis for the 5 MΦ clusters**
**A.** Heatmap of top 20 most highly enriched genes for each MΦ cluster. **B.** Top 10 GO BP terms using top 100 differentially upregulated genes for each MΦ cluster.

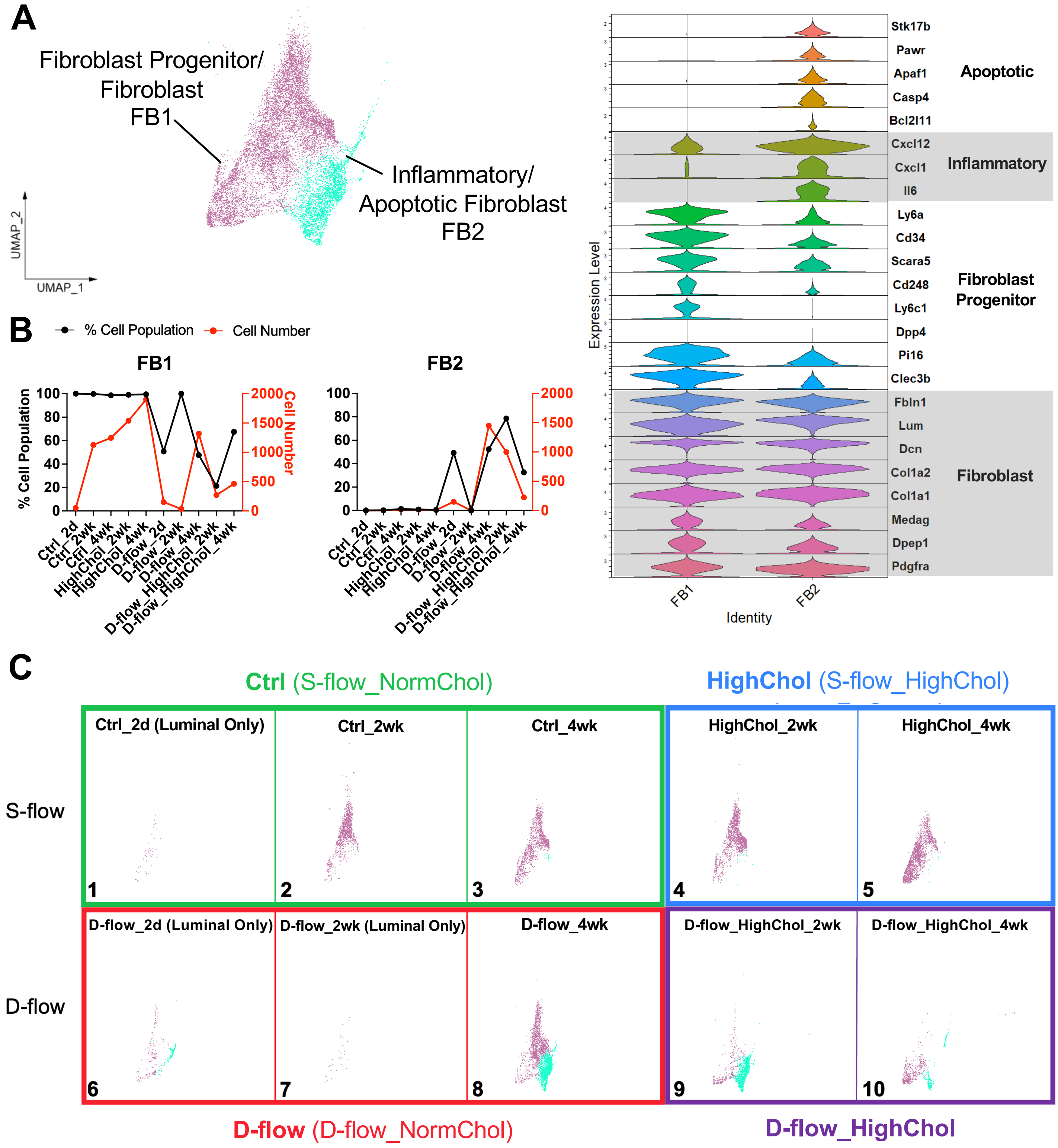
A
Fibroblast Progenitor/ Fibroblast FB1
Inflammatory/ Apoptotic Fibroblast FB2
UMAP_2
UMAP_1
Stk17b
Pawr
Apaf1
Casp4
Bcl2l11
Apoptotic
Cxcl12
Cxcl1
Il6
Inflammatory
Ly6a
Cd34
Scara5
Cd248
Ly6c1
Dpp4
Pi16
Clec3b
Fibroblast Progenitor
Fbln1
Lum
Dcn
Col1a2
Col1a1
Medag
Dpep1
Pdgfra
Fibroblast
Expression Level
FB1
FB2
Identity
B
% Cell Population
Cell Number
FB1
FB2
100
80
60
40
20
0
2000
1500
1000
500
0
Ctrl_2d
Ctrl_2wk
Ctrl_4wk
HighChol_2wk
HighChol_4wk
D-flow_2d
D-flow_2wk
D-flow_4wk
D-flow_HighChol_2wk
D-flow_HighChol_4wk
C
Ctrl (S-flow_NormChol)
HighChol (S-flow_HighChol)
S-flow
Ctrl_2d (Luminal Only)
Ctrl_2wk
Ctrl_4wk
HighChol_2wk
HighChol_4wk
1
2
3
4
5
D-flow
D-flow_2d (Luminal Only)
D-flow_2wk (Luminal Only)
D-flow_4wk
D-flow_HighChol_2wk
D-flow_HighChol_4wk
6
7
8
9
10
D-flow (D-flow_NormChol)
D-flow_HighChol

D

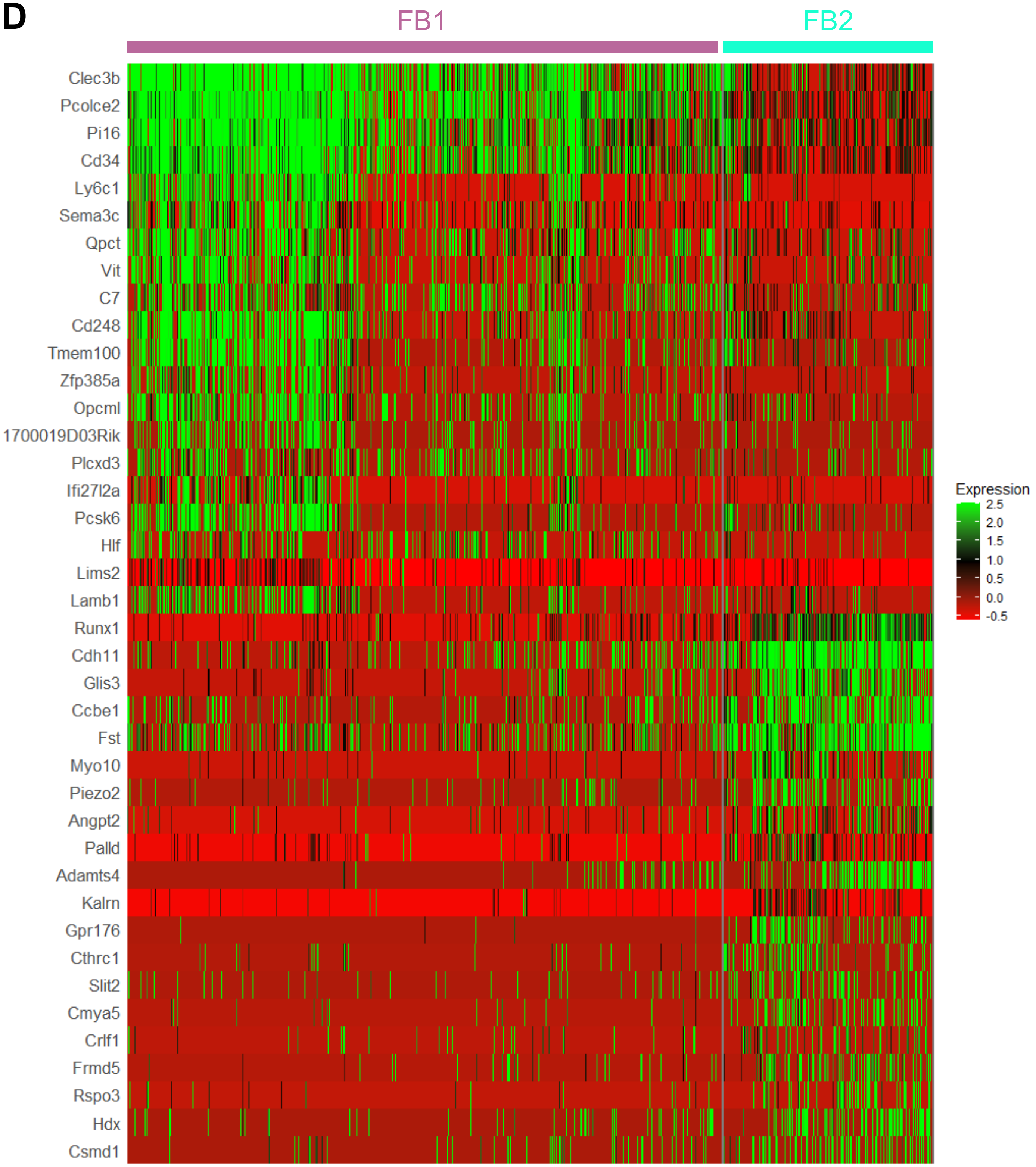
FB1
FB2
Clec3b
Pcolce2
Pi16
Cd34
Ly6c1
Sema3c
Qpct
Vit
C7
Cd248
Tmem100
Zfp385a
Opcml
1700019D03Rik
Plcxd3
Ifi27l2a
Pcsk6
Hlf
Lims2
Lamb1
Runx1
Cdh11
Glis3
Ccbe1
Fst
Myo10
Piezo2
Angpt2
Palld
Adamts4
Kalrn
Gpr176
Cthrc1
Slit2
Cmya5
Crlf1
Frmd5
Rspo3
Hdx
Csmd1
Expression
2.5
2.0
1.5
1.0
0.5
0.0
-0.5

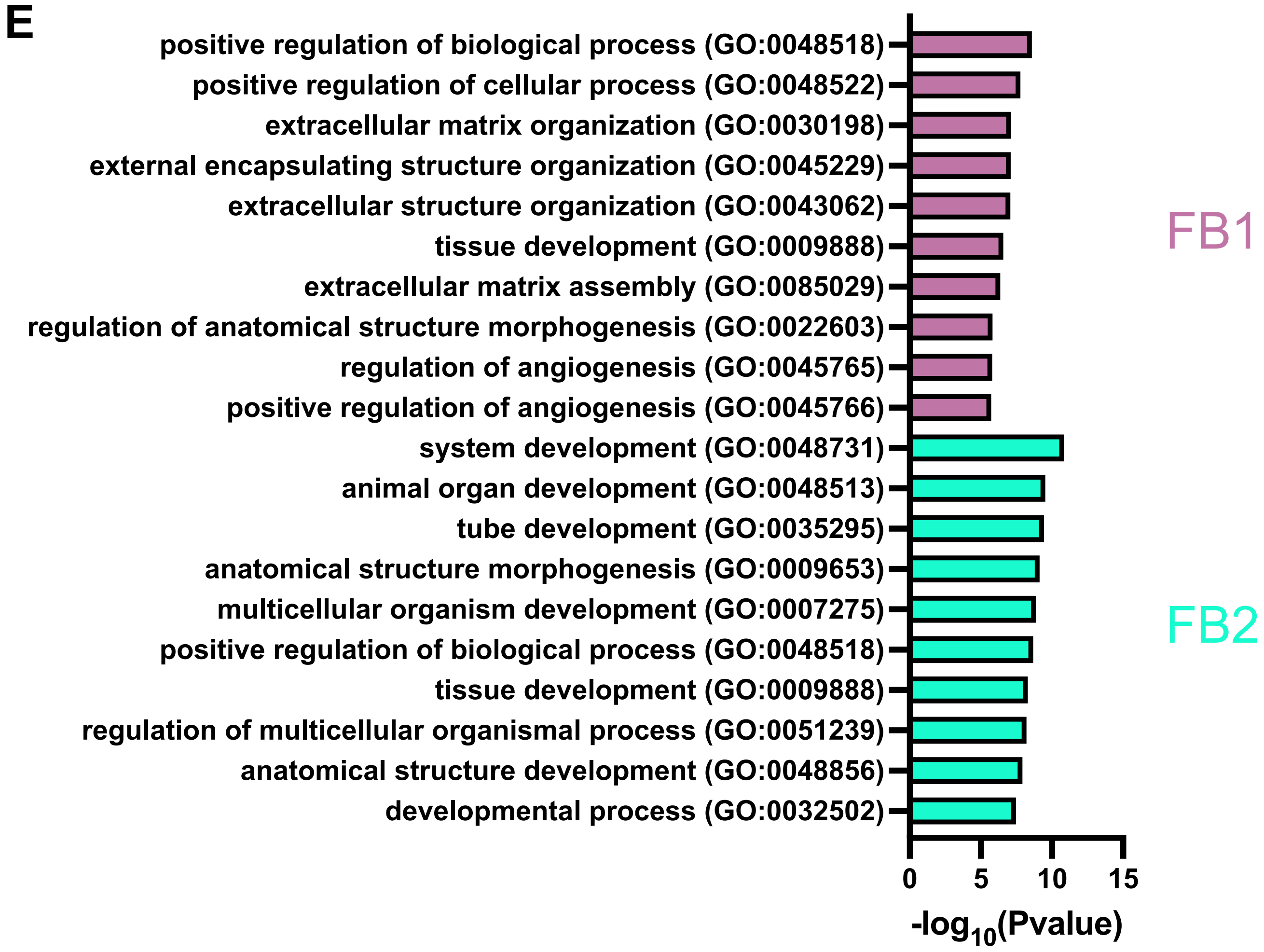


**Supplementary Figure 9. D-flow increases inflammatory and apoptotic fibroblast population**
**A**. UMAP plot of 2 FB clusters (10,957 cells in total) and stacked violin plot shows expression levels of genes used to annotate each FB cluster. FB clusters include fibroblast progenitor/fibroblast FB1 and inflammatory/apoptotic fibroblast FB2. **B**. Cell number and % cell population (cell number for each FB cluster normalized by the total number of FBs per group) quantifications for each FB cluster across all 10 experimental groups. **C**. UMAP plot for each experimental group is shown. N= 5-20 mice for each condition. **D.** Heatmap of top 20 most highly enriched genes for each FB cluster. **E.** Top 10 GO BP terms using top 100 differentially upregulated genes for each FB cluster.

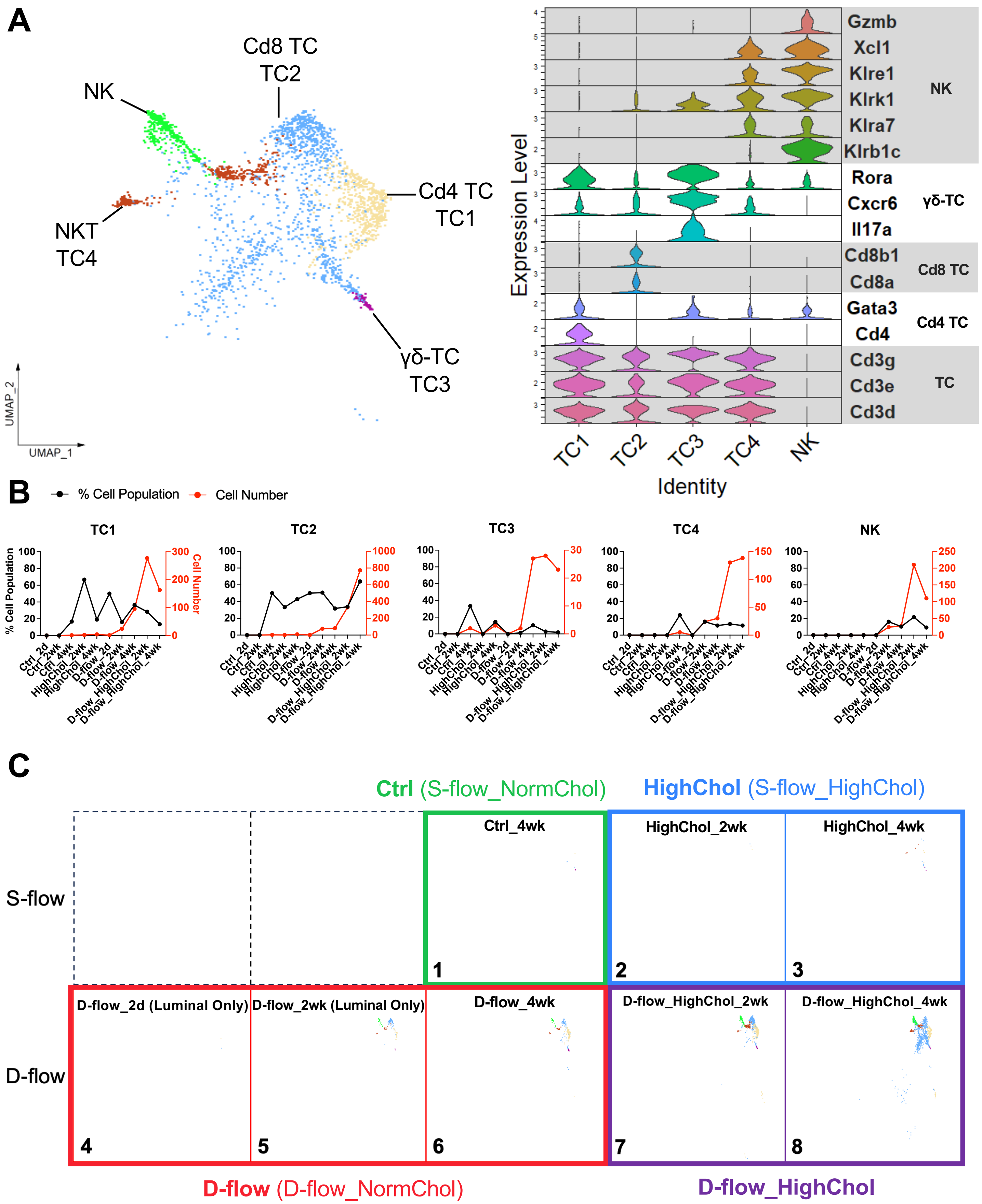
A
NK
Cd8 TC
TC2
Cd4 TC
TC1
NKT
TC4
γδ-TC
TC3
UMAP_1
UMAP_2
Gzmb
Xcl1
Klre1
Klrk1
Klra7
Klrb1c
Rora
Cxcr6
Il17a
Cd8b1
Cd8a
Gata3
Cd4
Cd3g
Cd3e
Cd3d
NK
γδ-TC
Cd8 TC
Cd4 TC
TC
Expression Level
TC1 TC2 TC3 TC4 NK
Identity
B
% Cell Population
Cell Number
TC1
TC2
TC3
TC4
NK
% Cell Population
Cell Number
Ctrl_2d
Ctrl_2wk
Ctrl_4wk
HighChol_2wk
HighChol_4wk
D-flow_2d
D-flow_2wk
D-flow_4wk
D-flow_HighChol_2wk
D-flow_HighChol_4wk
C
Ctrl (S-flow_NormChol)
HighChol (S-flow_HighChol)
S-flow
D-flow
Ctrl_4wk
HighChol_2wk
HighChol_4wk
D-flow_2d (Luminal Only)
D-flow_2wk (Luminal Only)
D-flow_4wk
D-flow_HighChol_2wk
D-flow_HighChol_4wk
1
2
3
4
5
6
7
8
D-flow (D-flow_NormChol)
D-flow_HighChol

D

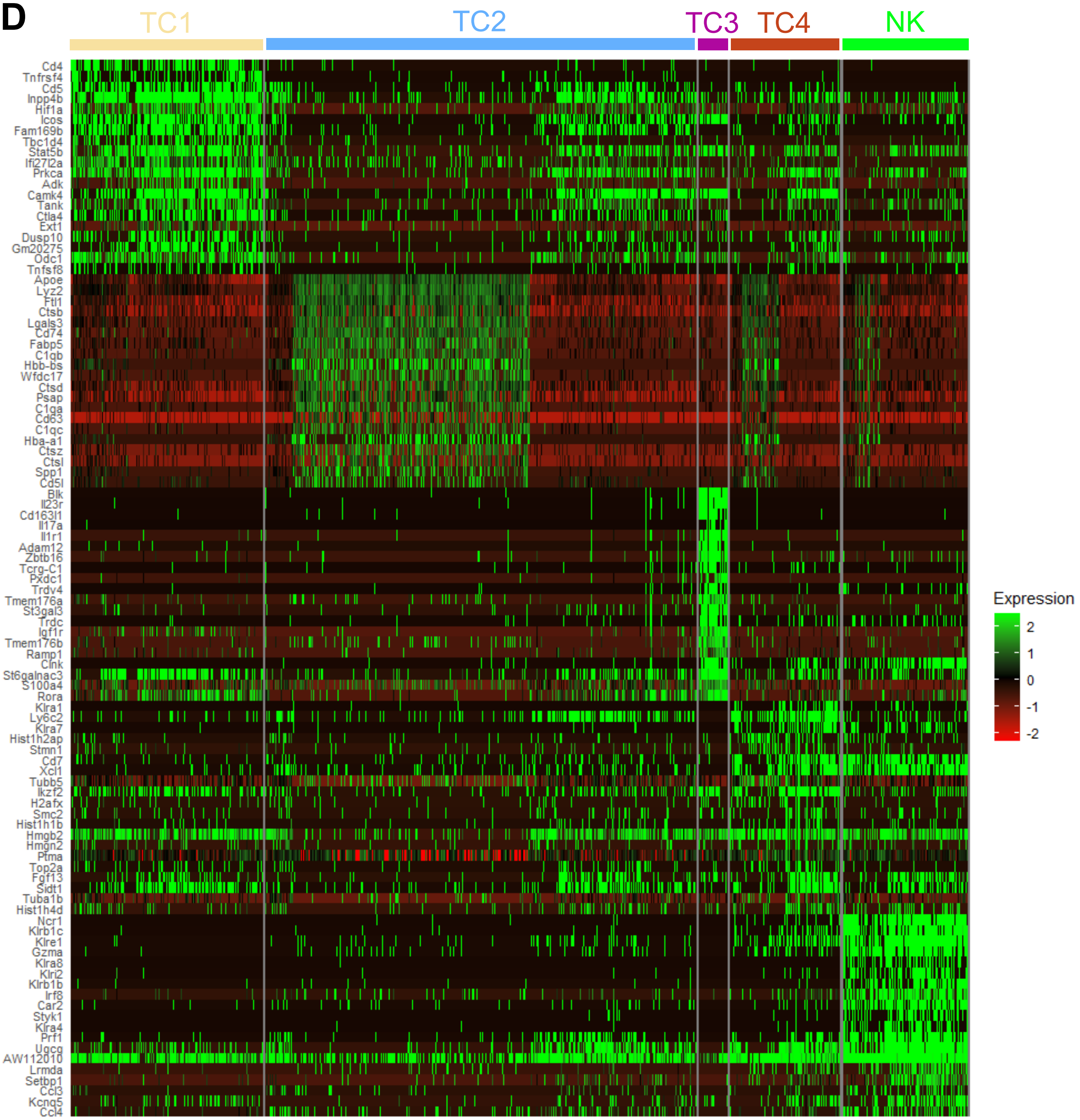
TC1
TC2
TC3
TC4
NK
Cd4
Tnfrsf4
Cd5
Inpp4b
Hif1a
Icos
Fam169b
Tbc1d4
Stat5b
Ifi27l2a
Prkca
Adk
Camk4
Tank
Ctla4
Ext1
Dusp10
Gm20275
Odc1
Tnfsf8
Apoe
Lyz2
Ftl1
Ctsb
Lgals3
Cd74
Fabp5
C1qb
Hbb-bs
Wfdc17
Ctsd
Psap
C1qa
Cd63
C1qc
Hba-a1
Ctsz
Ctsl
Spp1
Cd5l
Blk
Il23r
Cd163l1
Il17a
Il1r1
Adam12
Zbtb16
Tcrg-C1
Pxdc1
Trdv4
Tmem176a
St3gal3
Trdc
Igf1r
Tmem176b
Ramp1
Clnk
St6galnac3
S100a4
Rora
Klra1
Ly6c2
Klra7
Hist1h2ap
Stmn1
Cd7
Xcl1
Tubb5
Ikzf2
H2afx
Smc2
Hist1h1b
Hmgb2
Hmgn2
Ptma
Top2a
Fgf13
Sidt1
Tuba1b
Hist1h4d
Ncr1
Klrb1c
Klre1
Gzma
Klra8
Klri2
Klrb1b
Irf8
Car2
Styk1
Klra4
Prf1
Ugcg
AW112010
Lrmda
Setbp1
Ccl3
Kcnq5
Ccl4
Expression
2
1
0
-1
-2

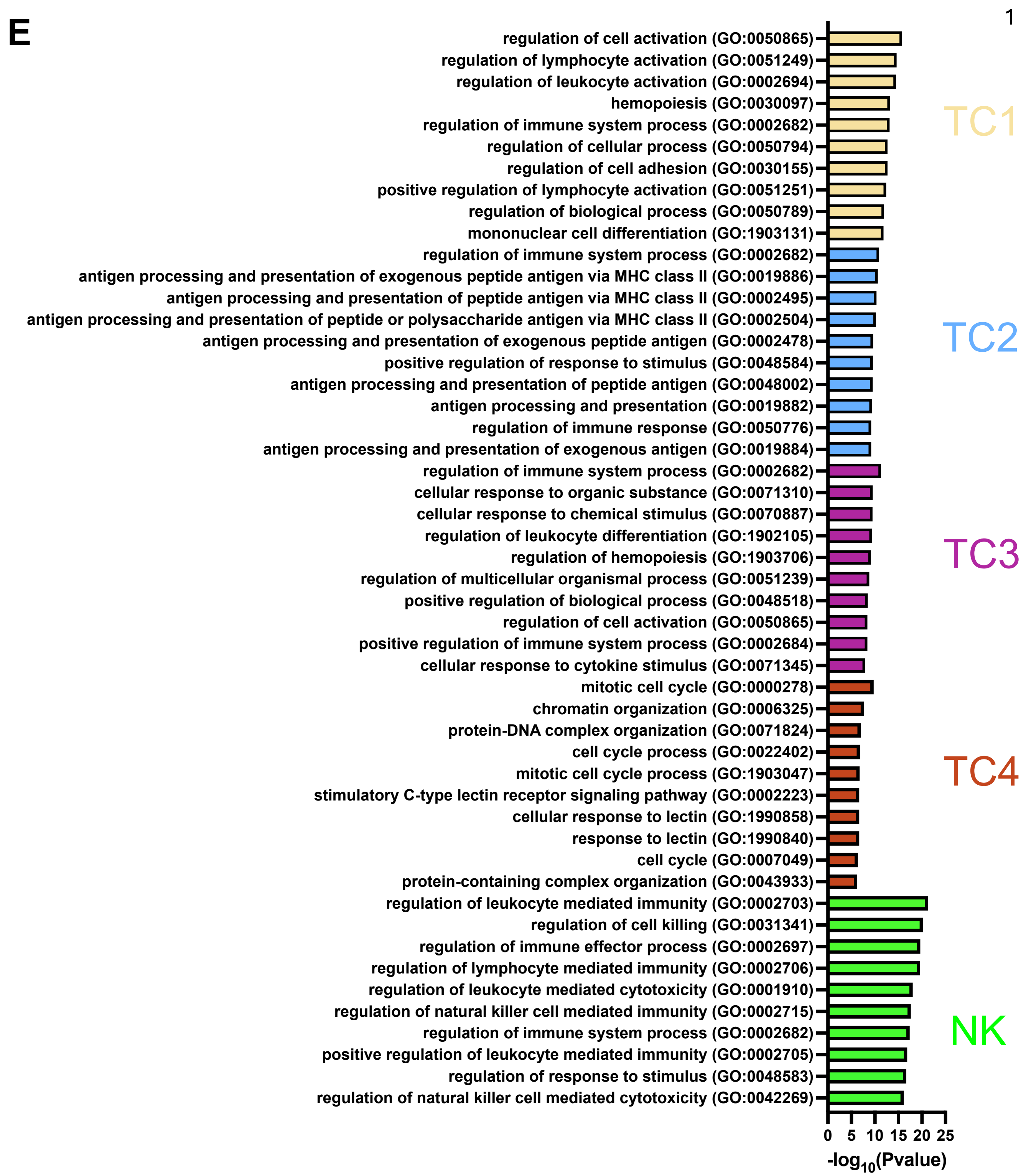


**Supplementary Figure 10. D-flow induces T cell infiltration, especially the Cd8 T cells, and exacerbated by hypercholesterolemia**

**A**. UMAP plot of 4 TC and 1 NK clusters (2,627 cells in total) and stacked violin plot shows expression levels of genes used to annotate each TC/NK cluster. TC clusters include Cd4 TC1, Cd8 TC2, γδ-T TC3, and NKT TC4. **B**. % cell population and cell number for each TC/NK cluster across all 10 experimental groups. **C**. UMAP plot for each experimental group is shown. N= 5-20 mice for each condition. **D.** Heatmap of top 20 most highly enriched genes for each TC and NK cluster. **E.** Top 10 GO BP terms using top 100 differentially upregulated genes for each TC and NK cluster.

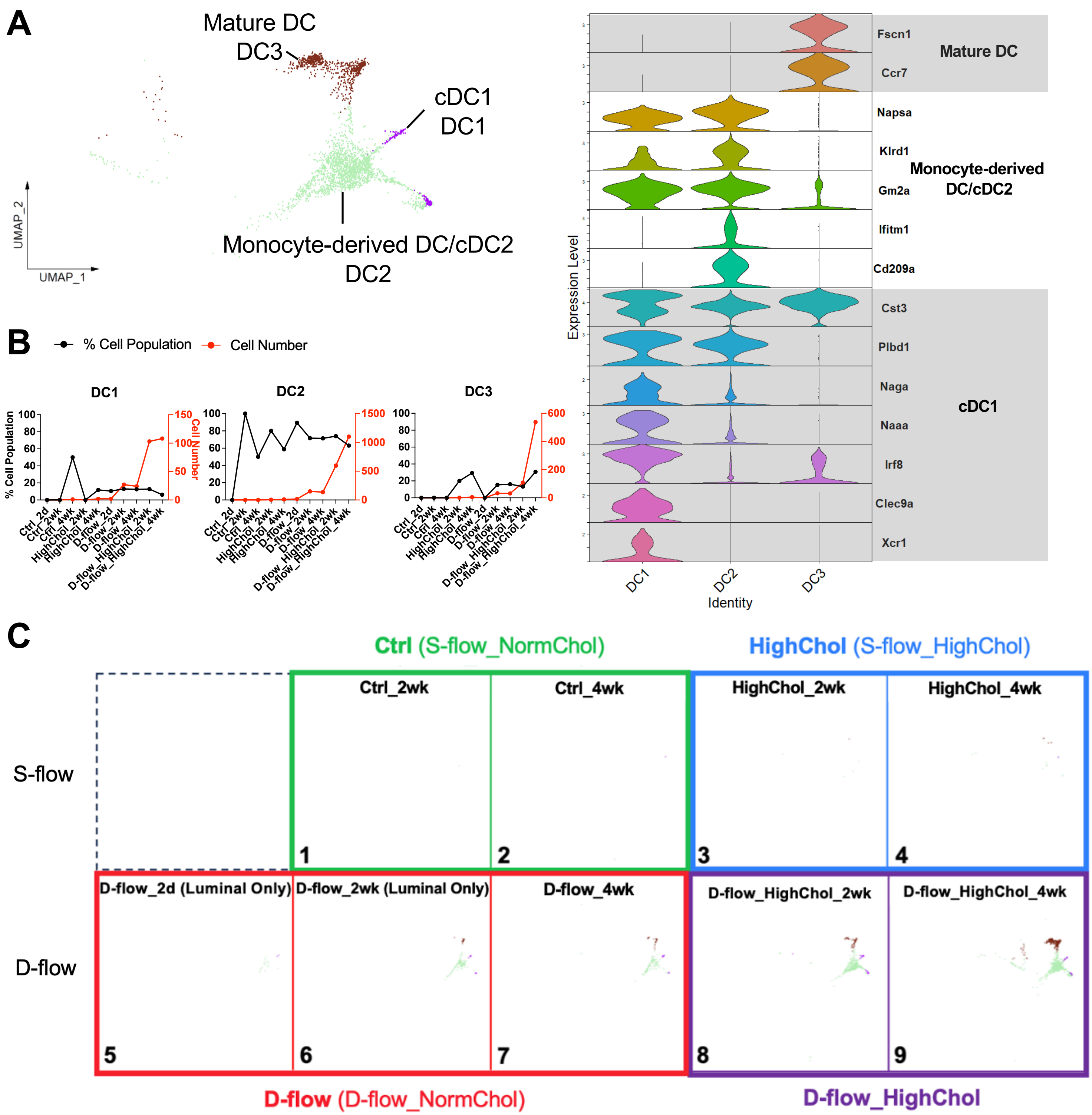
A
Mature DC
DC3
cDC1
DC1
Monocyte-derived DC/cDC2
DC2
UMAP_2
UMAP_1
Fscn1
Ccr7
Mature DC
Napsa
Klrd1
Gm2a
Ifitm1
Cd209a
Monocyte-derived DC/cDC2
Cst3
Plbd1
Naga
Naaa
Irf8
Clec9a
Xcr1
cDC1
Expression Level
DC1
DC2
DC3
Identity
B
% Cell Population
Cell Number
DC1
DC2
DC3
% Cell Population
Cell Number
Ctrl_2d
Ctrl_2wk
Ctrl_4wk
HighChol_2wk
HighChol_4wk
D-flow_2d
D-flow_2wk
D-flow_4wk
D-flow_HighChol_2wk
D-flow_HighChol_4wk
C
Ctrl (S-flow_NormChol)
HighChol (S-flow_HighChol)
S-flow
D-flow
Ctrl_2wk
Ctrl_4wk
HighChol_2wk
HighChol_4wk
D-flow_2d (Luminal Only)
D-flow_2wk (Luminal Only)
D-flow_4wk
D-flow_HighChol_2wk
D-flow_HighChol_4wk
D-flow (D-flow_NormChol)
D-flow_HighChol

D

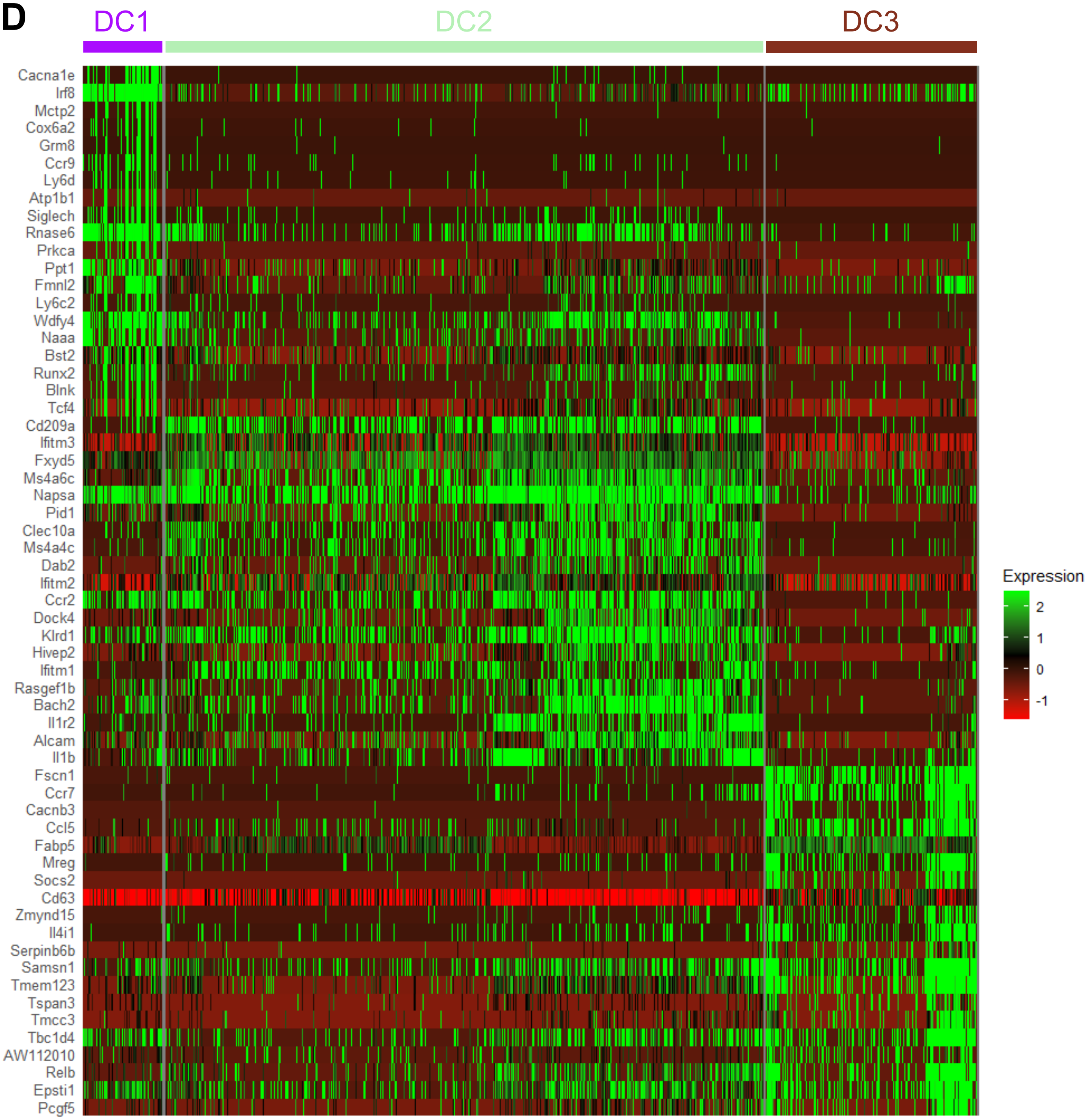
DC1
DC2
DC3
Cacna1e
Irf8
Mctp2
Cox6a2
Grm8
Ccr9
Ly6d
Atp1b1
Siglech
Rnase6
Prkca
Ppt1
Fmnl2
Ly6c2
Wdfy4
Naaa
Bst2
Runx2
Blnk
Tcf4
Cd209a
Ifitm3
Fxyd5
Ms4a6c
Napsa
Pid1
Clec10a
Ms4a4c
Dab2
Ifitm2
Ccr2
Dock4
Klrd1
Hivep2
Ifitm1
Rasgef1b
Bach2
Il1r2
Alcam
Il1b
Fscn1
Ccr7
Cacnb3
Ccl5
Fabp5
Mreg
Socs2
Cd63
Zmynd15
Il4i1
Serpinb6b
Samsn1
Tmem123
Tspan3
Tmcc3
Tbc1d4
AW112010
Relb
Epsti1
Pcgf5
Expression
2
1
0
-1

**E**

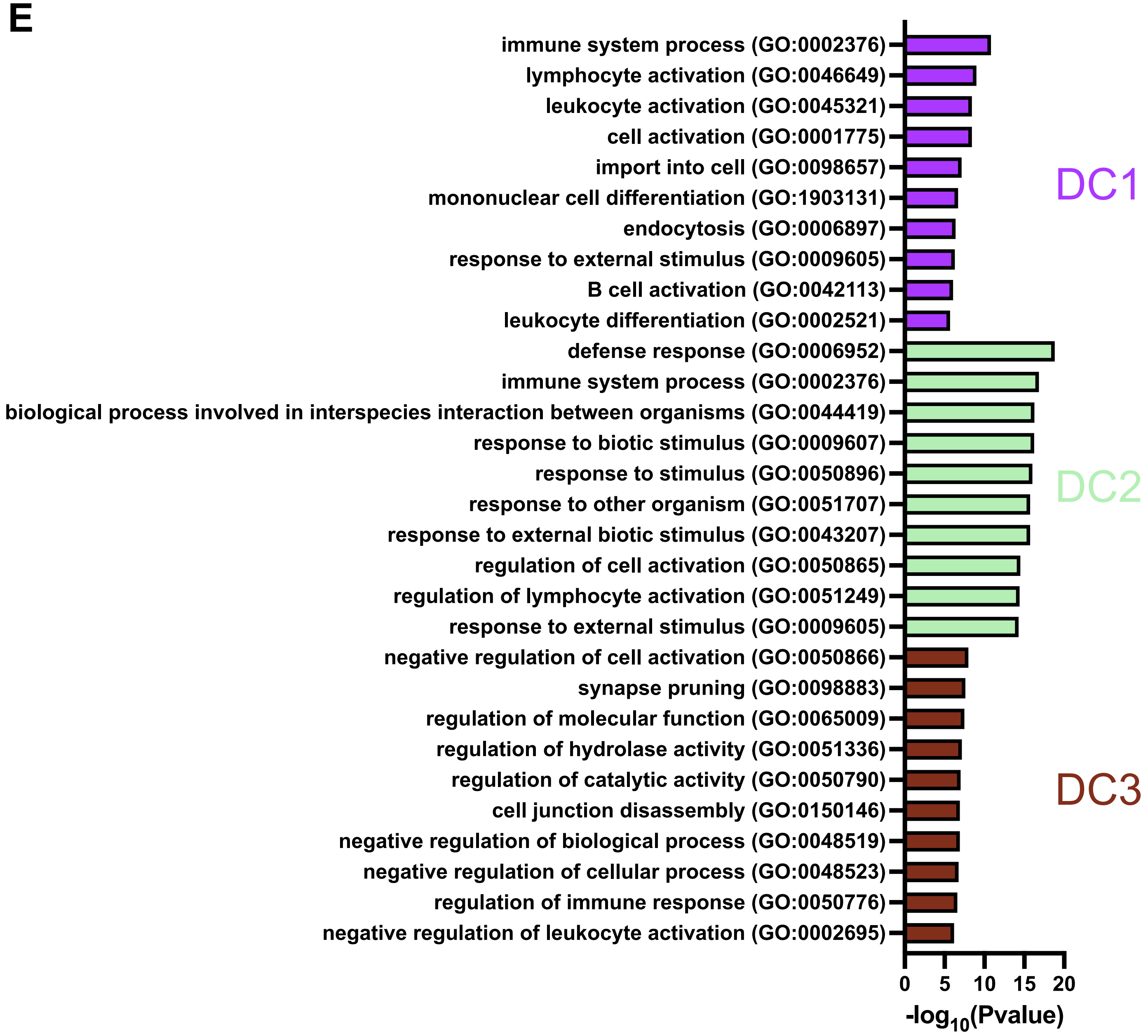


**Supplementary Figure 11. D-flow induces dendritic cell infiltration, especially the monocyte-derived DC and cDC2, and exacerbated by hypercholesterolemia**

**A**. UMAP plot of 3 DC clusters (2,997 cells in total) and stacked violin plot showing expression levels of genes used to annotate each DC cluster. DC clusters include cDC1 DC1, monocyte-derived DC/cDC2 DC2, and mature DC DC3. **B**. Cell number and % cell population (cell number for each DC cluster normalized by the total number of DCs per group) quantifications for each DC cluster across all 10 experimental groups. **C**. UMAP plot for each experimental group is shown. N= 5-20 mice for each condition. **D.** Heatmap of top 20 most highly enriched genes for each DC cluster. **E.** Top 10 GO BP terms using top 100 differentially upregulated genes for each DC cluster.

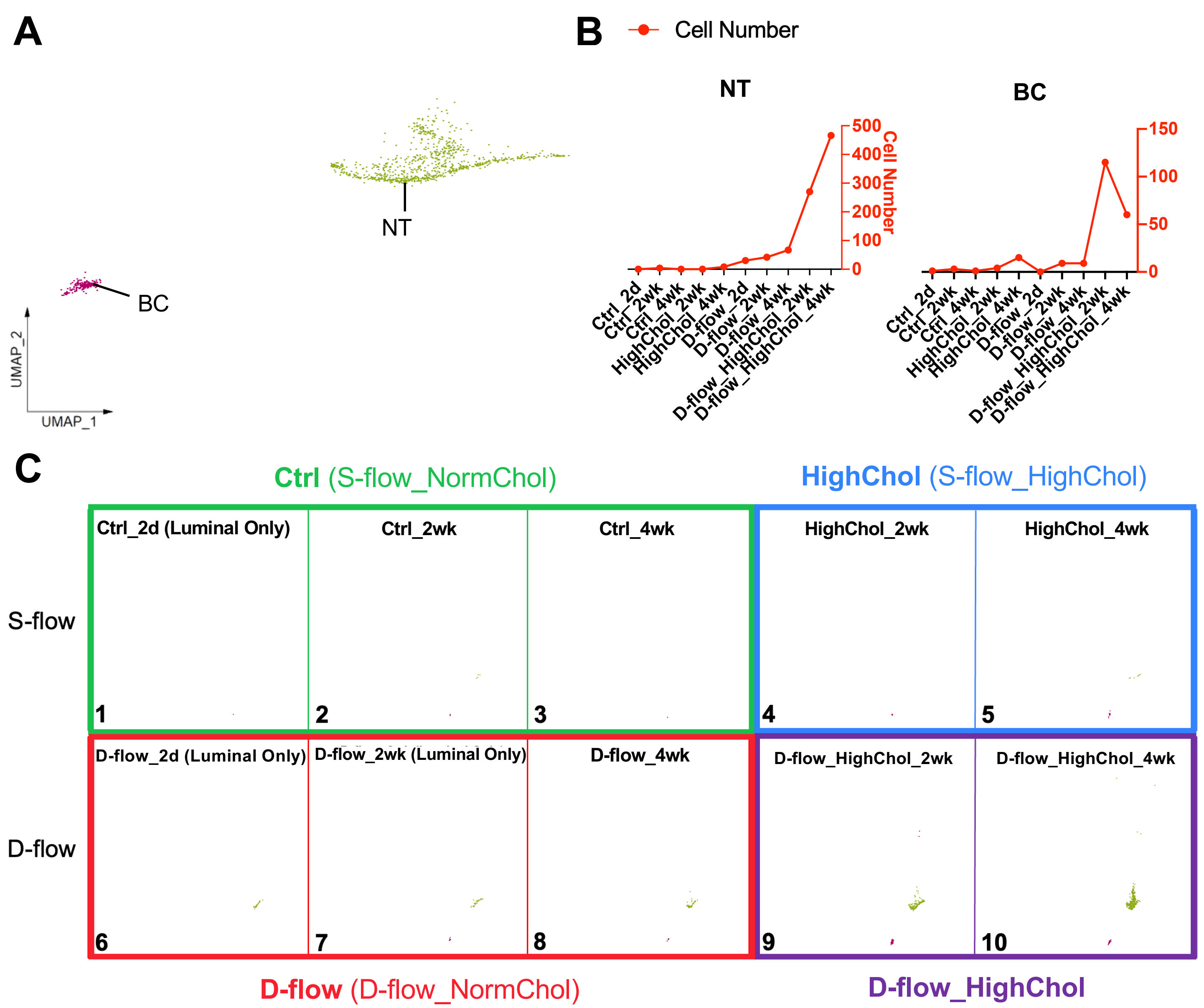

A
NT
BC
UMAP_2
UMAP_1
B
Cell Number
NT
BC
Cell Number
500
400
300
200
100
0
150
100
50
0
Ctrl_2d
Ctrl_2wk
Ctrl_4wk
HighChol_2wk
HighChol_4wk
D-flow_2d
D-flow_2wk
D-flow_4wk
D-flow_HighChol_2wk
D-flow_HighChol_4wk
C
Ctrl (S-flow_NormChol)
HighChol (S-flow_HighChol)
S-flow
D-flow
Ctrl_2d (Luminal Only)
Ctrl_2wk
Ctrl_4wk
HighChol_2wk
HighChol_4wk
D-flow_2d (Luminal Only)
D-flow_2wk (Luminal Only)
D-flow_4wk
D-flow_HighChol_2wk
D-flow_HighChol_4wk
1
2
3
4
5
6
7
8
9
10
D-flow (D-flow_NormChol)
D-flow_HighChol


1

D

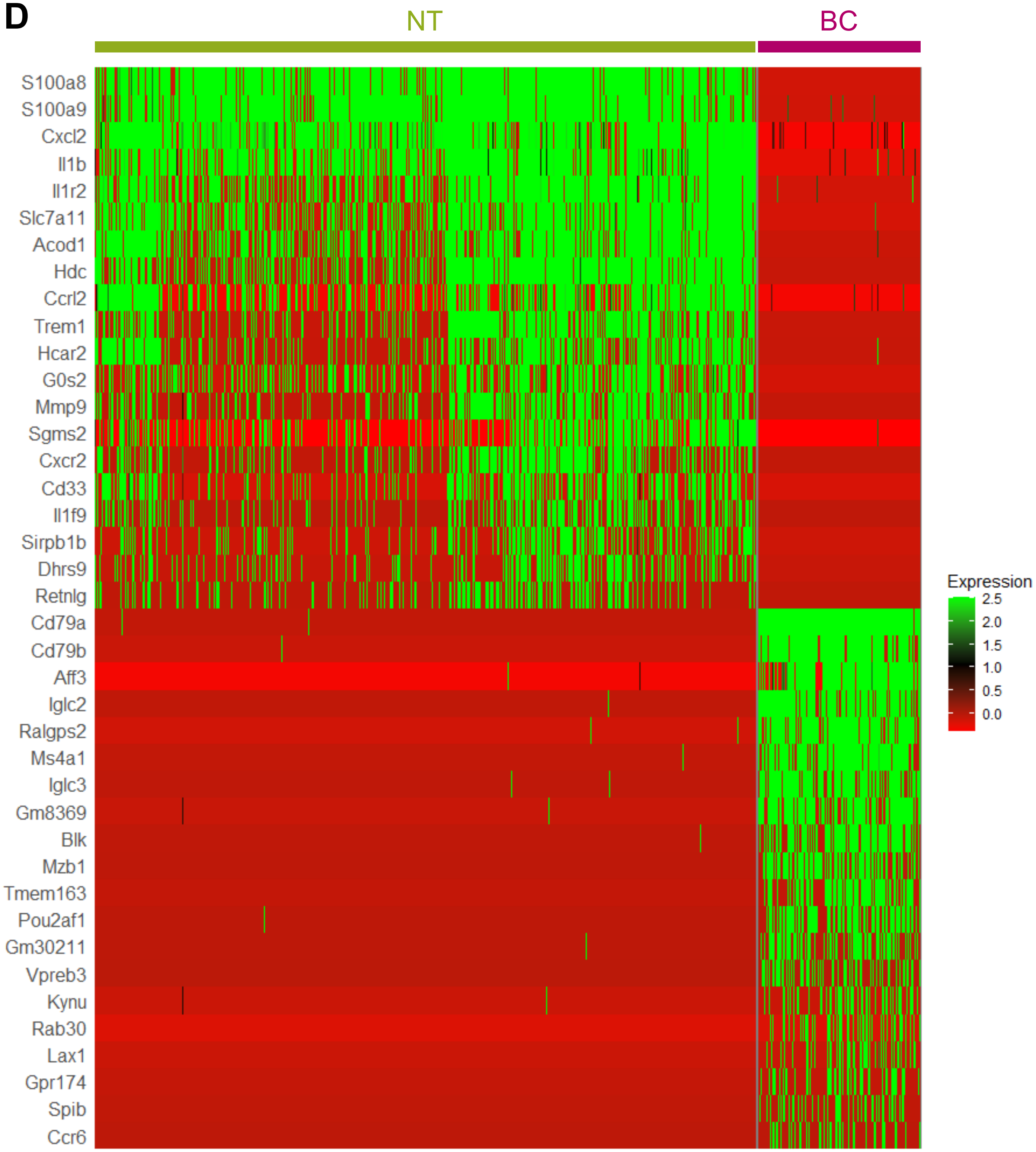

NT
BC
S100a8
S100a9
Cxcl2
Il1b
Il1r2
Slc7a11
Acod1
Hdc
Ccrl2
Trem1
Hcar2
G0s2
Mmp9
Sgms2
Cxcr2
Cd33
Il1f9
Sirpb1b
Dhrs9
Retnlg
Cd79a
Cd79b
Aff3
Iglc2
Ralgps2
Ms4a1
Iglc3
Gm8369
Blk
Mzb1
Tmem163
Pou2af1
Gm30211
Vpreb3
Kynu
Rab30
Lax1
Gpr174
Spib
Ccr6
Expression
2.5
2.0
1.5
1.0
0.5
0.0

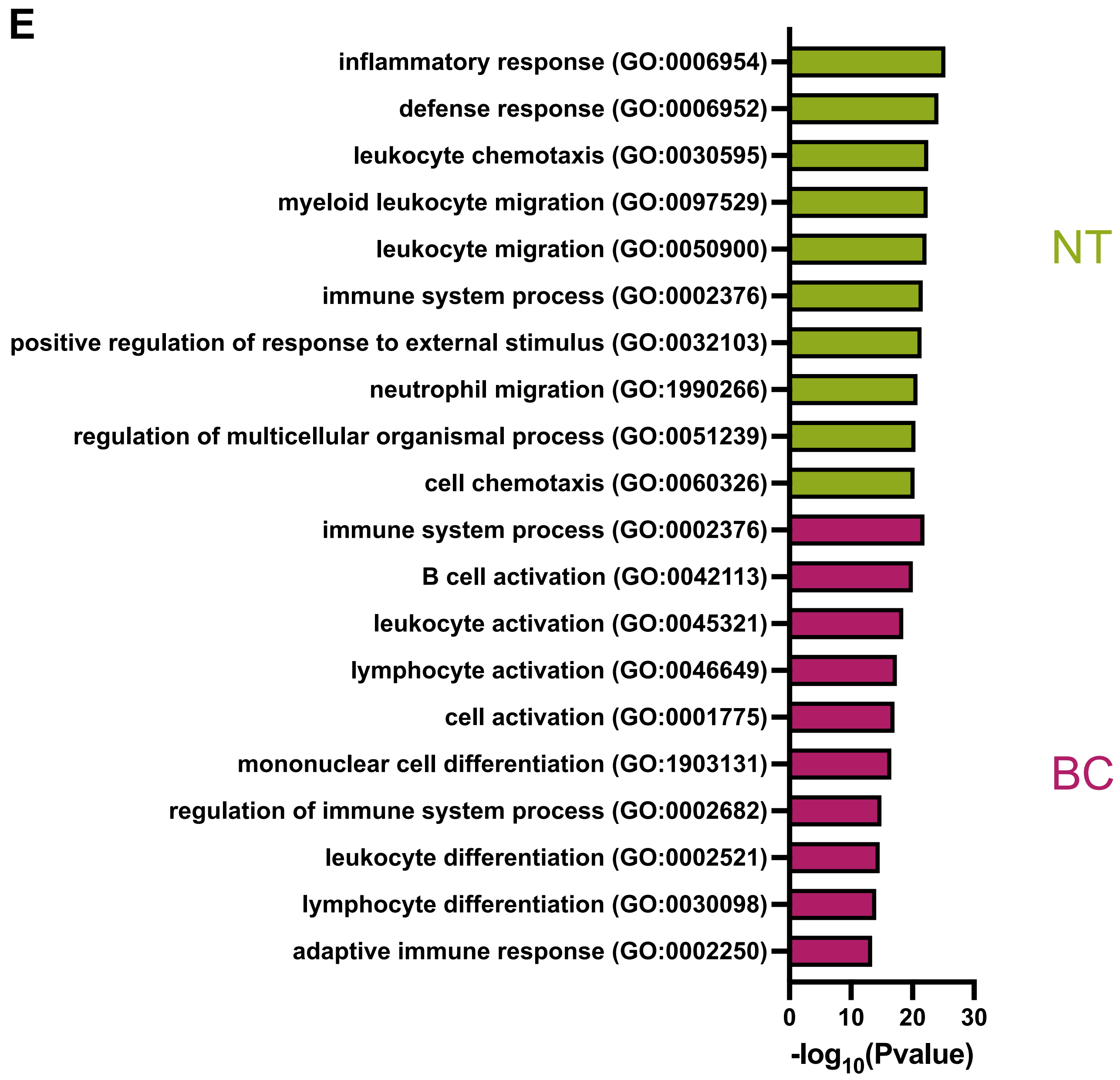


**Supplementary Figure 12. D-flow under hypercholesterolemia increases the neutrophil and B cell accumulation**
**A**. UMAP plot of NT and BC clusters (887 and 217 cells, respectively). **B**. Cell number and % cell population (cell number for each NT and BC cluster normalized by the total number of NTs and BCs per group) quantifications for the NT and BC clusters across all 10 experimental groups. Only the cell numbers are shown. **C**. UMAP plot for each experimental group is shown. N= 5-20 mice for each condition. **D.** Heatmap of top 20 most highly enriched genes for the NT and BC clusters. **E.** Top 10 GO BP terms using top 100 differentially upregulated genes for NT and BC clusters.

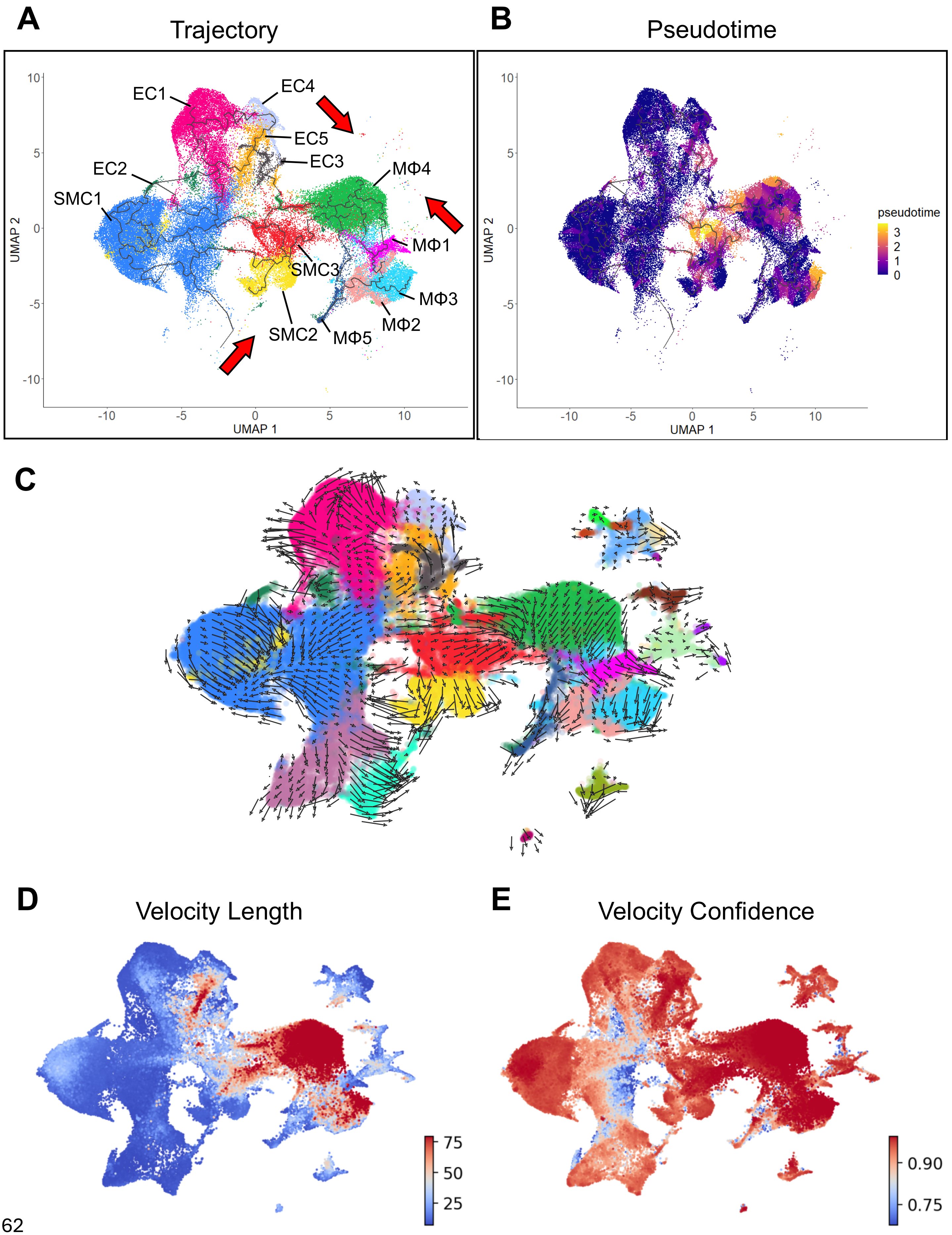
A
Trajectory
EC1
EC4
EC5
EC3
EC2
SMC1
MΦ4
MΦ1
SMC3
MΦ3
MΦ2
SMC2
MΦ5
UMAP 1
UMAP 2
B
Pseudotime
pseudotime
UMAP 1
UMAP 2
C
D
Velocity Length
E
Velocity Confidence

**Supplementary Figure 13. Trajectory analyses show EC-derived foam cells share similar transcriptomic profiles with foam cells derived from SMCs and MΦs**

**A-B**. Monocle 3 trajectory analysis of ECs, SMCs, and MΦs from the mouse scRNA-seq data visualized on the UMAP plot labelled by cell cluster annotation (**A**) and pseudotime (**B**). EC1, SMC1, MΦ1, and MΦ5 were set as roots on the UMAP plot (dark blue = earliest time point, yellow = latest time point). **C**. RNA velocity vector field depicts cell fate transitions of the mouse scRNA-seq data mapped onto the UMAP plot labelled by cell cluster annotation. **D**. Velocity length map of individual cells (red = high rate of cell differentiation, blue = low rate of cell differentiation). **E**. Velocity confidence map of individual cells (red = high confidence, blue = low confidence).

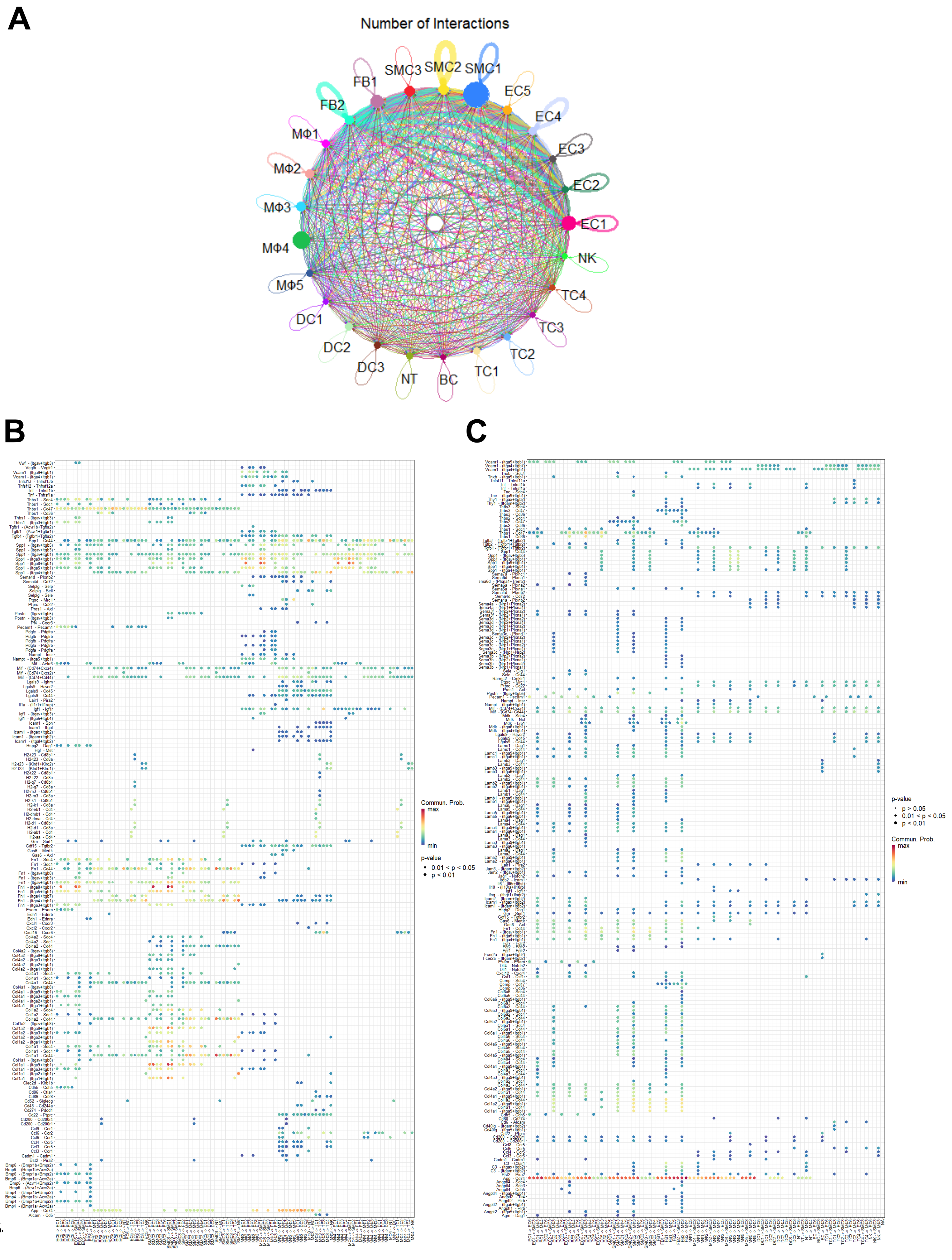
A
Number of Interactions
SMC3
SMC2
SMC1
FB1
EC5
FB2
EC4
MΦ1
EC3
MΦ2
EC2
MΦ3
EC1
MΦ4
NK
MΦ5
TC4
DC1
TC3
DC2
TC2
DC3
TC1
NT
BC
B
C
Commun. Prob.
max
min
p-value
0.01 < p < 0.05
p < 0.01
p-value
p > 0.05
0.01 < p < 0.05
p < 0.01
Commun. Prob.
max
min

**Supplementary Figure 14. *CellChat* cell-cell communication analysis identifies signaling pathways involving immune-like and foam cells derived from ECs, SMCs, and MΦs**

**A**. Circle plot depicts cell-cell communication network of 113 predicted signaling pathways among all cell clusters from the mouse scRNA-seq data. **B-C.** Bubble plots show all significant ligand-receptor interactions (red = maximum communication probability, blue = minimum communication probability) sent **(B)** or received (**C)** by the foam cell clusters (EC5, SMC3, MΦ3, and MΦ4).

1

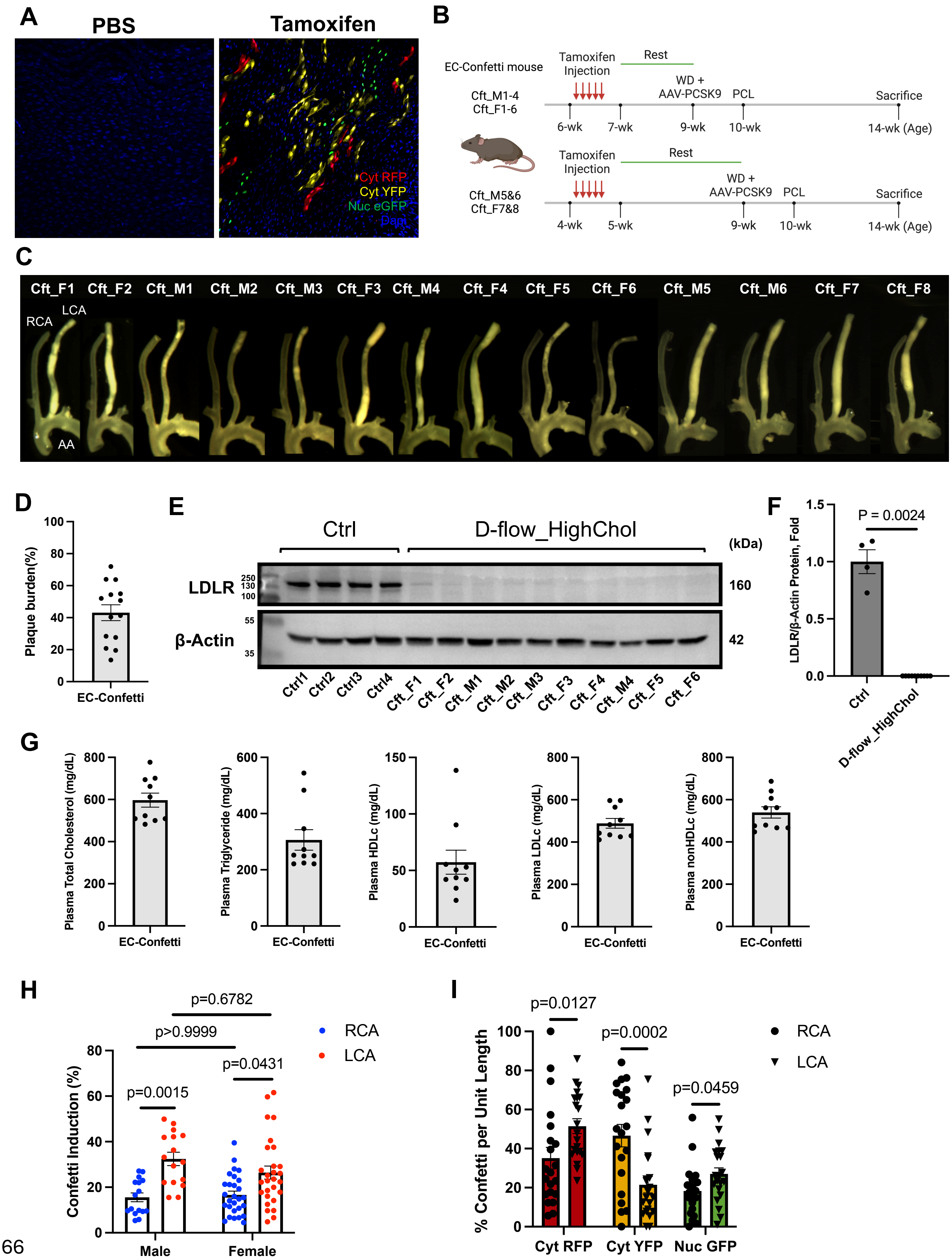
A
PBS
Tamoxifen
Cyt RFP
Cyt YFP
Nuc eGFP
Dapi
B
EC-Confetti mouse
Tamoxifen Injection
Rest
WD + AAV-PCSK9
PCL
Sacrifice
Cft_M1-4
Cft_F1-6
6-wk
7-wk
9-wk
10-wk
14-wk (Age)
Cft_M5&6
Cft_F7&8
4-wk
5-wk
9-wk
10-wk
14-wk (Age)
C
Cft_F1 Cft_F2 Cft_M1 Cft_M2 Cft_M3 Cft_F3 Cft_M4 Cft_F4 Cft_F5 Cft_F6 Cft_M5 Cft_M6 Cft_F7 Cft_F8
RCA
LCA
AA
D
Plaque burden(%)
EC-Confetti
E
Ctrl
D-flow_HighChol
(kDa)
LDLR
β-Actin
160
42
Ctrl1 Ctrl2 Ctrl3 Ctrl4 Cft_F1 Cft_F2 Cft_M1 Cft_M2 Cft_M3 Cft_F3 Cft_F4 Cft_M4 Cft_F5 Cft_F6
F
P = 0.0024
LDLR/β-Actin Protein, Fold
Ctrl
D-flow_HighChol
G
Plasma Total Cholesterol (mg/dL)
Plasma Triglyceride (mg/dL)
Plasma HDLc (mg/dL)
Plasma LDLc (mg/dL)
Plasma nonHDLc (mg/dL)
EC-Confetti
H
p=0.6782
p>0.9999
p=0.0431
p=0.0015
RCA
LCA
Confetti Induction (%)
Male
Female
I
p=0.0127
p=0.0002
p=0.0459
RCA
LCA
% Confetti per Unit Length
Cyt RFP
Cyt YFP
Nuc GFP

66

**Supplementary Figure 15. Timeline for lineage tracing study on EC-Confetti mice and validations of confetti EC induction and atherosclerotic plaque development at 4 wks post-PCL**

**A.** Images of mouse descending aorta confirm the expressions of confetti$^+$ ECs upon tamoxifen induction. **B.** Timeline for EC-confetti mice receiving hypercholesterolemia and d-flow treatments. **C.** Representative gross images (N=14) of LCA, RCA, and aortic arch collected from 21 EC-Confetti mice used for lineage tracing study. **D.** Plaque burden analysis in LCAs of EC-confetti mice (N=14). **E.** Western blot to validate LDLR knockout by AAV8-PCSK9 for the 10 mice used, with β-Actin as the loading control. Ctrl 1-4 are from scRNA-seq experiment at 4 wks. (N=4 for ctrl; N=10 for D-flow_HighCol group). **F.** Quantifications for western blots. **G.** Plasma lipid level analyses of total cholesterol, triglycerides, high-density lipoprotein cholesterol (HDLc), low-density lipoprotein cholesterol (LDLc), and non-HDLc (N=10) in EC-confetti mice. **H.** Confetti induction rate in RCA and LCA, grouped by the gender of mice (N=16 to 28). **I.** Color distribution of Confetti+ ECs in the RCAs and LCAs. Compared to the contralateral RCAs, the LCAs revealed 46.71% and 47.79% increase in RFP+ and GFP+ ECs, respectively, whereas the number of YFP+ ECs reduced by 53.90% under d-flow+hypercholesterolemia. All quantifications are presented as mean ± SEM. P value in western blot analysis was calculated using 2-tailed unpaired Student t test. P values in confetti induction rate were calculated by Kruskal Wallis one-way ANOVA due to nonnormality.

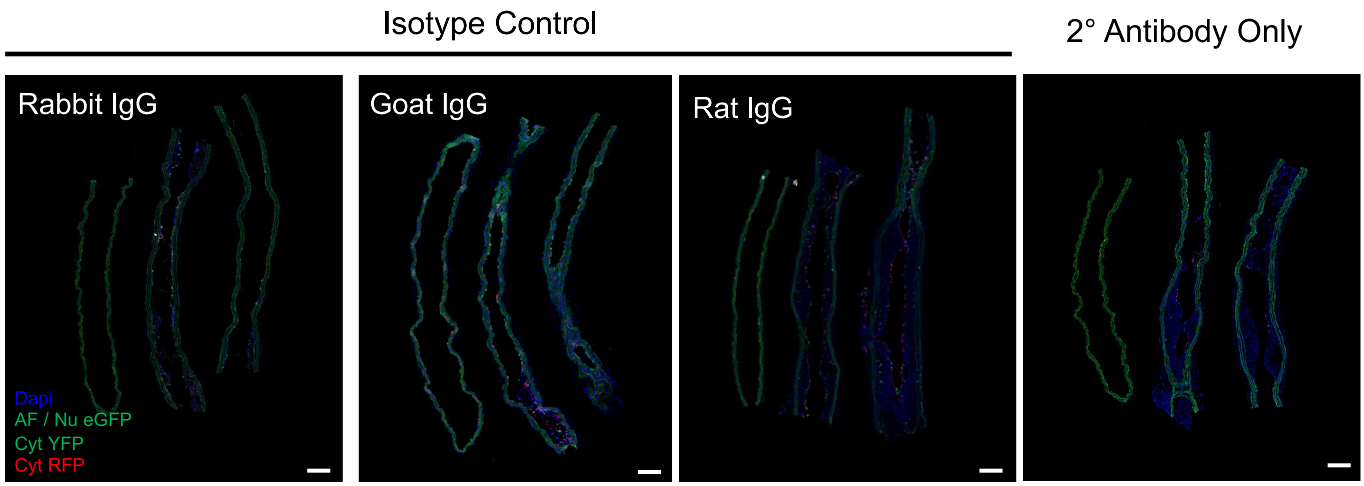


**Supplementary Figure 16. Isotype and secondary-only controls to test the specificity of the primary antibodies.** Images of carotid arteries incubated with rabbit and rat IgG (Thermo, 31903) a day prior to primary antibody addition or secondary antibody alone confirm the absence of non-specific binding of primary antibodies used for all inflammation, EndMT, EndIT, and EndFT markers. Scale bar: 200 µm.

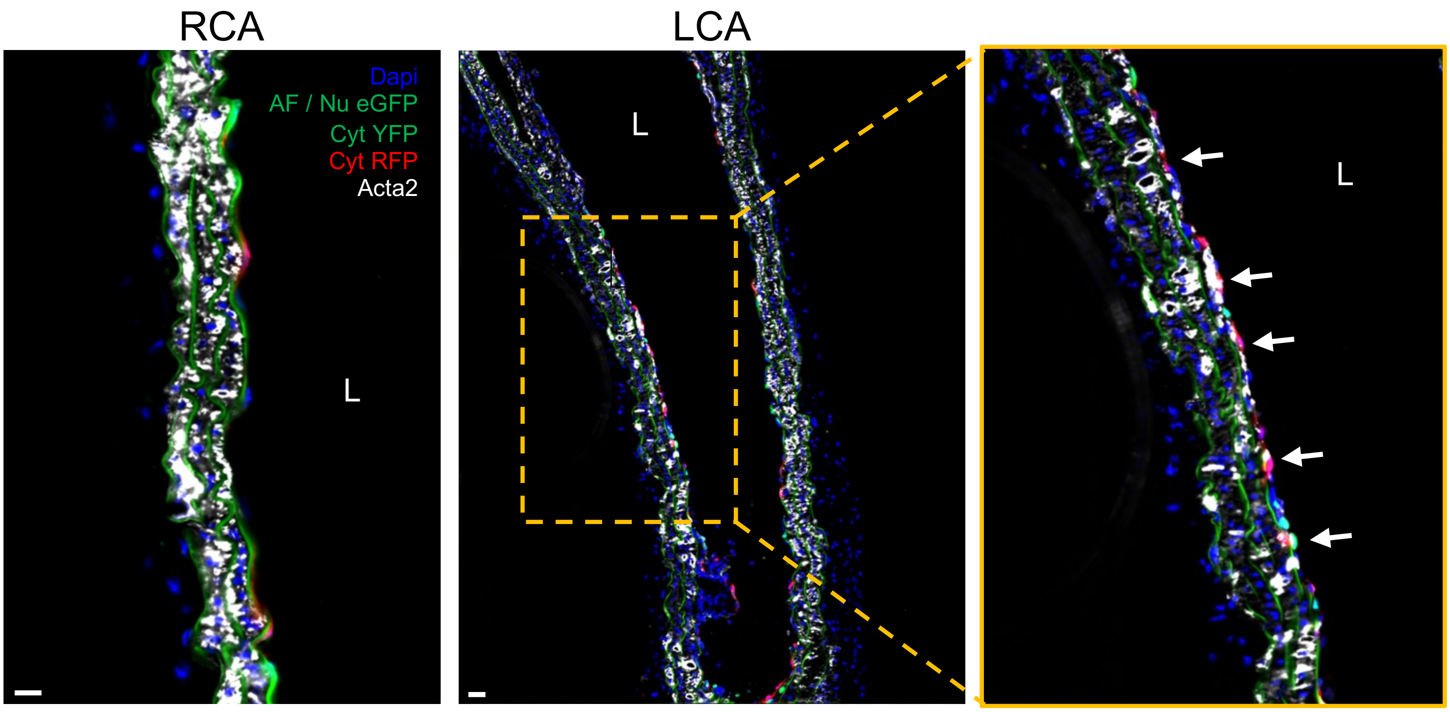


**Supplementary Figure 17. Representative immunofluorescence images of EC-Confetti mice show Acta2 expression in the medial layer of RCA and LCA**

40X widefield immunofluorescence images of RCA and LCA immunostained for EndMT marker Acta2. White arrow denotes $Acta2^{+}$ $Confetti^{+}$ EC. Scale bar: 20 μm. L: Lumen.

1

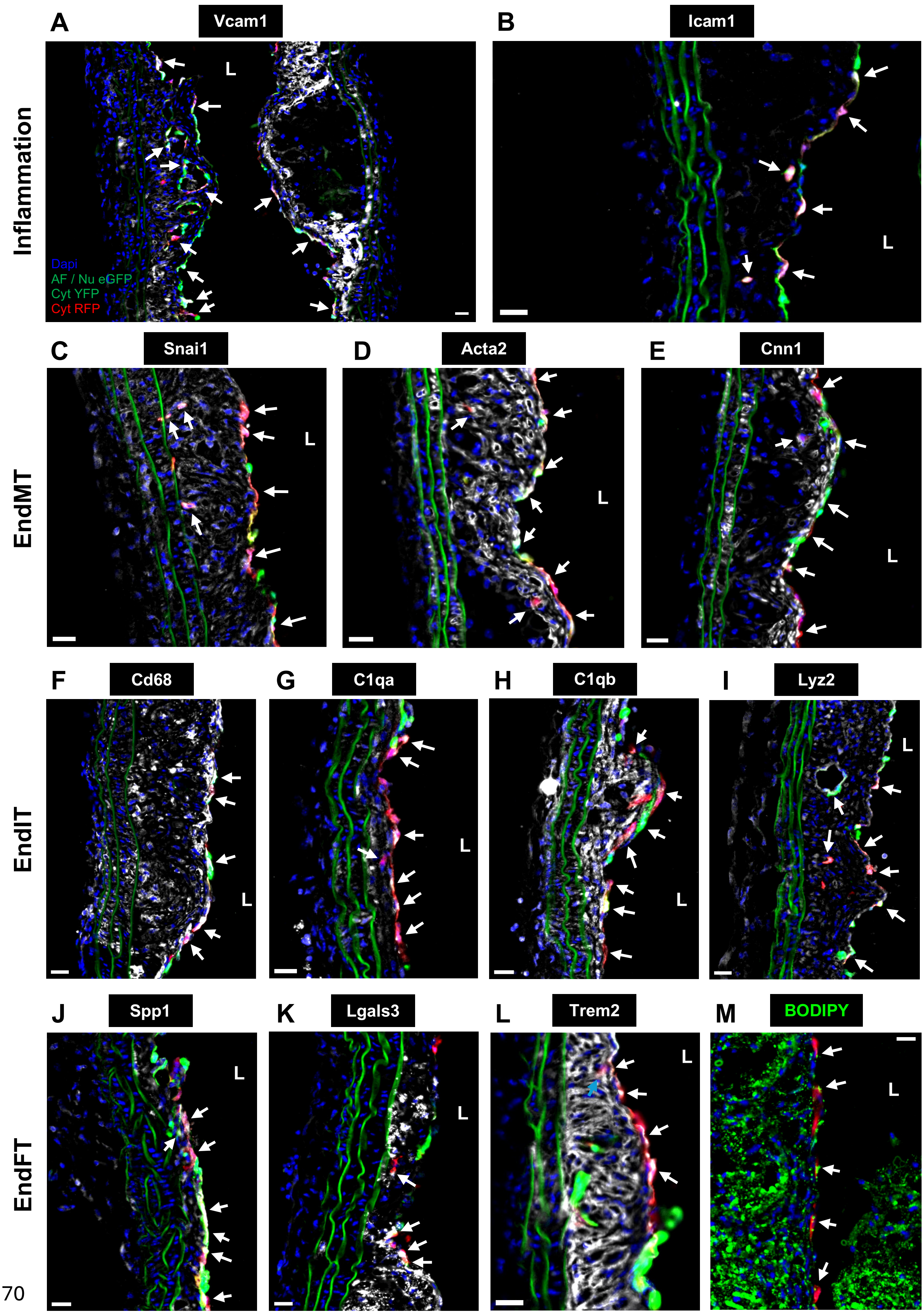
A
Vcam1
B
Icam1
Inflammation
L
Dapi
AF / Nu eGFP
Cyt YFP
Cyt RFP
C
Snai1
D
Acta2
E
Cnn1
EndMT
F
Cd68
G
C1qa
H
C1qb
I
Lyz2
EndIT
J
Spp1
K
Lgals3
L
Trem2
M
BODIPY
EndFT

70

**Supplementary Figure 18. Representative immunofluorescence images of EC-Confetti mice show FIRE (endothelial inflammation, EndMT, EndIT, and EndFT) in the plaque area of LCAs under d-flow and hypercholesterolemia at 4 weeks post-PCL.**
40X widefield immunofluorescence images of RCA and LCA immunostained for endothelial inflammation (Vcam1 and Icam1, **A-B**); EndMT (Snai1, Acta2, and Cnn1, **C-E**); EndIT (Cd68, C1qa, C1qb, and Lyz2, **F-I**); and EndFT (Spp1, Lgals3, Trem2, and BODIPY, **J-L**). White arrow denotes $FIRE^{+}$ $Confetti^{+}$ EC. Scale bar: 20 µm. L: Lumen.

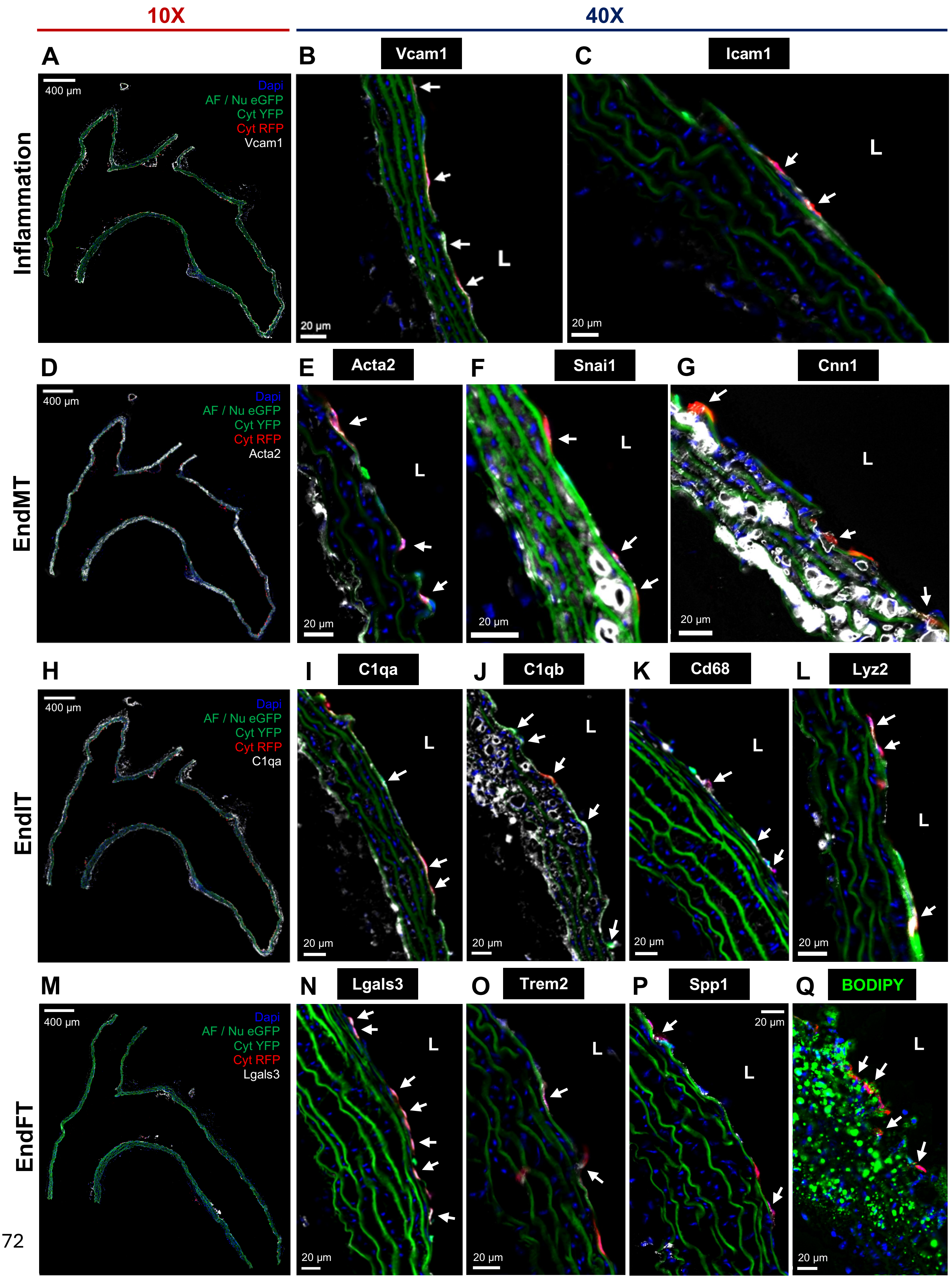
10X
40X
A
B
Vcam1
C
Icam1
Inflammation
400 µm
Dapi
AF / Nu eGFP
Cyt YFP
Cyt RFP
Vcam1
L
20 µm
D
E
Acta2
F
Snai1
G
Cnn1
EndMT
Acta2
H
I
C1qa
J
C1qb
K
Cd68
L
Lyz2
EndIT
C1qa
M
N
Lgals3
O
Trem2
P
Spp1
Q
BODIPY
EndFT
Lgals3

72

1

**Supplementary Figure 19. Representative immunofluorescence images of EC-Confetti mice showing markers of FIRE (endothelial inflammation, EndMT, EndIT, and EndFT) in the aortic arches.** Longitudinal sections of the aortic arches from EC-Confetti mice treated with d-flow and hypercholesterolemia at 4 weeks post-PCL, as described in Figure 6, were imaged by fluorescence microscopy with markers of endothelial inflammation (Vcam1 and Icam1, **A-C**); EndMT (Snai1, Acta2, and Cnn1, **D-G**); EndIT (Cd68, C1qa, C1qb, and Lyz2, **H-L**); and EndFT (Spp1, Lgals3, Trem2, and BODIPY, **M-Q**). **A, D, H,** and **M** show merged images of confetti and FIRE markers at low magnification (10X), while the rest show 40X images. Confetti signals show eGFP (green), YFP (green), and RFP (red). All FIRE markers are shown in white except for green BODIPY (**Q**). White arrows indicate confetti$^{+}$ ECs co-expressing the FIRE markers.

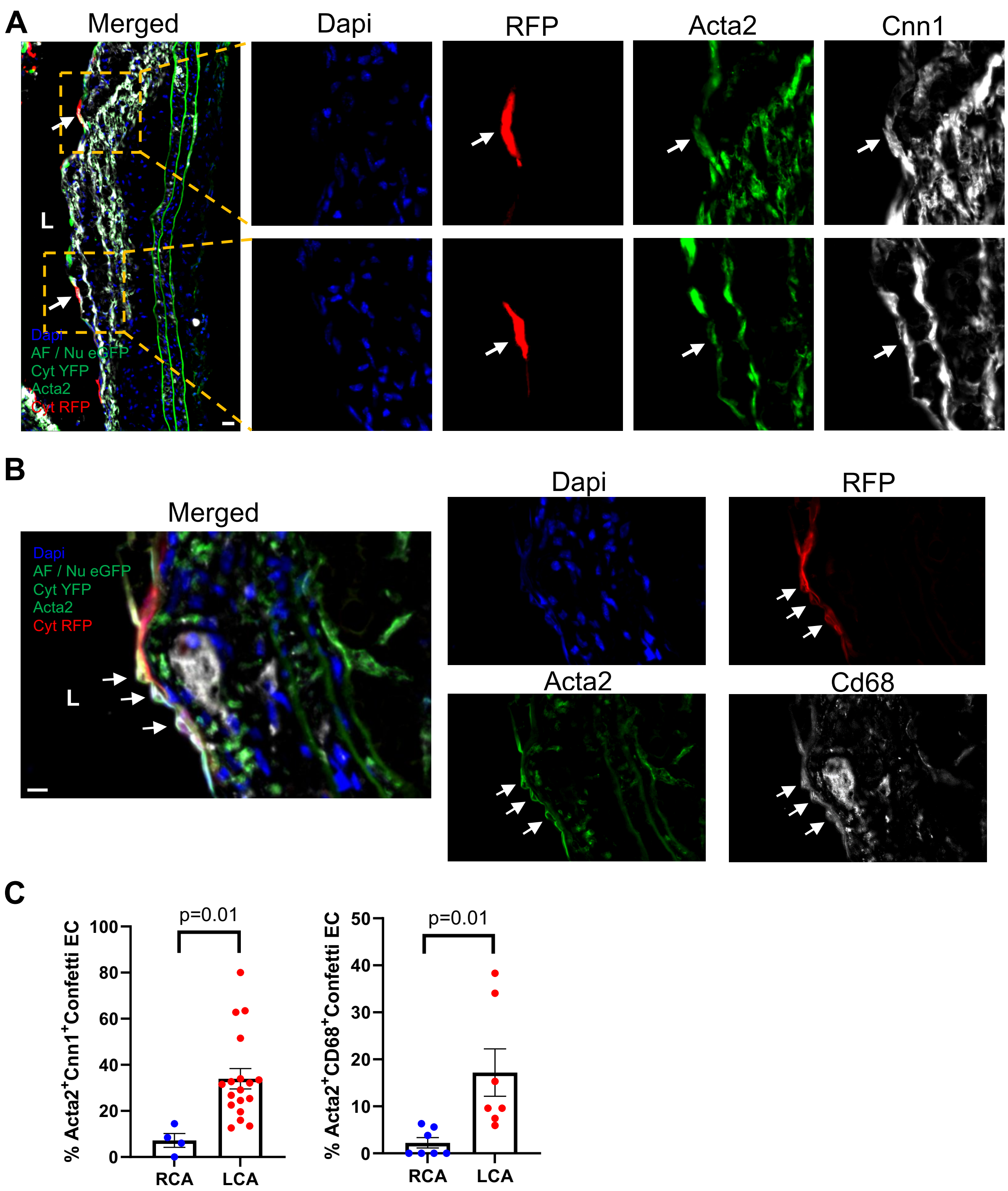


**Supplementary Figure 20. Representative immunofluorescence images of EC-Confetti mice show co-expression of EndMT (Acta2 & Cnn1) and EndMT/EndIT markers (Acta2 & Cd68) in the plaque area of LCAs under d-flow and hypercholesterolemia at 4 weeks post-PCL.** 40X widefield immunofluorescence images of LCA co-expressing EndMT (Acta2, green & Cnn1, white) and EndMT/EndIT markers (Acta2, green & Cd68, white). **A** shows representative images of luminal RFP$^{+}$ EC in the LCA expressing Acta2 and Cnn1. White arrow denotes EndMT$^{+}$ RFP$^{+}$ EC. Scale bar: 20 µm. L: Lumen. **B** shows representative images of luminal RFP$^{+}$ EC in the LCA expressing both Acta2 and Cd68. White arrow denotes EndMT$^{+}$ EndIT$^{+}$ RFP$^{+}$ EC. **C** shows quantifications comparing the % Acta2+/Cnn1+/RFP+ ECs (left) and % Acta2+/Cd68+/RFP+ ECs (right) in the RCAs vs LCAs. Scale bar: 20 µm. L: Lumen.

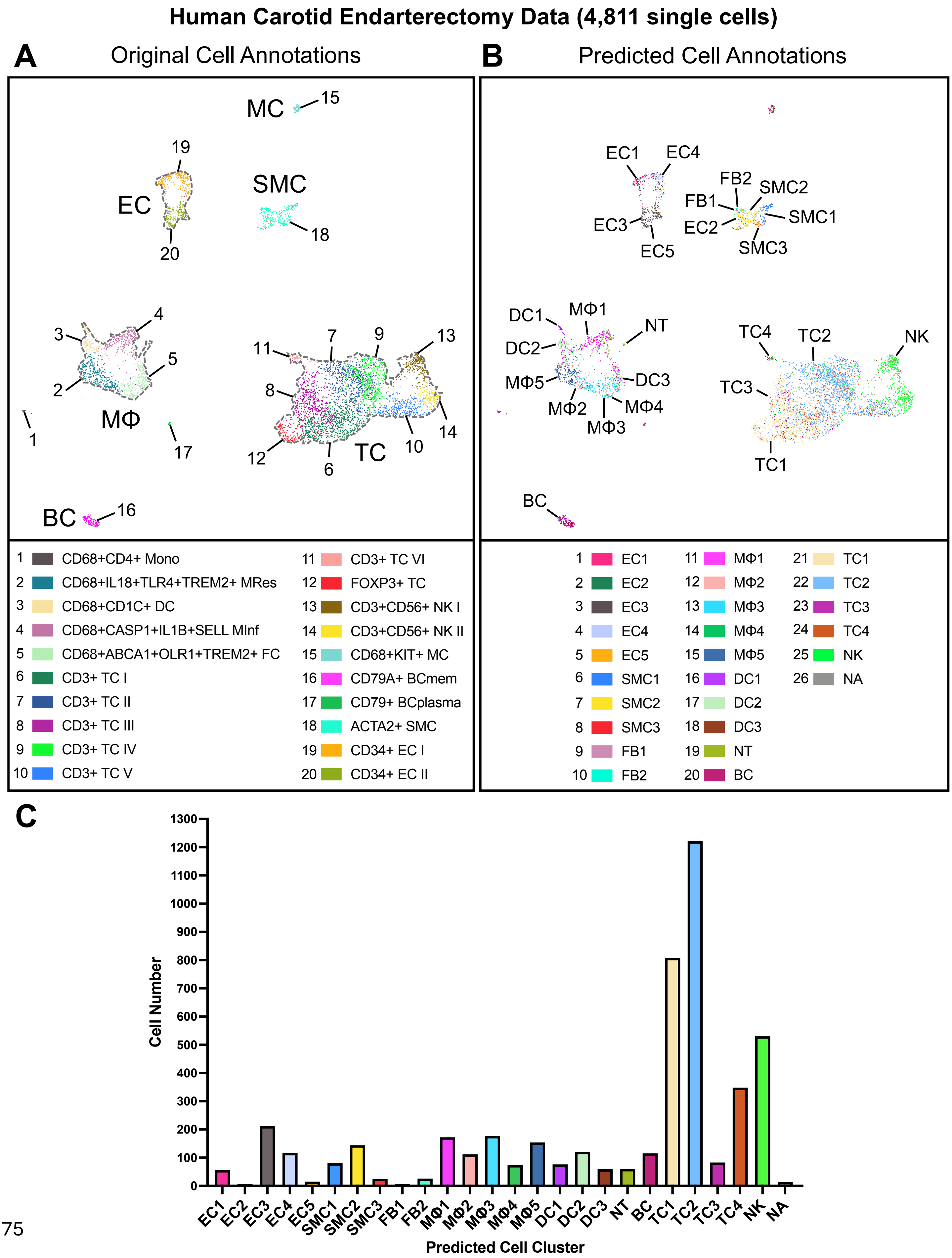

Human Carotid Endarterectomy Data (4,811 single cells)
A
Original Cell Annotations
MC
15
19
EC
SMC
18
20
4
3
5
2
MΦ
1
17
11
7
9
13
8
12
6
TC
10
14
BC
16
1 CD68+CD4+ Mono
2 CD68+IL18+TLR4+TREM2+ MRes
3 CD68+CD1C+ DC
4 CD68+CASP1+IL1B+SELL MInf
5 CD68+ABCA1+OLR1+TREM2+ FC
6 CD3+ TC I
7 CD3+ TC II
8 CD3+ TC III
9 CD3+ TC IV
10 CD3+ TC V
11 CD3+ TC VI
12 FOXP3+ TC
13 CD3+CD56+ NK I
14 CD3+CD56+ NK II
15 CD68+KIT+ MC
16 CD79A+ BCmem
17 CD79+ BCplasma
18 ACTA2+ SMC
19 CD34+ EC I
20 CD34+ EC II
B
Predicted Cell Annotations
EC1
EC4
FB2
SMC2
FB1
SMC1
EC3
EC2
EC5
SMC3
DC1
MΦ1
NT
DC2
MΦ5
DC3
MΦ4
MΦ2
MΦ3
TC4
TC2
NK
TC3
TC1
BC
1 EC1
2 EC2
3 EC3
4 EC4
5 EC5
6 SMC1
7 SMC2
8 SMC3
9 FB1
10 FB2
11 MΦ1
12 MΦ2
13 MΦ3
14 MΦ4
15 MΦ5
16 DC1
17 DC2
18 DC3
19 NT
20 BC
21 TC1
22 TC2
23 TC3
24 TC4
25 NK
26 NA
C
Cell Number
1300
1200
1100
1000
900
800
700
600
500
400
300
200
100
0
EC1
EC2
EC3
EC4
EC5
SMC1
SMC2
SMC3
FB1
FB2
MΦ1
MΦ2
MΦ3
MΦ4
MΦ5
DC1
DC2
DC3
NT
BC
TC1
TC2
TC3
TC4
NK
NA
Predicted Cell Cluster

**Supplementary Figure 21. Reanalysis of publicly available human scRNA-seq data of carotid plaques shows evidence of potential foam cells derived from MΦs, SMCs, and ECs**

**A**. UMAP plot of 4,811 single cells from 44 human carotid endarterectomy samples from a publicly available scRNA-seq data with 20 unique cell clusters is shown. **B**. UMAP plot of the human carotid scRNA-seq data shown in **A** with predicted cell annotations from reference map analysis using our scRNA-seq data as a reference. Foam cell clusters (EC5, SMC3, and MΦ3/4) are highlighted in yellow. **C**. Quantification of cell number for each predicted cell cluster from reference map analysis of human carotid scRAN-seq data shown in **B**.

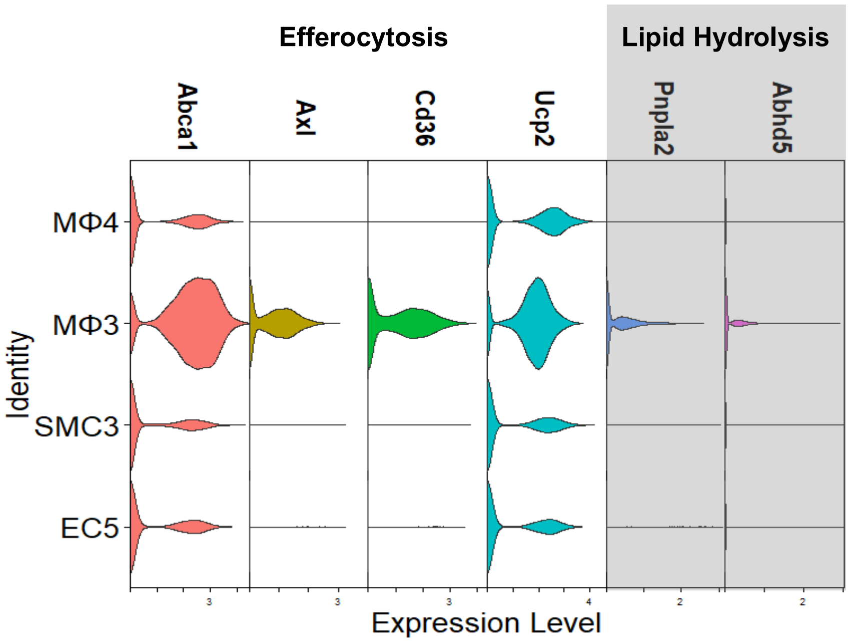


**Supplementary Figure 22. Foam cells developed by d-flow under hypercholesterolemic condition for 4 weeks lose expression of markers for efferocytosis and lipid hydrolysis**
Stacked violin plot showing expression levels of markers of efferocytosis (*Abca1*, *Axl*, *Cd36*, and *Ucp2*) and lipid droplet hydrolysis (*Pnpla2*, or *Atgl*, and *Abhd5*, or *Cgi58*) for foam cell clusters (EC5, SMC3, MΦ3, and MΦ4).
